# Proteome allocation of the microbiome reveals how diet and metabolic dysbiosis impact disease

**DOI:** 10.1101/2024.11.21.624724

**Authors:** Juan D. Tibocha-Bonilla, Rodrigo Santibáñez-Palominos, Yuhan Weng, Manish Kumar, Karsten Zengler

## Abstract

The gut microbiome plays a critical role in human health, spurring extensive research into host-microbe interactions using multi-omic technologies. Although these tools offer valuable insights, they often fall short in capturing the complexity of microbial interactions and assigning causality to disease onset, progression, and treatment. Proteome allocation is intertwined with microbial interactions, yet there is little mechanistic knowledge. Here, we deploy models of metabolism and gene expression (ME-models) that predict proteome allocation. Despite their potential, their deployment has been limited by the laborious reconstruction process. To address this, we reconstructed 495 ME-models for human gut microbes using an automated pipeline. By integrating ME-models with multi-omics data from patients with inflammatory bowel disease (IBD), we identified taxa responsible for variations in amino acids, short-chain fatty acids, and pH in the gut of IBD patients. Thus, this approach provides a mechanistic understanding and paves the way for new intervention strategies.

## Introduction

The gut microbiome plays a central role in human health,^1^ and imbalances in the microbiome are strongly associated with diseases.^2–4^ Understanding microbial composition and metabolic activity provides insights into host-microbe interactions that drive disease onset and progression^5^. However, these interactions are complex^6^ and require intensive experimental approaches combined with computational models and bioinformatics tools.^6^ While metagenomics ^2,7^ and metatranscriptomics^6,7^ can assess the dynamic abundance and infer the possible metabolic activity of microorganisms, integrating these data to unravel the mechanisms^8,9^ driving host-microbe interactions remains challenging^10^. Thus, new approaches for multi-omic data integration and causality elucidation are needed. Here, we propose the use of genome-scale metabolic and expression models to bridge omics data and define causality for microbiome activity related to disease.

Genome-scale metabolic models (M-models) have been at the forefront of predicting microbial growth, and crosstalk^6,11^ They are powerful tools for a wide range of biotechnological and medical applications.^12^ However, M-models have limitations that restrict their use and conclusions. For example, M-models neglect the condition-specific biosynthetic cost of enzymes and pathways^13^ (Figure 1A), leading to unrealistic growth rate predictions^14^ and metabolic flux variability.^15^ Additionally, the fixed biomass composition in M-models requires extensive manual corrections to reflect changes due to environmental perturbations.^16,17^ In dynamic environments such as the human gut, vast experimental data would therefore be required to increase prediction accuracy. Obtaining this kind of data at high phylogenetic resolution is challenging for most bacteria. Moreover, data generated from bacteria in isolation rarely represents bacterial metabolism *in situ*.

**Figure 1.**
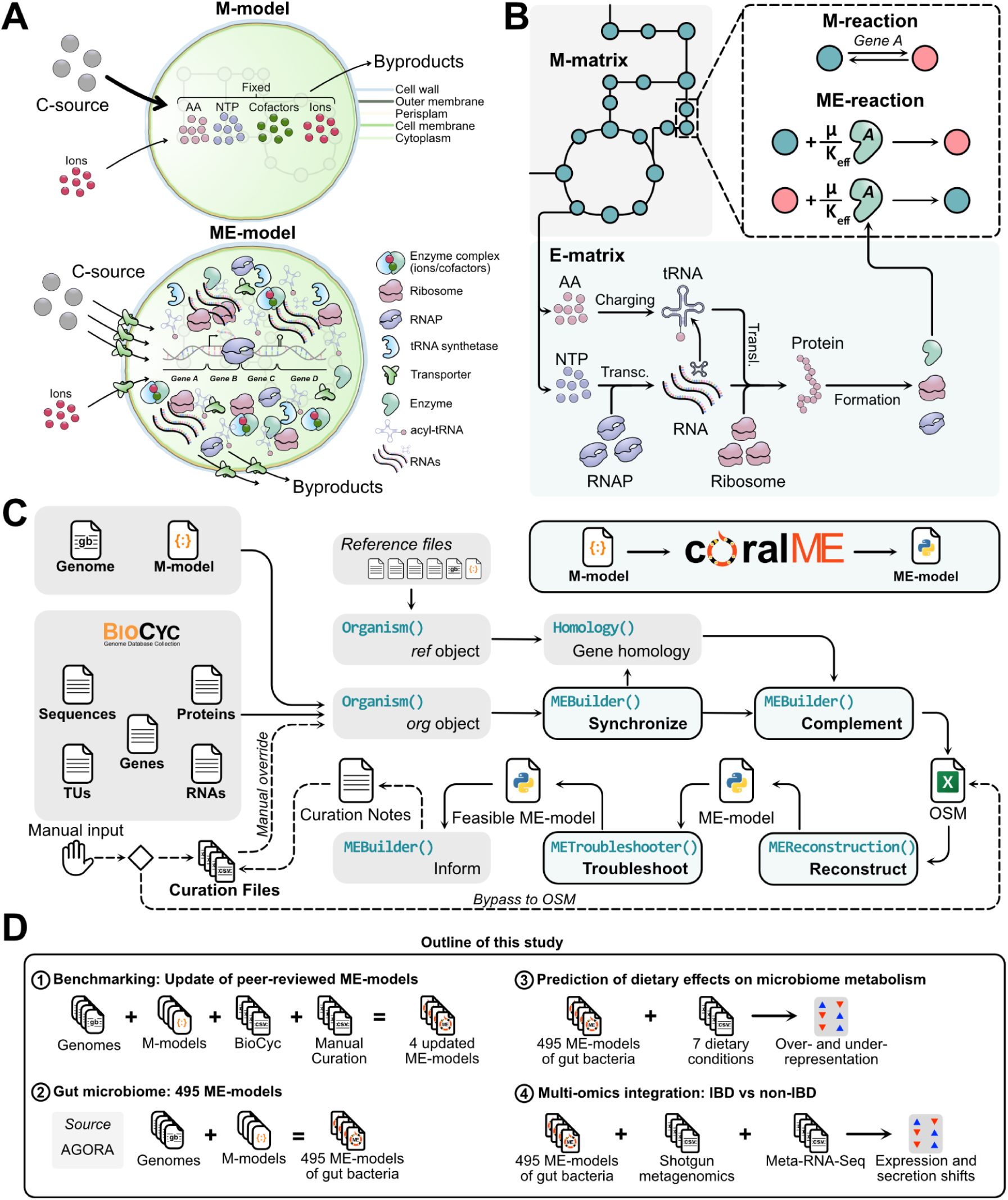
Workflow to generate and utilize ME-models. **(A)** Biomass composition in M-model and ME-model, the latter reflecting condition-specific resource requirements that the ME-model computes through optimization, rather than being fixed in the biomass objective function. **(B)** Network visualization of the interplay between metabolic (M-matrix) and expression (E-matrix) networks. The M-matrix provides precursors and energy used for gene expression, which in turn provides the necessary enzymes to catalyze M- and E-matrix reactions. Enzymes are accounted for in the metabolic reactions coupling coefficients dependent on the growth rate (*μ*) and the effective turnover rate (*K_eff_*).^13^ **(C)** The coralME flow of information. Transcription unit (TU) composition, gene sequences, and gene annotation are provided by four files: *Genome* in GenBank format with the nucleotide sequence (mandatory), and the *TUs, Genes,* and the *Sequences* file in FASTA format. The *Genome* file and the optional *RNAs* file specify the type and sequence of RNAs to simulate. Next, GPRs in the *M-model* file (mandatory as either SBML or JSON format) and the optional *Proteins* file define complex compositions and locations. Additionally, the *M-model* file provides the metabolic network and its associations with enzymatic complexes. **(D)** Sequential applications of coralME performed in this study. First, four peer-reviewed ME-models were updated to prove correctness and effectiveness of the pipeline. Once the pipeline was benchmarked, we deployed it to reconstruct 495 ME-models of gut bacteria and predicted over- and under-representation of bacterial genera in seven dietary conditions. Lastly, we integrated multi-omics data of an inflammatory bowel disease (IBD) study to elucidate metabolic, expression, and secretion shifts associated with the disease.

Recognizing these shortcomings, next-generation models of metabolism and gene expression (ME-models) have been introduced. ME-models simulate condition-specific biosynthetic costs by coupling substrates with enzymes and metabolic fluxes with gene transcription, translation, translocation, and macromolecular assembly^13,14,18^ (Figure 1, A and B). Consequently, flux variability in ME-models is drastically reduced,^18^ as fewer pathways are equally optimal. ME-models also optimize for parsimonious strategies under nutrient-depleted conditions^14,18^ and predict activation of fermentation and overflow pathways under nutrient excess due to proteome limitation.^19,20^ These self-limiting properties yield consistently biologically relevant results,^13,14^ equipping ME-models to predict metabolic exchanges in the human gut for compounds profoundly impacting the host.^21^ Despite their improved accuracy and scope, ME-models are currently not widely used in microbiome research due to the lack of tools aiding their development and the time-intensive process of reconstructing and curating these nonlinear models (Figure S1).^13,22,23^

Here, we developed an automated pipeline for high-throughput ME-model reconstruction of bacteria (Figure 1C) and leveraged hundreds of ME-models to describe dietary effects and multi-omics-aided predictions of secretory profiles in the human gut microbiome. We first benchmarked our COmprehensive Reconstruction ALgorithm for ME-models (coralME) by updating peer-reviewed ME-models^13,18–20^ and reconstructing ME-models from 17 manually curated M-models, assessing completeness with varying degrees of manual curation (Figure 1D). Then, we reconstructed 495 ME-models for the human gut microbiome from the AGORA resource^9,24^ using a single desktop computer (Figure 1D). We simulated the effect of seven diets on the human gut microbiome, demonstrating the impact of iron and zinc availability and the quality of available carbon sources on bacterial growth and overflow metabolism under anaerobic and microaerobic conditions. Next, we integrated metagenomics and metatranscriptomics data from an inflammatory bowel disease (IBD) study^8^ to elucidate major shifts in metabolic activity and unravel complex secretion profiles of the microbiome (Figure 1D). We identified drivers of global changes in gut pH and levels of ammonium, amino acids, and short chain fatty acids (SCFAs), among others. Together, our findings demonstrate that high-throughput reconstruction of ME-models enables in-depth analysis previously infeasible, providing a detailed mechanistic understanding of the microbiome and opening the door for new intervention and treatment strategies.

## Results

### Reconstructing mechanistic models for the gut microbiome

We automated the reconstruction of functional ME-models (Figure 1C) with minimal user setup (Figure S2) for diverse bacteria, such as representatives of the human gut microbiome. As minimal input, coralME requires an M-model and a compatible GenBank file^25^ to reconstruct a ME-model. Optionally, it can use additional information from BioCyc^26^. Reconstruction of ME-models occurs in four steps: i) data reading and synchronization, ii) data complementation, iii) reconstruction, and iv) troubleshooting (Figure 1C). For more details on coralME development, architecture, and inputs, see STAR Methods and Supplementary Text 1.

### Benchmarking of coralME against peer-reviewed ME-models

We benchmarked coralME against all available peer-reviewed ME-models to ensure the quality of automatically reconstructed ME-models. First, we reconstructed a new ME-model for *Escherichia coli* using coralME, named *i*Eco1689-ME, by updating the previous reconstruction *i*JL1678b-ME.^13^ The reconstruction used updated and curated input files from the original publication.^13^ Due to non-equivalent changes introduced in the code and correction of the input data, *i*Eco1689-ME shows minor discrepancies compared to *i*JL1678b-ME^13^ (Supplementary Tables 1 and 2). We performed an in-depth characterization of the reactions and fluxes between these two ME-models. Reaction fluxes showed an approximately one-to-one linear correlation (log10-transformed, slope ≈ 0.996, R^2^ ≈ 0.998, Figure 2A). Most of the fluxes are distributed similarly (Figure 2B). The differences in the absolute value of fluxes were no higher than 0.76 mmol/gDW/h, and >96% of the active reactions showed differences below 0.01 mmol/gDW/h. Furthermore, the majority of fluxes (93%) have a log_2_ fold change (FC) in the [–1, 0] range (Figure 2C), confirming the high similarity between the two ME-models. Finally, due to the predicted expression of eight new genes in *i*Eco1689-ME, the energy required for translation is minimally higher (0.08 mmol ATP/gDW and 0.14 mmol GTP/gDW) than in *i*JL1678b-ME (Figure S3).

**Figure 2.**
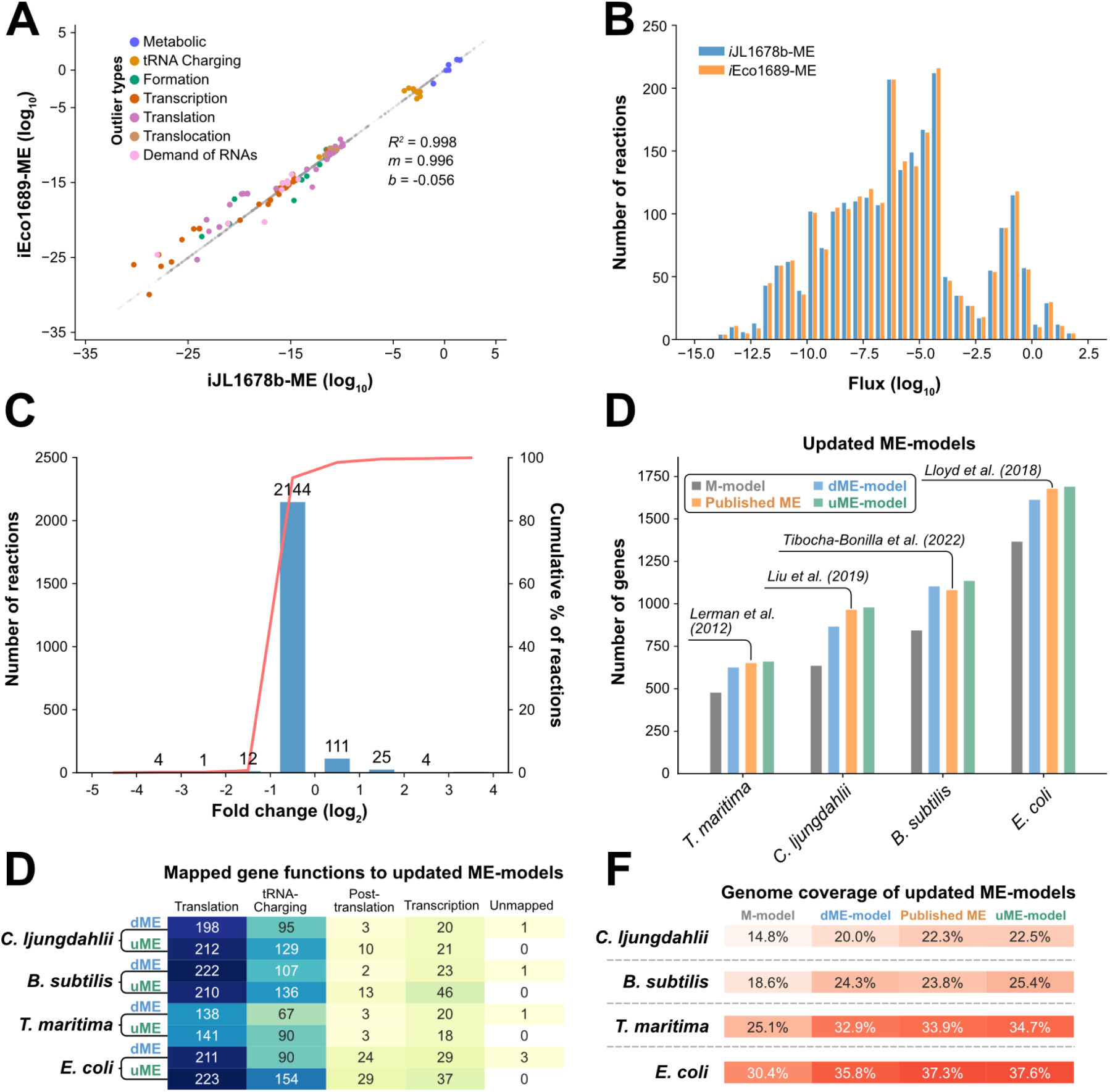
Reconstruction and benchmarking of coralME against peer-reviewed ME-models. The coralME software reconstructed a ME-model using updated information of gene products, reactions, reaction stoichiometries and subcellular location (more information in Supplementary Tables 1 and 2). **(A)** Linear correlation of paired, non-zero fluxes after optimization and log_10_ transformation, comparing *i*Eco1689-ME to *i*JL1678b-ME. The slope (*m*, standard error 0.001), intercept (*b*, error 0.016), and coefficient of determination (R^2^) are shown. 130 fluxes were identified as outliers and highlighted (log_10_ difference ≥ 0.01). **(B)** Distribution of non-zero fluxes [log_10_(abs(value)) transformed] after optimization of both ME-models. **(C)** Distribution of fold-change ratio coralME/COBRAme [log_2_(value) transformed] for the non-zero fluxes after optimization of both ME-models. Cumulative distribution is shown as a red line. **(D)** Comparison of mapped gene functions in M-models, drafts (dME-models), published ME-models, and updated ME-models (uME-models) of *B. subtilis*, *C. ljungdahlii*, *T. maritima*, and *E. coli*. **(E)** Comparison of genome coverage in M-, dME-, published ME-, and the uME-models of *B. subtilis*, *C. ljungdahlii*, *T. maritima*, and *E. coli*, sorted by genome coverage in the M-models. **(F)** Comparison of the number of genes in M-, dME-, published ME-, and uME-models (gray, blue, orange, and green, respectively) of *B. subtilis*, *C. ljungdahlii*, *T. maritima*, and *E. coli*, sorted by gene number in the M-models.

After verifying the correctness of *i*Eco1689-ME reconstructed using coralME, we updated existing ME-models for *Bacillus subtilis,*^19^ *Clostridium ljungdahlii,*^20^ and *Thermotoga maritima.*^18^ We generated draft ME-models (hereinafter dME-models) and incorporated information from the original publications, generating updated ME-models (hereinafter uME-models). Interestingly, 80% to 90% of gene functions were automatically mapped in the uME-models (Figure 2D). The remaining 10% to 20% of missing gene functions (difference between uME- and dME-models in Figure 2E) largely correspond to tRNA and rRNA modifying enzymes, which are distributed across the translation and tRNA charging pathways. Next, manual curation was performed using the generated Curation Notes (Figure 1A) and the information available from the respective studies, resulting in the uME-models *i*Bsu1154-ME (*B. subtilis*), *i*Clj972-ME (*C. ljungdahlii*), and *i*Tma666-ME (*T. maritima*). Notably, genome coverage in dME-models (genes in the ME-model/total genes in the genome) was at most 1.5% less compared to their uME-models (Figure 2F). The genome coverage of these three uME-models and the *i*Eco1689-ME exceeded that of the published ME-models by 0.2% to 1.5%, with a total of 1153, 972, 666, and 1689 genes for the uME-models of *B. subtilis*, *C. ljungdahlii, T. maritima*, and *E. coli*, respectively (Figure 2D).

To further ensure coralME’s capability of reconstructing ME-models from diverse bacteria, we reconstructed 17 dME-models from peer-reviewed M-models of eight different phyla (Figure S4). A full description of these reconstructions is provided in Supplementary Text 2. Hundreds of expression machinery genes were mapped to their function in the dME-models (Figure S5, A and B). Genome coverage of the dME-models was 3.5% to 22.9% higher than M-models, totaling 23.9% to 54.2% (Figure S5C).

### Proteome allocation drives the metabolic responses to dietary changes

Next, we investigated metabolic profiles of the complex gut microbiota using ME-models. We reconstructed ME-models for 495 gut bacteria, increasing the number of genes on average by 19.2% over M-models (Figure S6). We simulated these ME-models under six different conditions, in addition to a “Western Diet” (WD) as a reference.^9,24^ The chosen conditions are linked to gut dysbiosis, namely iron deficiency^27,28^ (Fe-), zinc deficiency^29^ (Zn-), and low oxygen availability (O2+), which occurs close to gut epithelial cells.^30^ We also monitored dietary perturbations, such as a higher intake of carbohydrates (CH+),^31,32^ proteins (Pr+),^33^ or lipids (FA+).^34^

Simulations show significant variations in the effect on the growth rate of the 495 bacteria (Figure 3, A and B). We then compared the growth rate variations under all conditions to the reference (WD) (Figure 3C). Limiting metal ion intake decreased due to their role as enzymatic cofactors (fold changes below 1.0), while a higher intake of organic carbon and oxygen increased growth rates (fold changes above 1.0). The distribution of growth rates revealed differential effects on microorganisms, so we performed Mann-Whitney U-tests to identify significant over- (Figure 3D) or underrepresentation (Figure 3E) of bacterial genera under the six conditions compared to WD. Under iron deficiency, a known contributor to gut dysbiosis,^27,28^ the genera *Alistipes* (p *≤* 0.03)*, Bacteroides* (p *≤* 0.02)*, Bifidobacterium* (p *≤* 0.02)*, Fusobacterium* (p *≤* 10^-4^)*, Helicobacter* (p *≤* 0.02)*, Neisseria* (p *≤* 0.004), and *Porphyromonas* (p *≤* 0.02) were significantly overrepresented (Figure 3D), while the genera *Lactobacillus* (p *≤* 0.01) and *Enterobacter* (p *≤* 0.05) were underrepresented, the latter consistent with prior literature.^35^ No genus was significantly underrepresented in zinc deficiency, but 29 were overrepresented (Figure 3D). As opposed to iron that serves as a cofactor in numerous enzymatic complexes, zinc is principally used during transcription as a cofactor in the β′ subunit of the RNA polymerase.^36^ This results in an overall decrease in growth rates under Zn deficiency (Figure 3, B and C), implying that overrepresented genera were less affected (Figure 3D). Overrepresentation was predicted for *Bacteroides* (*p ≤* 10^-17^), *Bifidobacterium* (*p ≤* 10^-17^), *Streptococcus* (*p ≤* 10^-12^), *Fusobacterium* (*p ≤* 10^-8^), *Campylobacter* (*p ≤* 10^-7^), and *Staphylococcus* (*p ≤* 10^-7^). In clear contrast, no significant over or underrepresentation in Fe- and Zn- diets was predicted by M-models (Figure S7); since these factors are fixed in the biomass reaction, M-models predict a growth rate decrease directly proportional to the reduced rate of iron or zinc uptake (Figure S8).

**Figure 3.**
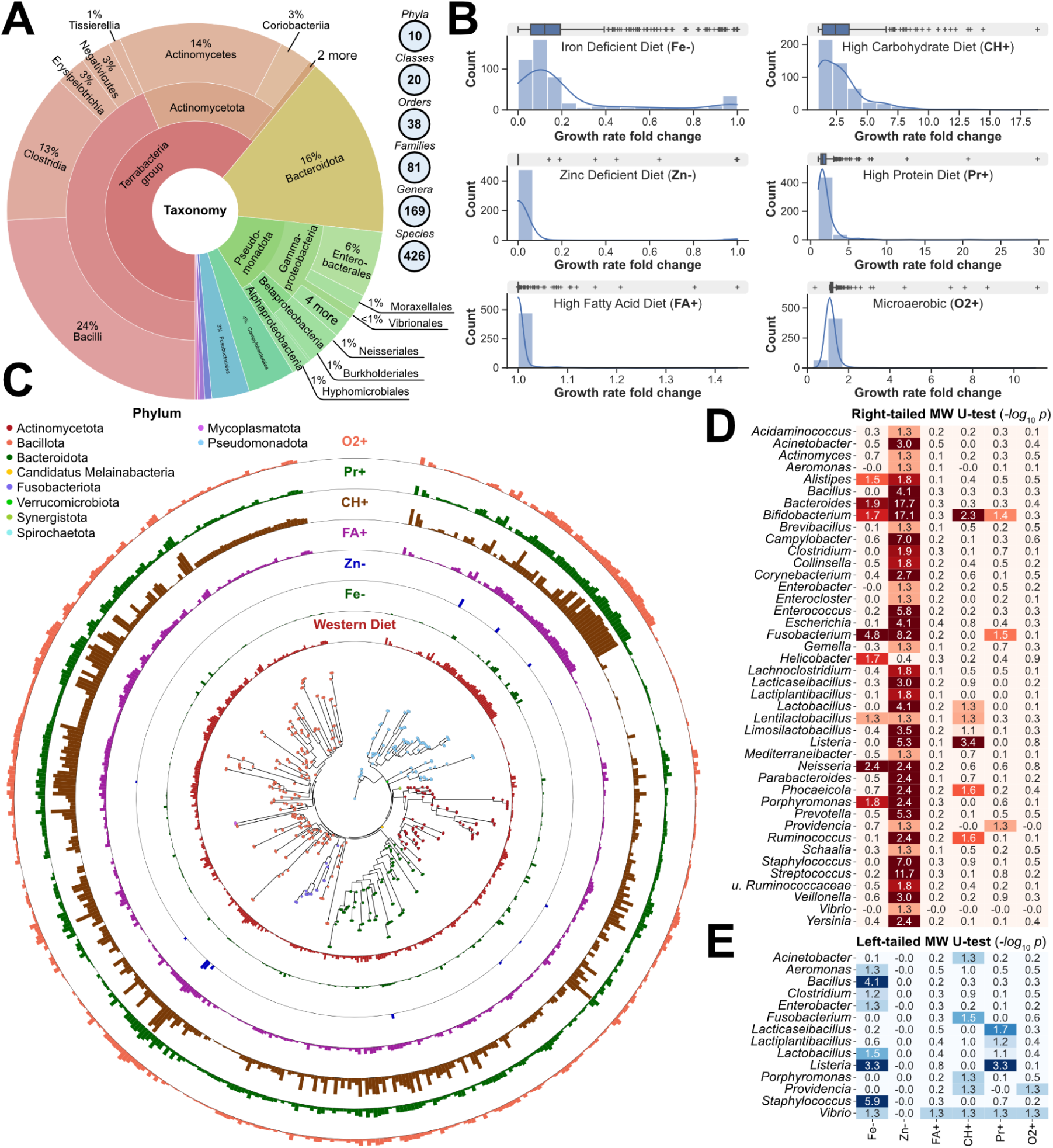
Reconstruction and predictions of 495 ME-models of human gut microorganisms under seven dietary conditions. **(A)** Taxonomic classification of models. **(B)** Phylogenetic tree (see color coded legend) of modeled members and distribution of their growth rates, grouped by phylum. **(C)** Histogram and boxplots of growth rate fold change distributions under the six diets compared to WD. Diets are iron deficiency (Fe-), zinc deficiency (Zn-), high fatty acid (FA+), high carbohydrate (CH+), high protein (Pr+), and microaerobic (O2+). **(D** and **C)** Genus enrichment analysis of members that are at an advantage (D) or disadvantage (Ee) in a certain diet compared to WD. Enrichment p-values were calculated with the Mann-Whitney U-test, right- and left-tailed for advantage and disadvantage, respectively. The heatmap shows p-values (*p*) as *-log_10_ p*. Significant enrichment is determined with *p<0.05*, equivalent to *-log_10_ p>1.3*.

A high carbohydrate diet is predicted to overrepresent *Listeria* (*p ≤* 10^-3^), *Bifidobacterium* (*p ≤* 10^-2^), *Phocaeicola* (*p =* 0.03), and *Ruminococcus* (*p =* 0.03). Carbohydrate transporter expression has been reported to be a determining factor in *Listeria* infection,^37^ and our simulations show 35% to 64% of these genes are upregulated, supporting that this genus is at a competitive advantage in a high carbohydrate diet. Furthermore, *Bifidobacterium* and *Ruminococcus* have been shown to be stimulated by fermentable carbohydrate abundance in the gut,^31,32^ corroborating our predictions. Interestingly, *Listeria* is predicted to be underrepresented in the high protein diet (p = 0.001), while *Bifidobacterium* (p = 0.03)*, Fusobacterium* (p = 0.05), and *Providencia* (p = 0.05) were overrepresented.

On the other hand, no significance was predicted for genera under the high fatty acid diet and the microaerobic environment, suggesting that these conditions affect genera evenly. However, when performing Mann-Whitney U-tests at the species level (Figure S9), we found significant overrepresentation solely for *Campylobacter jejuni* in O2+ (*p* = 0.05). This microorganism is a microaerophilic pathogen resistant to oxidative stress,^38^ which corroborates our simulations. Again, these findings showcase the predictive capabilities of ME-models, as no significance was predicted by M-models for *C. jejuni* or any other bacteria under this condition (Figure S7).

ME-model simulations also allowed us to assess the secretory profile in all tested dietary conditions. In total, 135 metabolites were identified as fermentation products, 16 of them ranked as the ten most excreted metabolites across all conditions, except Zn-. Specifically, acetate and formate were the first and second most frequently excreted metabolites, followed by succinate, phenylalanine, and propionate (Table S1). As the simulated conditions lack oxygen allowing M- and ME-models to predict overflow, we determined qualitative and quantitative differences in simulated secretion profiles. For instance, 46 M- and 56 ME-models simulated simultaneous acetate and butyrate excretion in WD (Figure S10). However, fluxes from M-models show a weak correlation with ME-model fluxes (Pearson Correlation Coefficient 0.83 for acetate, 0.61 for butyrate). Additionally, seven M-models predict acetate consumption with simultaneous butyrate production (Figure S10) or with other excreted metabolites (Table S2). Although acetate consumption is feasible, especially under the simulated rich medium condition,^39^ acetic acid import increases intracellular pH^40^ that an M-model can dissipate without proteome reallocation. On the contrary, none of the ME-models predict the consumption of acetate, reflecting a condition where gene expression constrains transporter synthesis to excrete proton overflow. This result further supports the necessity for ME-models to predict uptake and consumption profiles accurately.

The most abundant category of overflow metabolites was SCFAs, secreted under all tested conditions. Acetate ranked the highest (Table S3), followed by formate, propionate, butyrate, D-lactate, L-lactate, and 2-oxobutanoate. The most frequent groups of secreted metabolites after SCFAs were amino acids, alcohols, nucleosides, and dicarboxylic acids, although the rank order varied depending on the diet (Table S4). Interestingly, 72% of our ME-models under WD simulate simultaneous overflow of SCFAs and amino acids (mainly by members of the *Bacillota*, *Actinomycetota*, and *Pseudomonadota*), while only 24% of ME-models excrete SCFAs, but no amino acids. As the production and consumption of SCFAs by the gut microbiota have been associated with health and disease,^41^ and amino acid and amino acid-conjugated cholic acid abundances have been reported in IBD patients,^42^ we next studied the potential role of specific microbes in IBD, integrating ME-models with metagenomics and metatranscriptomics data.

### Elucidating metabolic and proteome allocation dynamics of the gut microbiota in patients with IBD

IBD is a lifelong medical condition characterized by chronic inflammation of the intestinal tract.^43^ The exact cause of IBD is still unknown and no effective treatment currently exists.^8^ However, complex interactions and correlation of various members of the human gut microbiota with IBD have been reported.^44^ Thus, metagenome and metatranscriptome experiments have been performed to identify associations of microorganisms with the onset and progression of the disease.^45,46^ A recent study found that these experiments reveal distinct properties of the microbiome in IBD, and that integrating those two data types provides a more comprehensive understanding of microbial dysbiosis.^8^ However, sequence data alone impedes the determination of potentially causal metabolic mechanisms in IBD. Here, we integrate multi-omics with ME-models to reveal net secretory changes in the human gut in IBD vs. Non-IBD and to identify the underlying drivers and their responsible microbes.

First, we processed metagenomics data^8^ using Songbird^7^ to quantitatively determine the abundance differentials (D) of bacteria across IBD and Non-IBD patients (Figure 4A). Six of the ten genera showing the largest differentials in IBD, namely *Enterobacter*^47^ (D = -0.93), *Campylobacter*^48^ (D = -0.91), *Fusobacterium*^49^ (D = -0.81), *Ureaplasma*^50^ (D = -0.79), *Weissella*^51^ (D = -0.77) and *Bartonella* ^52^ (D = -0.71) comprise of known pathogenic or opportunistic bacteria. Overgrowth of *Desulfovibrio* (D = -1.51), considered a commensal bacterium in the healthy gut, has been associated with human diseases.^53^ The observed presence of human pathogens among the largest negative differentials (Figure 4A and Table S5) agrees with reports that the abundance of pathogens in IBD correlates with the onset of multiple comorbidities.^54^ On the other hand, nine of the ten genera most associated with Non-IBD are not typically associated with infections (Figure 4A and Table S5), and only one species of *Dialister* (+1.17), *D. pneumosintes*, has been reported to be pathogenic.^55^

**Figure 4.**
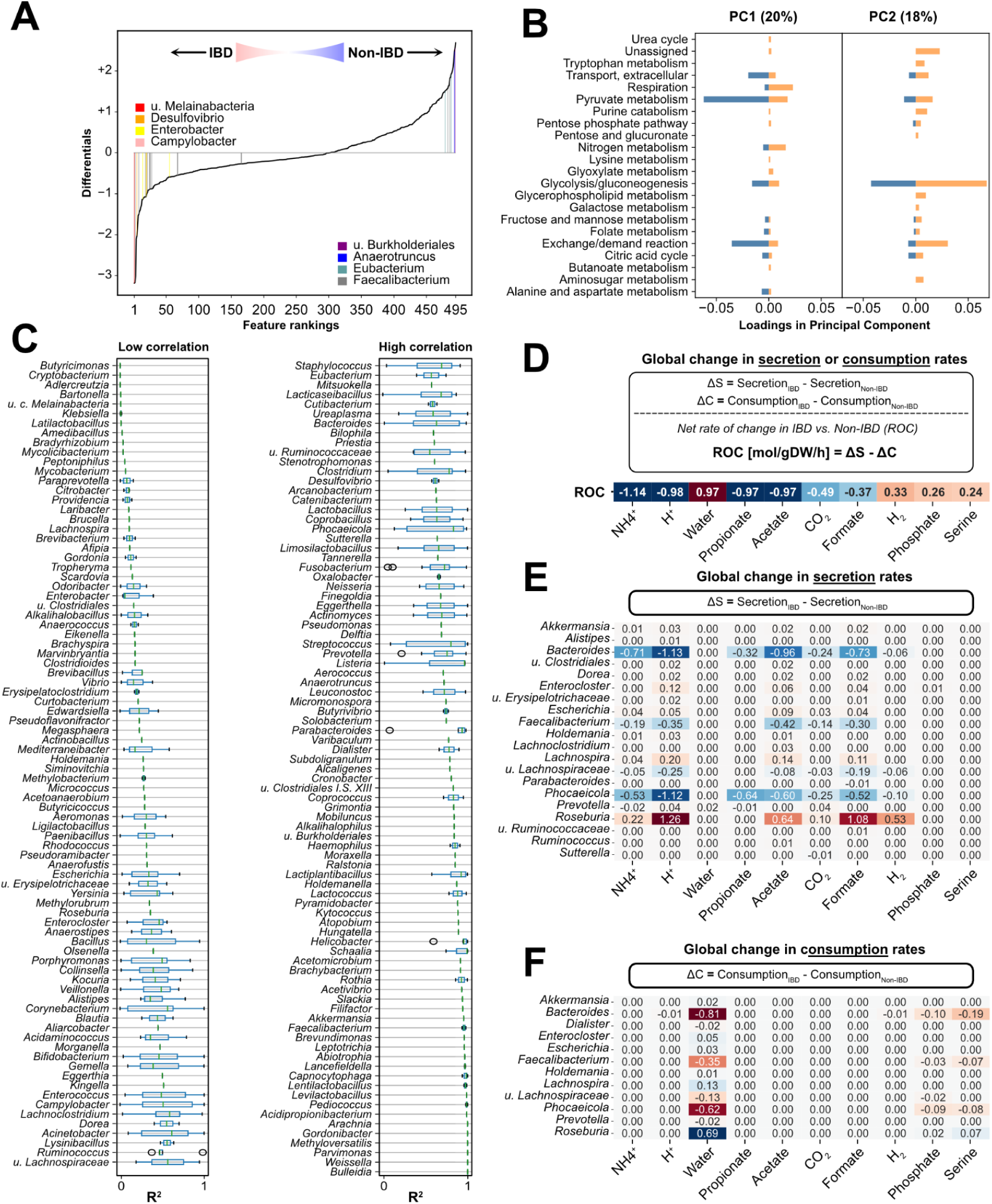
Multi-omics-integrated predictions of metabolic secretion in the gut in IBD vs. Non-IBD patients. **(A)** Differential abundance analysis using Songbird. The plot shows feature rankings after sorting microorganisms by their differential and the magnitude of the estimated differentials. Positive differentials indicate a stronger association with Non-IBD, and negative differentials show a stronger association with IBD. **(B)** Average loadings of metabolic subsystems in PC1 and PC2 of the performed PCA on predicted metabolic fluxes of 495 cME-models in IBD and Non-IBD. **(C)** Distribution of correlation (coefficient of determination, R^2^) between metabolic fluxes predicted in IBD and Non-IBD by each cME-model. Outliers are indicated by gray circles. **(D)** Net rate of change of ten metabolites with the highest variation. **(E** and **F)** Individual contributions of bacterial genera to the net rate of change in secretion **(E)** and consumption rates **(F)**. Contributions are colored in a gradient of red to blue according to their value, where red signifies a contribution to positive ROC, and blue signifies a contribution to negative ROC. The complete dataset for all predicted changes is provided in Table S6.

Next, we integrated metatranscriptomics data to generate condition-specific ME-models (hereinafter, cME-models) for IBD and Non-IBD and performed a Principal Component Analysis (PCA) on the metabolic fluxes (Figure S11). The loadings (coefficients) in the PC1 (20%) and PC2 (18%) show that the highest metabolic variance is explained by variance in growth rates, as shown by higher loadings of central carbon pathways, such as glycolysis, pyruvate metabolism, and amino acid synthesis (Figure 4B). The PCA revealed that metabolic fluxes in IBD and Non-IBD do not cluster separately, but individual organisms showed distinct flux distributions in the two conditions. This is exemplified by *Amedibacillus dolichus*, *Klebsiella aerogenes*, and *Klebsiella pneumoniae*, which have the highest Euclidean distance projected on the principal components of the PCA (Figure S11).

Then, we investigated the variations in the metabolic fluxes of individual organisms to elucidate which microorganisms exhibited different metabolism in IBD compared to Non-IBD. We calculated a coefficient of determination (R^2^) between its metabolic fluxes in IBD and Non-IBD (Figure 4C). A low R^2^ highlights genera with the highest predicted metabolic flux variation. We found high variability (R^2^ < 0.1) for metabolic fluxes of species within the genera *Butyricimonas*, *Cryptobacterium*, *Adlercreutzia*, *Bartonella*, *Latilactobacillus*, *Amedibacillus*, *Bradyrhizobium*, and *Mycolicibacterium*, among others (Figure 4C). On the other hand, several genera, such as *Bulleidia*, *Wiesella*, and *Parvimonas*, showed no variation between IBD and Non-IBD (Figure 4C). Interestingly, differential abundance analysis of genera (Figure 4A) did not correlate with their metabolic flux variation, as *Faecalibacterium* (D = 1.70, R^2^ = 0.99) and *Burkholderiales* strains (D = 2.56, R^2^ = 0.91) were predicted to have little metabolic variation (Figure 4C). However, some genera with higher abundance differentials showed higher metabolic flux variation, such as *Campylobacter* (D = -0.94, R^2^ = 0.53), *Eubacterium* (D = 0.74, R^2^ = 0.57), *Desulfovibrio* (D = 1.52, R^2^ = 0.62), and *Enterobacter* (D = -0.62, R^2^ = 0.16). Thus, there is a major discrepancy between abundance differentials and metabolic variations ^8^ (Figure S12) as indicated by a PCC of 0.095 (R^2^ = 0.09).

Finally, we integrated cME-models with microbial abundances to determine the net change in metabolites that become available to the host. We calculated the net rate of change (ROC) in IBD vs. Non-IBD of metabolite consumption and secretion by all bacteria to assess whether the host would be exposed to different metabolite levels in IBD compared to Non-IBD. High metabolite levels are associated with positive ROC values, while negative ROC values indicate metabolite depletion in the gut (Figure 4D). A breakdown of the ten largest ROC values highlights the main contributors to secretion (Figure 4E) and consumption (Figure 4F), elucidating key drivers behind secretory shifts in health and disease. The largest ROCs correspond to protons (H^+^), ammonium (NH ^+^), and SCFAs such as acetate, formate, and propionate. At the same time, there was a predicted increase in the net rate of hydrogen (H_2_), phosphate, and serine (Figure 4D). Complete consumption and secretion profiles are provided in Table S6.

The negative net rate of change of H^+^ (ROC *=* -0.98 mol/gDW/h) caused by decreased secretion rates (Figure 4E) indicates an increase in pH, corroborating previous findings that gastrointestinal pH profiles are higher in patients with IBD.^56^ Our simulations show that H^+^ secretion decrease is mediated mainly by *Bacteroides* (ΔS = -1.13 mol/gDW/h), *Faecalibacterium* (ΔS = -0.35 mol/gDW/h), *Phocaeicola* (ΔS = -1.12 mol/gDW/h), and *Lachnospiraceae* strains (ΔS = -0.25 mol/gDW/h), which outweighs the increase in H^+^ secretion by *Enterocloster* (ΔS = +0.12 mol/gDW/h), *Lachnospira* (ΔS = +0.20 mol/gDW/h), and *Roseburia* (ΔS = +1.26 mol/gDW/h).

Similarly, the net rate of change of SCFAs was predicted to be negative for acetate, formate, and propionate (Figure 4D and Table S6). SCFA production in the gut of patients with IBD has been reported to decrease when compared to Non-IBD as a result of shifts in the abundance of fermenting bacteria.^57,58^ In accordance with these observations, our simulations reveal that the negative ROC of SCFAs are a product of a major decrease in SCFA secretion (ΔS < 0) by *Bacteroides*, *Faecalibacterium*, *Phocaeicola*, and *Lachnospiraceae* strains (Figure 4E). However, a secretion increase (ΔS > 0) was predicted for *Roseburia* and *Lachnospira*, though at a smaller rate that was outweighed by the four genera with larger negative secretion change (ΔS < 0) (Figure 4E). It is worth noting that the predicted ROCs for SCFAs and H^+^ were correlated (Figure 4, E and F), as SCFAs are transported through proton symporters, which highlights the combined effect of the gut microbiome on SCFAs and pH levels in an IBD gut.

Interestingly, our simulations predict a positive net rate of change for serine (ROC = +0.24 mol/gDW/h) in IBD vs. Non-IBD (Figure 4D). Results determined that there is a decrease in the consumption of serine by *Bacteroides* (ΔC = -0.19 mol/gDW/h), *Faecalibacterium* (ΔC = -0.07 mol/gDW/h), and *Phocaeicola* (ΔC = -0.08 mol/gDW/h), countering the increase in serine consumption by *Roseburia* (ΔC = +0.07 mol/gDW/h) (Figure 4F). Serine levels in the gut are largely dependent on dietary intake,^59^ meaning that a reduction in its consumption by gut microorganisms is bound to increase its availability in the gut of IBD patients. Enterobacterial blooms are characteristic in IBD^45^ and it has been shown that pathogens such as adherent-invasive *E. coli* (AIEC), depend on serine catabolism for invasion in the inflamed intestine.^59^ Our cME-models propose a mechanism by which AIEC and other Enterobacteria are at a competitive advantage in the gut of IBD patients, which might contribute to their reported high blooming frequency.^59,60^

Finally, we explored how dietary changes can affect the IBD gut. We calculated the sensitivity of microbes in the IBD gut to dietary changes by simulating diets from our previous analysis (Figure 3) (Figure S13). Under a low iron diet, several genera are underrepresented, including *Bacteroides*, which we determined as a major contributor to changes in metabolite levels and pH in IBD. Conversely, *Bacteroides* and *Phocaeicola* are overrepresented in a low zinc diet, implying a strong potential to induce adverse effects in an IBD gut. While these predictions require extensive clinical validation, they further showcase the strength of ME-models for unraveling mechanistic insights.

## Discussion

Modeling the human gut microbiome and integrating it with multi-omics data is key to understanding disease progression and designing clinical interventions.^6^ While M-models have been applied for this, they have limitations, especially at the microbiome scale, such as fixed cofactor and micronutrient usage and biomass composition,^61^ and neglecting the biosynthetic costs of enzymes, allowing for biologically implausible fluxes.^18,62^ ME-models overcome these issues by associating functions with biomass components like proteins in metabolic conversion, RNAs in gene expression, and cofactors in enzymatic modification.^13^ This study demonstrated that ME-models can predict dysbiosis caused by zinc and iron deficiency, which M-models cannot, due to their fixed biomass compositions. Moreover, ME-models accurately predicted that acetate uptake is detrimental to the proton membrane potential and cellular homeostasis,^63^ which M-models failed to do. Although previous studies attempted to model biomass variation using M-models and experimentally derived biomass composition,^17,64^ the amount of detailed data required for such analysis is currently out of reach for studies of complex communities and microbiomes.

With the development of the automated tool coralME, we have solved the main bottleneck of ME-model reconstruction.^22^ CoralME was thoroughly tested on 21 diverse microbes (Figure 2 and Figure S4) and deployed to reconstruct 495 ME-models of gut bacteria from the AGORA resource,^9,24^ encompassing 178 genera from 14 phyla. Simulations predicted the metabolic advantages of microbial genera in the human gut under seven dietary conditions. We identified significant over- or underrepresentation of genera with increased carbohydrate and protein intake and decreased iron and zinc intake while higher lipid intake had little influence on the microbiota. These simulations contextualize several reports on the effects of dietary cofactors and micronutrients in shaping the microbiome,^27,28,35,37–39^ and provide novel insight on their mechanistic basis.

We integrated multi-omics data from human subjects to compare metabolic activity and quantitative abundance profiles of gut microbiota in IBD patients with healthy participants. Microbial relative abundance has been used as a proxy to determine the metabolic contribution,^6^ for instance, in the determination of drug metabolism performed by members of a microbiome.^9^ However, relative abundance suffers from sequencing depth bias,^7^ unable to determine the absolute abundance of a strain, and therefore, the correct metabolic contribution of a subpopulation in a microbiome. In turn, we performed differential abundance analysis using Songbird^7^ and reported metabolic variations from simulations. Simulations revealed that differential abundance analysis and metabolic shifts are complementary, indicating that abundance variations do not inform on metabolic variations, and vice versa, and thus, it is required to integrate both for a comprehensive understanding of the gut microbiome in IBD. Our analysis found pathogenic bacteria are differentially abundant in IBD patients, in line with reported high incidences of comorbidities.^54^ Integrating differential abundance with metatranscriptomic data resulted in context-specific ME-models, revealing metabolic variations across bacteria. For example, *Faecalibacterium* showed minimal adaptation in nucleotide metabolism, while *Butyricimonas* completely rearranges central carbon, nucleotide and amino acid metabolism in the IBD gut (Figure 4C). We also determined how microbiome metabolism changes could affect the host, revealing that major metabolic shifts were driven by a few abundant genera, namely *Bacteroides*, *Faecalibacterium*, *Phocaeicola*, and *Roseburia*. Overall, these simulations explain previously reported changes in intestinal pH,^56^ metabolite levels,^57,58^ and vulnerability to Enterobacterial blooms in IBD patients.^59^

While our findings cannot be immediately translated into clinical interventions, they can guide strategies to offset dysbiosis and comorbidities common in IBD. For example, a high-protein diet favors *Bifidobacterium* (Figure 3), beneficial in IBD,^65^ and does not cause metabolic dysbiosis (Figure 4). Conversely, oral iron supplementation, used in up to 80% of IBD patients,^66^ could exacerbate gut dysbiosis, suggesting intravenous iron as a potential alternative.^67^ Furthermore, a low zinc diet could be detrimental by favoring *Bacteroides* and *Phocaeicola*, supporting the reported positive effects of oral zinc supplementations.^68^ While an extensive clinical study would be necessary to evaluate these predictions, coralME enables the generation of high-quality ME-models quickly, facilitating systems biology studies at a massive scale, filling gaps in microbiome research, and aiding in the development of transformative intervention strategies.

## STAR Methods

### Key resource table

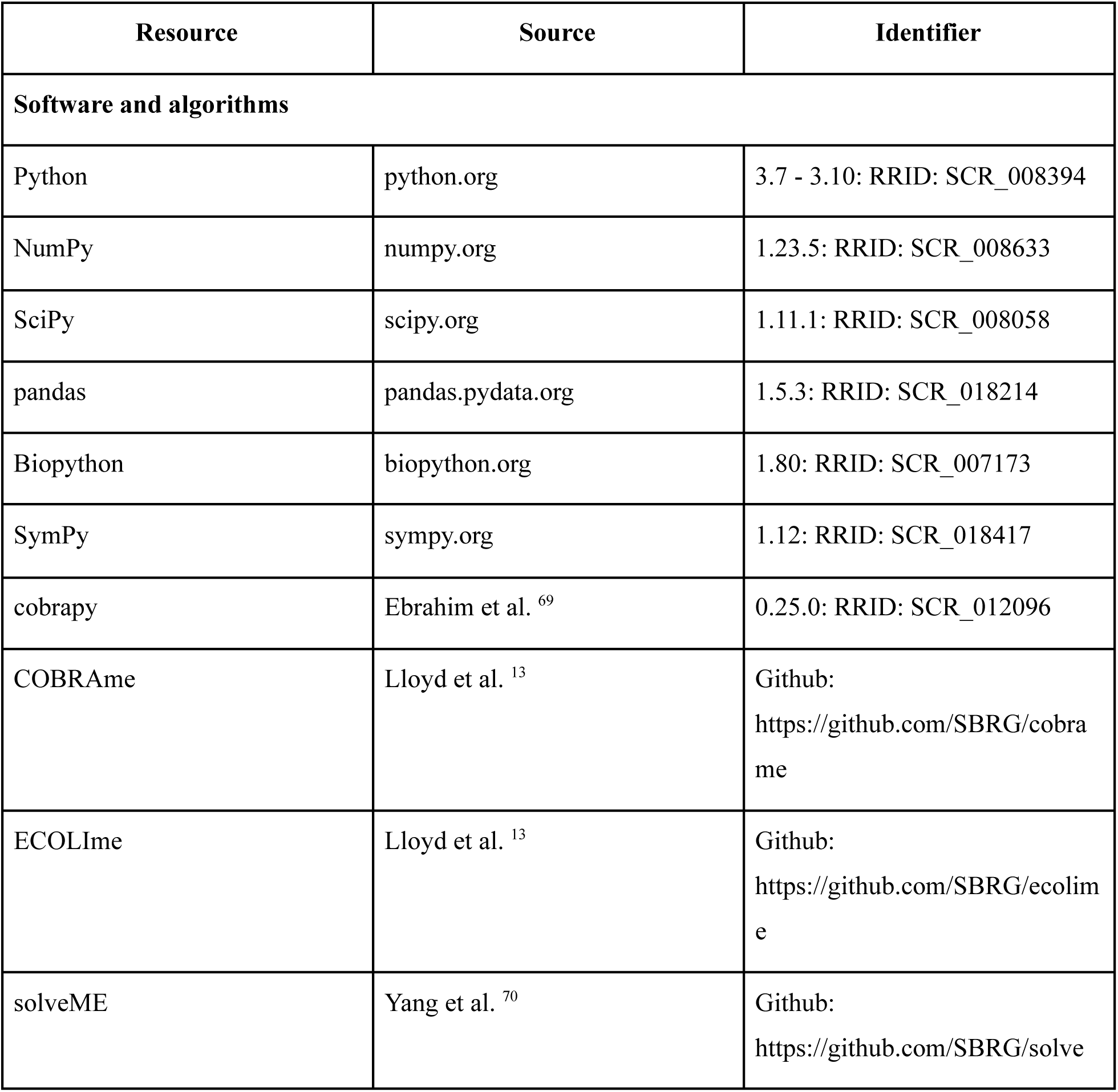
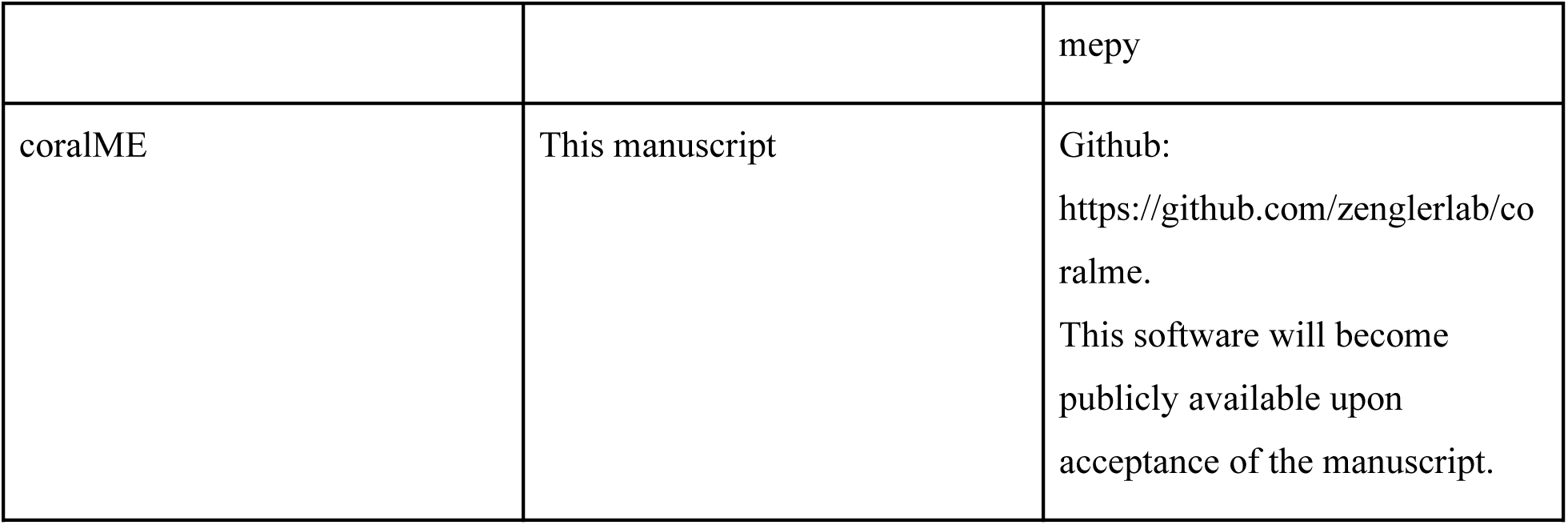

## Resource availability

### Lead contact

Further information and requests for resources and reagents should be directed to and will be fulfilled by the lead contact, Karsten Zengler.

### Materials availability

The reconstructed ME-models and all related files are available at https://github.com/ZenglerLab/coralME-models and for the gut bacteria at https://github.com/ZenglerLab/coralME-gut.

The coralME software is open-source and freely available under an MIT license from GitHub^71^/Zenodo and the Python Package Index (*Note: software will become publicly available upon acceptance of the manuscript*). All model reconstructions of this work were performed with coralME version 1.0.0, available at https://github.com/ZenglerLab/coralME.

## Method details

### Development and architecture of coralME

The coralME software was developed in Python as it derives from COBRAme,^13^ ECOLIme,^13^ and solveME,^70^ all of them available only from their GitHub repositories and not compatible with recent versions of COBRApy.^69^ coralME has been tested and is fully compatible with Python version 3.7 to 3.10, COBRApy version 0.25.0, and can use the solvers Quad MINOS,^72^ Gurobi Optimizer^73^ (Gurobi Optimization, Inc., Houston, TX), and the IBM ILOG CPLEX^74^ (IBM Corp., NY). The compatibility of coralME with the latter two linear programming (LP) solvers is included, as the user needs to install Fortran, obtain a license for the Quad MINOS source code, and compile it using f2py from the numpy python package^75^ for the specific computing architecture.^75^ Under Windows and MacOS, Gurobi is the default solver, configured to use quadruple-precision floating-point arithmetic, while CPLEX can only execute double-precision floating-point arithmetic. As expected, CPLEX returns faster optimizations and feasibility checks compared to Gurobi, but less accurately (Figure S14). The gut microbiome gained broad interest due to its association with chronic human diseases, as it shows hallmark compositional and metabolic traits. While composition is usually assessed via shotgun metagenomics, microbial presence alone does not provide insights into metabolic activity in vivo. Metabolic models have long been proposed to describe metabolic shifts in the human gut, but flux uncertainty and lack of constraints hinder their ability to produce robust simulations. Here, we deploy models of metabolism and gene expression (ME-models), which predict biomass requirements, account for cofactor utilization, and are constrained by the simulated proteome. We reconstructed 495 ME-models to describe metabolic responses to macro- and micronutrients that cannot be captured by standard metabolic models. Utilizing ME-models and multi-omics data from patients with IBD, we identified key metabolic shifts in their gut microbiome. Thus, results from ME-model reconstruction and multi-omics integration showcase the next-generation of data-based mechanistic modeling for microbiome research.

A significant fraction of modules in coralME were adapted from COBRAme, ECOLIme, and solveME.^13,70^ A derivation of the COBRAme software is included now in the *core*, *io*, and *util* modules of coralME, the *builder* module derives from ECOLIme, and the *solver* module derives from solveME. The imported code was thoroughly inspected and corrected. Corrections were incorporated to coralME, particularly for two calculations: the ATP requirement for RNA degradation after processing of rRNA operons and the stoichiometric coefficient of carrier proteins if the reaction is reversible. Regarding the former, COBRAme does not determine if a gene overlaps with another gene in the same transcriptional unit (TU), potentially leading to a negative value of the ATP requirement if the TU contains a non-coding RNA. In the case of the latter, COBRAme assigns a negative dilution term to a protein in the reverse reaction. Both calculations were reimplemented, setting new rules executed when the reaction stoichiometry is updated to avoid a negative value for the ATP requirement and setting dilution terms only to substrates. Another new capacity implemented in coralME is the modeling of generic components in metabolic reactions that COBRAme^13^ cannot perform. The use of generic components allows the modeling of poorly defined gene-protein-reactions associations that combinatorially explode into hundreds or thousands of reactions-complex associations in the ME-model.

### Inputs and file processing in coralME

As a minimal input, coralME requires an M-model and a compatible GenBank file^25^ to reconstruct an ME-model and optionally use additional information from BioCyc.^26^ Then, coralME automatically infers protein complex compositions and reaction associations from the M-model and obtains gene sequences from the GenBank file. For convenience, coralME will provide files (see Supplementary Text 1) that the user can edit to add new information or to override inferred associations and values. As a consequence, there is no need to generate multiple input files manually, as coralME was designed specifically to read, parse, and map data automatically to ensure the consistency of the models. Once all mandatory (and optional) files are provided, the reconstruction is fully automated and generates a draft ME-model that can be troubleshot to identify and repair potentially essential gaps. The entire data processing, ME-model reconstruction, and troubleshooting are documented step-by-step in log files. Missing or unresolvable issues in the input files are informed through detailed and compiled warnings (See Curation Notes, Figure 1C). Figure 1 highlights the source of mandatory and optional information used for the reconstruction and the overall architecture of the developed workflow.

The coralME software reconstructs ME-models in four steps: i) data reading and synchronization, ii) data complementation, iii) reconstruction, and iv) troubleshooting. In the first step, coralME reads input files that include an M-model (e.g. from BiGG^76^), a genome sequence and annotation file (e.g. from GenBank^25^), and optional files from BioCyc.^26^ The class *Organism* defines database-like objects. One *Organism* object contains all the necessary information of an organism (*org* in Figure 1C) to reconstruct an ME-model. Similarly, a second *Organism* object contains relevant information from a reference organism (*ref* in Figure 1C). While the ME-model for *E. coli* (MG1655 K-12) is currently the most comprehensive model, any organism’s ME-model can be used as a reference if it contains a GenBank file to perform a homology search and formatted files mapping information within the reference. If the user provides data from BioCyc, data synchronization is performed by the *MEBuilder* class (Figure 1C), completing GenBank annotations using BioCyc data. In the second step, further information is transferred from the reference organism’s genome and annotation by performing a Best Bidirectional Hit analysis using BLASTp.^77^ The ME-model features are then condensed into an Organism-Specific Matrix file (OSM) containing all the building blocks for the reconstruction. In the third step, the *MEReconstruction* class reads the OSM and a configuration file and automatically generates the ME-model. Then, in the fourth step, the *METroubleshooter* class evaluates the ME-model, identifies potential gaps in the M- and E-matrix through an algorithm that efficiently traverses different ME-model component categories, and tests it for feasibility at a defined growth rate (e.g. 10^-3^ h^-1^).

An important feature of the coralME workflow is the OSM file (See Figure 1C). It connects the data processing and reconstruction stages of the modeling process, allowing the user to explore the inferred information and gene function mappings in the ME-model. Since the OSM file includes information about the homology between target and reference organisms, the user can manually curate the OSM directly and skip the *Synchronize* and *Complement* steps (Figure 1C). Importantly, manual editing of the OSM file permits correction, deletion, and addition of relationships between genes and functions. New information can include, for example, stoichiometry of complexes, cofactor use, mapping of complexes to generic components or *meta-complexes* (e.g. ribosome), association to M-reactions or ME-model *subreactions*, and information regarding RNA modifications, tRNA to codon associations, associated chaperones and peptidases, subcellular location, and translocation pathways.

### Algorithm in coralME modeling steps

Step 1) Synchronize: a MEBuilder object reads the mandatory (*Genome* and *M-model*) and optional input files (*Genes, Proteins, RNAs, TUs,* and *Sequences*) and parses them into coralME-readable objects that are stored in Organism objects for both the main organism and the reference used as a template. Then, it merges the information coming from all files and raises warnings when contradictions occur and when expected information is missing altogether, e.g. ribosomal proteins or tRNA ligases, among others.

Step 2) Complement: the MEBuilder object performs BLASTp^77^ against a reference organism and complements input data to obtain a new configuration file with inferred inputs, the Organism-Specific Matrix (OSM), and internal representations of the reference and target models. The MEBuilder can be executed once, as the BLASTp result and the OSM are saved to independent, editable files. Steps 1 and 2 must be run again if the user modifies the M-model (e.g. adding, removing, editing reaction properties) as the default option is to overwrite the OSM and the inferred configuration file. The MEBuilder object (*builder*) contains an initialized and empty ME-model (*builder.me_model*), the processed input data (e.g. *builder.df_data*), the input data from the reference (*builder.ref*), data to reconstruct the target ME-model (builder.org), and the results from BLASTp (*builder.homology*). The MEBuilder method also creates a modified GenBank file and a modified M-model file. If BioCyc data is used, the GenBank file might contain corrected or new information regarding genes and sequences. Also, if provided data from BioCyc, coralME writes a new file including information regarding TUs (named “TUs_from_biocyc.txt”).

Step 3) Reconstruct: a MEReconstruction object loads the new configuration file, mandatory inputs (i.e. OSM, GenBank file, and M-model), and optional files from BioCyc to reconstruct a draft ME-model (dME-model). If coralME is run from Step 3, the *builder* object only contains the ME-model, and input data should be accessed through *builder.me_model.internal_data*.

Step 4) Troubleshoot: a METroubleshooter object tests the feasibility of the generated ME-model, and if the ME-model is not functional, it identifies and tests gaps in a predefined order. The predefined order to test is i) metabolites that are deadends in the ME-model, but not in the M-model, ii) cofactors, which are commonly added gaps when the M-model did not include them, iii) all metabolites that represent deadends, iv) all metabolites, v) charged tRNAs, vi) complexes, vii) RNAs, viii) proteins, viii) processed proteins such as lipoproteins, and ix) generic components, e.g, *generic_tRNA_GAG_glu L_c* that represents the Glu-tRNA^GAG^ ME-metabolite. For each group of components, the algorithm adds sink reactions for all of them and tests the model’s feasibility. If the ME-model is functional, the algorithm checks for feasibility if one sink reaction is removed at a time. After checking every new sink reaction, the troubleshot ME-model contains only sink reactions that are essential. To accelerate the identification of gaps, new sink reactions are filtered out if there is no flux through them. If the ME-model is not functional, the algorithm proceeds to the next group, appending the new components to test to the previous group, and checks again for feasibility removing one sink reaction at a time. Once troubleshooting of the ME-model is completed (i.e. the ME-model is functional), it is optimized for growth rate.

A number of new features are included in coralME to aid reconstruction and simulation. coralME relies extensively on the BioPython project to extract nucleotide sequences using TU start and end positions, to detect the translation table of CDS, and to obtain an amino acid sequence from the gene start and stop positions. The detection of the translation table allows, for example, modeling of *Mycoplasmas* and *Spiroplasmas* that encode tryptophan for the codon UGA, otherwise defined as a stop codon. Also, coralME detects programmed frameshifts that previously needed user input. Another important feature is the detection of tRNA features from the GenBank file and the automated building of the “tRNA to codon” and the “codon to amino acid” dictionaries that previously needed to be set manually by the user. Moreover, the user can set a variable mapping misacylation of tRNA or let the software infer it in the case of the non-discriminant tRNA^Glx^ and tRNA^Asx^.

For simulating ME-models, coralME relies on the numerical evaluation of the stoichiometric matrix and bounds using a predefined growth rate, i.e. an ME-model optimization is a series of feasibility checks.^70^ The new *feasibility*, *feas_gurobi*, and *feas_cplex* methods allow the user to determine if the ME-model is functional at any growth rate and to determine the vector of fluxes if the growth rate is feasible using Quad MINOS, Gurobi, and CPLEX, respectively. For Windows and MacOS users, the new *optimize_windows* method allows the use of Gurobi (default, quadratic-precision float-point arithmetic) or CPLEX (double-precision float-point arithmetic). Moreover, the user can set the symbolic variable to use as a growth rate (default ‘mu’); setting a different symbolic variable allows the exploration of growth rates in co-cultures and higher-order models when μ_n_ ≠ μ_m_. Also, the LP problem is determined through a refactored *construct_lp_problem* method that can be evaluated directly or indirectly using lambda functions. Both evaluation methods are included in coralME. The direct evaluation method is faster than the computation of lambda functions, being optimal for single feasibility checks (e.g. troubleshooting of ME-models) but prohibitively slower if enough repeated evaluations are required as in optimization methods.

### Automated reconstruction of ME-models

Reconstruction of ME-models using coralME is performed as shown in Figure 1C. coralME takes two mandatory inputs (*Genome* and *M-model*) and five optional inputs (*TUs*, *Genes*, *Sequences*, *RNAs*, and *Proteins*) that are downloaded from BioCyc (see documentation). Combined, these inputs provide coralME with TU composition, gene sequences, gene function, RNA types, complex compositions, and protein locations.

The *Genome* file corresponds to a genome GenBank file of the organism, which must be functionally annotated and contain the genomic sequence. The *M-model* file is an SBML or JSON file containing the M-model of the organism whose ME-model is being reconstructed. It is necessary that the locus tags in *Genome* are consistent with the gene identifiers used in the *M-model*, which is usually the case if the GenBank files are downloaded from the M-model reconstruction publication. For organisms in the BiGG ^76^ database, the direct links to the GenBank files are available.

The five optional files (*TUs, Genes*, *Sequences*, *RNAs,* and *Proteins*) can be downloaded from BioCyc’s Special SmartTables of the preferred organism. A suitable BioCyc database must contain the gene identifiers used in the M-models reconstruction (see “Description of Inputs” in Supplementary Text 1). If a suitable BioCyc database is unavailable, the ME-model can be reconstructed without these files (only *Genome* and *M-model*). Still, the final ME-model will lose resolution regarding TU composition, organism-specific complex compositions, and their subcellular locations. For example, in the case of *E. coli* the files to download are under the names: “All genes of E. coli K-12 substr. MG1655” (*Genes* and *Sequences*), “All proteins (polypeptides + protein complexes) of E. coli K-12 substr. MG1655” (*Proteins*), “All RNAs of E. coli K-12 substr. MG1655” (*RNAs*), “All transcription units of E. coli K-12 substr. MG1655” (*TUs*). When downloading this information for other organisms, the SmartTables have the same name, with the organism name replaced accordingly. For some files, some columns must be manually added, so we provide a tutorial on how to download these files in our documentation (Supplementary Text 1).

All analyses of this work were performed with coralME version 1.0.0, available at https://github.com/ZenglerLab/coralME. A complete description of coralME’s architecture, inputs and outputs, and tutorials is provided in Supplementary Text 1. The reconstructed ME-models and all related files are available at https://github.com/ZenglerLab/coralME-models in coralME-models/clean/.

Computations were performed on a 64-bit Ubuntu 22.04.3 LTS (Jammy Jellyfish); AMD Ryzen 9 7900X @ 4.70 GHz (12 cores, 24 threads); 4 × 32GB 6000 MHz DDR5 RAM.

### Solving the non-linear programming problem in ME-models

We kept the same formulation of the non-linear programming (NLP) in ME-models as described in COBRAme,^13^ which is as shown in Equation 1. The coupling constraints and complete formulation of the NLP is described in detail by O’Brien et al.^14^. The NLP is solved using a binary search algorithm described by Yang et al.^70^

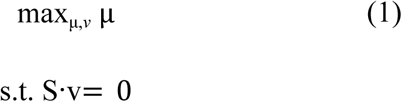

Simulations were performed using the Quad MINOS software courtesy of Prof. Michael A. Saunders at Stanford University.^70,72^ The Quad MINOS solver was compiled under Ubuntu 22.04 with gfortran version 5 and python 3.10 (pip 22.3.1, wheel 0.38.4, numpy 1.21.6, scipy 1.7.3, cython 0.29.32, and cpython 0.0.6). coralME provides compiled Quad MINOS solvers for users using python 3.7 to python 3.10 and Ubuntu with gfortran version 5 only.

### Update of the *E. coli* ME-model, simulation, and benchmarking

The input files to reconstruct the updated *E. coli* ME-model are as follows: The assembly NC_000913 version 2 from NCBI (https://www.ncbi.nlm.nih.gov/nuccore/NC_000913.2/), the iJO1366b^78^ M-model, and corrected input data used to reconstruct *i*JL1678b-ME.^13^ The input data was modified to comply with new formatting requirements. The information from *i*JL1678b-ME includes complexes stoichiometry, protein modifications, generics, RNA modifications (including their position), RNA modifiers, and the stoichiometry of *meta-complexes* (e.g. degradosome), enzyme associated to processes (e.g. translation initiation), among other types of information. The ME-model can be reconstructed using the Jupyter notebook and companion files in the GitHub repository (https://github.com/ZenglerLab/coralME-models/) in coralME-models/published/ecoli. The protocol to reconstruct the ME-model skipped the BLASTp step to determine homology with a reference genome, but it includes the troubleshooting step.

The maximum growth rate was set to 1.0 h^-1,^ and the maximum growth rate error was set to 10^-6^ h^-1^. After simulation, reaction name standardization was performed to pair and compare the majority of the reactions to *i*JL1678b-ME. Null fluxes were discarded, and non-zero fluxes were transformed using the log10 of the absolute value prior to comparing them using a linear regression (scipy.stats.linregress function, scipy version 1.9.0, python 3.10). The absolute differences of fluxes were grouped into intervals. For fold-change analysis, the flux in the coralME-model was divided by the flux obtained from *i*JL1678b-ME. Flux ratios coralME/COBRAme were then transformed using log2 and grouped into intervals.

Unique reactions were compared manually if they were refactored due to non-equivalent changes. Six non-zero reactions are unique to COBRAme, and 84 reactions are unique to coralME. Regarding the reactions unique to *i*JL1678b-ME, three are sink reactions for deleted components (RNase m5, m16, m23) and have no comparable fluxes in coralME. The remaining three reactions consist of one reaction that models the biosynthesis of lipoyl-ACP from octanoyl-ACP (*acp_lipoate_synthase_FWD_SPONT*), a reaction that is now included as a subreaction in coralME; and two reactions that synthesize the holo-ACP complex (*formation_EG50003-MONOMER* and *ACPS11_FWD_EG12221-MONOMER*) were refactored into one reaction (*formation_EG50003-MONOMER_mod_pan4p (1)*, Figure S15 and Table S7). Complete summary of manual curation of *i*JL1678b-ME performed is shown in Table S8.

### Update to the ME-models of B. subtilis, C. ljungdhalii, and T. maritima

We reconstructed draft ME-models of *B. subtilis*, *C. ljungdhalii*, and *T. maritima* using the M-models and gene annotation files provided with each publication. The associated M-models were *i*YO844,^79^ *i*JL680^20^ (a derivative model from *i*HN637^80^ with new metabolic reactions and genes), and *i*LJ478,^81^ respectively. Genome assemblies were NC_000853, NC_014328, and NC_000964, respectively. The optional files were downloaded from BioCyc as described in the documentation (Supplementary Text 1).

The draft ME-models were updated using the information provided in each publication by adapting all files to coralME-complying formats. We included all previously defined protein complexes, TUs, tRNA and rRNA modifications (with updates as mapped by coralME), and enzyme-reaction associations. Curation was performed until no further troubleshooting or gap-filling was required. The draft ME-models are available at https://github.com/ZenglerLab/coralME-models in coralME-models/clean/ under the *bsubtilis*, *cljungdahlii*, and *tmaritima* directories. The updated ME-models and all related files are available in coralME-models/published/ under the aforementioned directory names.

### Reconstruction and analysis of draft ME-models from peer-reviewed manually curated M-models

We reconstructed the draft ME-models of 21 bacteria (4 included in previous ME-modeling studies and 17 new ones), which included the most recent M-model of all 18 unique bacterial species available in BiGG^76^ (Table S9). The genome annotation files were retrieved from their respective sources in the BiGG database to the genome in NCBI. For the remaining four, *Liberibacter crescens,*^82^ *Mycoplasma mycoides,*^83^ *Vibrio cholerae,*^84^ and *Nitrosomonas europaea,*^85^ M-models and genomes were retrieved from their respective publications. We then downloaded the necessary BioCyc database files following the protocol described in Supplementary Text 1.

### High-throughput reconstruction and analysis of 495 ME-models derived from AGORA

We reconstructed 495 ME-models derived from AGORA.^24^ First, we downloaded all available M-models from https://www.vmh.life/files/reconstructions/AGORA/1.03/reconstructions/ and their corresponding genomes from https://www.vmh.life/files/reconstructions/AGORA/genomes/. The genomes include coding DNA sequences (CDS) but not their annotations, which are important for the identification of the ribosomal, RNAP, sigma factors, and aminoacyl-tRNA synthetase proteins. We annotated the gene sequences translating them first to protein sequences and using KofamScan,^86^ to then integrate the gene sequences and annotations into GenBank files using in-house scripts. As the genome files from AGORA do not include the whole genome sequence and neither identify readily rRNAs nor tRNAs, genome sources obtained from AGORA2^9^ were used to download genomes from NCBI and match files to M-models. The genomic sequences and tRNAs and rRNAs annotations were extracted from the downloaded NCBI genomes and copied to GenBank files containing gene annotations for protein sequences. Once genomes and M-models were paired, the large-scale reconstruction of ME-models was performed in parallel.

The original genome sequences consisting mostly of CDS were complemented to obtain rRNA and tRNA sequences using matching genomes from NCBI. A number of problems were identified before, during, and after reconstruction that reduced the number of potential ME-models from 773 to 495 ME-models (see Figure 3A). Specifically, 17 strain names did not match, reducing the input number of M-model/GenBank pairs to 754. During data processing, 154 M-models reported no genome coverage with the corresponding GenBank file, and reconstruction was halted. Then, 105 finished dME-models reported incomplete tRNA sets, as one or more essential tRNA genes were missing from the NCBI files. Although coralME can troubleshoot missing tRNAs, we discarded dME-models with gap-filled expression machinery as they are not suitable for further analysis. Finally, we obtained 495 dME-models, covering 169 different genera (Figure 3A).

For simulations of ME-models, medium composition for a “Western Diet” (WD) was retrieved from https://github.com/opencobra/COBRA.papers, 2018_microbiomeModelingToolbox folder. In the case of low iron, low zinc, and microaerobic conditions, WD was modified as follows: i) For metal ion deficiency cases, zinc and iron, the lower bounds of exchange reactions were set to 1% of the predicted optimal value in the reference case. ii) For high carbon availability cases, such as carbohydrate, protein and fatty acids, the exchange reaction lower bound was set to –2.0 for all metabolites in WD that were classified in these groups respectively. This ensured excess, i.e. the growth rate is not limited by carbon sources. iii) For low oxygen availability, we set the oxygen exchange reaction lower bound to –0.01 mmol/gDW/h, which is a low value compared to typical oxygen uptake rates of bacteria (in the order of 1.0–10 mmol/gDW/h).^87^ To compare fluxes, M-models were modified with the specific diet composition and simulated using the quad MINOS solver through the coralME interface.

A phylogenetic tree for the genomes of the resulting 495 ME-models was obtained using PhyloPhlAn,^88^ and growth rate and flux data were plotted using GraPhlAn.^89^ Pearson Correlation Coefficients were determined using the scipy.stats.pearsonr function,^90^ and Principal Component Analysis was performed using scikit-learn 1.3.1,^91^ with sklearn.decomposition.PCA, with two principal components.

### Metagenomics and metatranscriptomics sequence alignment from IBD and Non-IBD samples

Raw metagenomics and metatranscriptomics sequencing data for 78 samples from IBD and Non-IBD samples were downloaded from the Sequence Read Archive (BioProject Accession PRJNA389280). Raw reads were trimmed using Trim Galore version 0.6.10 (DOI 10.5281/zenodo.7598955). Trimmed reads were then aligned using Bowtie2^92^ to a *pangenome* generated from genomics or gene sequences for all 495 genome sequences from the AGORA resource included in this study. Reads were aligned to either the *pangenome* from genomic sequences (for metagenomics) or gene sequences (for metatranscriptomics). Read counts were calculated using Woltka version 0.1.5.^93^ Sample relative abundance was calculated from the metagenomics read counts by dividing the raw read count by the total number of reads in the sample.

### Integration of metatranscriptomic data in cME-models

Metatranscriptomic data was harnessed to develop condition-specific ME-models (cME-models) of the gut microbiome, distinguishing between IBD and Non-IBD states. The integration of metatranscriptomic data into these models involved the determination of a binary flux-constraining factor (FCF) derived from gene expression levels, by categorizing genes as active (FCF = 1) or inactive (FCF = 0) based on their expression. We defined inactive genes as those with zero aligning read counts from any sample, and active otherwise. FCF values served to modify the upper bound of the transcription rate of mRNA, *u_m_*, in transcription reactions (FCF\**u_m_*).

### Differential abundance analysis of metagenomic data in IBD vs. Non-IBD

The compositional nature of metagenomics data makes controlling false positives challenging.^94^ In order to mitigate the bias due to compositional effects, we conducted differential abundance analyses on the IBD/Non-IBD metagenomic data using Songbird v. 1.0.4 through QIIME2 v. 2020.6.0.^7^ The reference frame must be present in all samples and show a low variation (small differential) between IBD and Non-IBD. Furthermore, the reference frame should not be in low abundance, as their stability can be confounded with the low accuracy in detecting it. When a stable reference frame is identified, abundance ratios and metabolic and gene expression rates can be integrated to estimate global metabolite secretion and uptake rates by bacteria in the gut in IBD vs. Non-IBD. We used the differentials to choose a suitable reference frame to describe the metabolic and secretory imbalance associated with IBD leveraged by condition-specific ME-models of gut bacteria. When sorting organisms increasingly by the magnitude of the differentials, *Bacteroides thetaiotaomicron, Bacteroides intestinalis*, and *Bacteroides sp. 20/3* show high read counts above the average of 10^4.6^ per organism (Table S4). However, *B. thetaiotaomicron*^95^ and *B. intestinalis*^96^ have been shown to significantly change in abundance in previous IBD studies, indicating that *Bacteroides sp. 20/3* is a more suitable reference frame. In our results, *Bacteroides sp. 20/3* is present in all samples, with an average read count of 10^5.2^ in the 87th percentile. For comparison, the other bacteria with smaller differentials had an average read count of 10^3.8^. We therefore chose *Bacteroides sp. 20/3* as a reference frame. We used abundance ratios with respect to the chosen reference frame to weigh individual metabolic and gene expression fluxes of the 495 cME-models to estimate global metabolic and secretory shifts.

### Estimation of the net rate of change of metabolites by bacteria in the gut

Exchange rates were calculated using the following procedure. Given a metabolite *m*, an organism *i*, and a condition *c* ∈ {IBD, Non-IBD}, let *m* be exchanged (secreted or consumed) by *i* in *c* at a specific rate *v*_*m,i*_(*c*) in mmol/gDW /h. With an absolute abundance of *i* in *c*, *X*_*i*_ (*c*) in *gDW_i_*, the mass balance equation describing the net rate of change of *m* by all bacteria in *c*, *r*_*m*_ (*c*) in mmol/h, is shown in Equation 2.

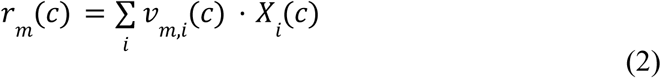

Using differentials estimated by Songbird,^7^ we choose a reference that can be expected to stay stable, which we then use to calculate abundance ratios. With a reference organism *ref*, the abundance ratio of *i* with respect to *ref* in *c* is defined as 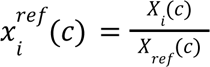 in 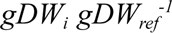 Abundance ratios are calculated for all *i* dividing the read counts of each sample in *c* by the read counts of *ref*, and then averaging to get one value for each condition *c*. Thus, we define the net rate of change of *m* with respect to *ref* in *c* as 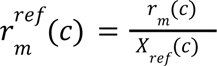 in mmol/gDW_ref_/h, which can be estimated as shown in Equation 3 using 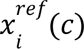 and *v*_*m,i*_ (*c*) calculated by the cME-models for all *i*. If 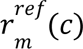 is positive, *m* is predicted to have net secretion in *c*, and if it is negative, *m* is predicted to have net consumption in *c*. Otherwise, there is no net exchange of *m* in *c*.

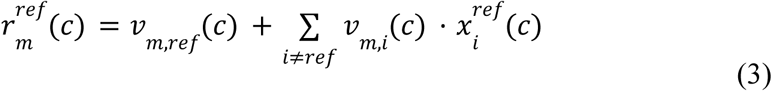

As *X_ref_* is expected to stay stable, we can assume that *X_ref_* (IBD) ≈ *X* (Non-IBD), so that 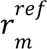 (IBD) and 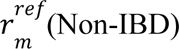 can be compared in the same way as *r* (IBD) and *r_m_* (Non-IBD). That is, if 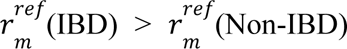 then *r_m_* (IBD) > *r_m_* (Non-IBD), and if 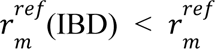 (Non-IBD) then *r_m_* (IBD) < *r_m_* (Non-IBD). Finally, we calculated the net rate of change of *m* in IBD vs. Non-IBD, ROC*_m_*, as shown in Equation 4. ROC is a combination of metabolite secretion and consumption by gut bacteria, so positive ROC values are favored by higher secretion rates and lower consumption rates, and vice versa.

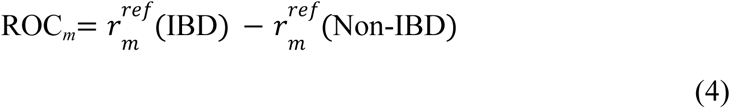

## Acknowledgments

JT would like to thank Oriane Moyne and Deepan Thiruppathy for their guidance and fruitful discussions on meta-omics data processing and alignment. JT and RS would like to thank Maxwell Neal for his meaningful input on the automated pipeline’s development and testing. All authors would like to thank Victor Nizet for improving the manuscript’s language and overall wording.

## Author contributions

Conceptualization: JTB, RSP, and KZ

Methodology: JTB, RSP and KZ

Investigation: JTB, RSP, YW, MK and KZ

Visualization: JTB and RSP

Supervision: KZ

Writing—original draft: JTB, RSP, YW, MK and KZ

Writing—review & editing: JTB, RSP, YW, MK and KZ

