## Supplementary Information for "Proteome allocation of the microbiome reveals how diet and metabolic dysbiosis impact disease"

This document includes:

Figures S1 to S21

Tables S1 to S3

Table S1 to S10

Document S1 and S2

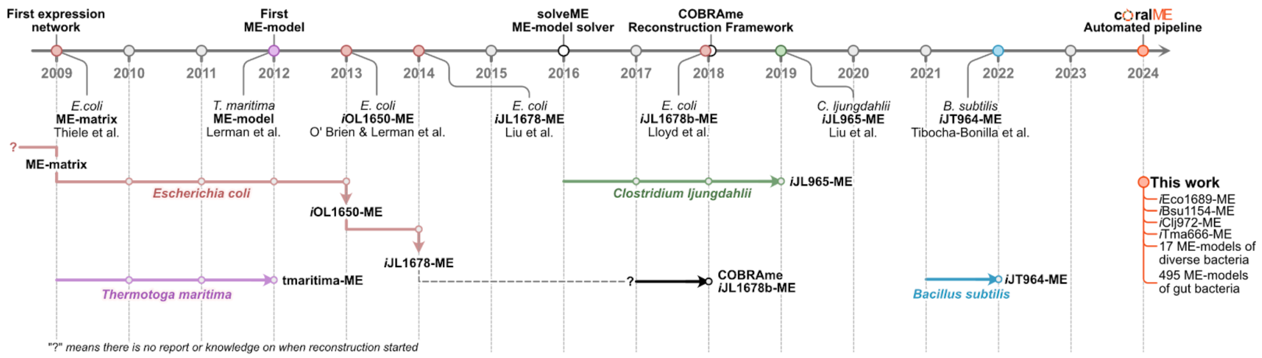

**Figure S1. Milestones in ME-modeling.** Reconstruction times for previously published ME-models and ME-modeling software.

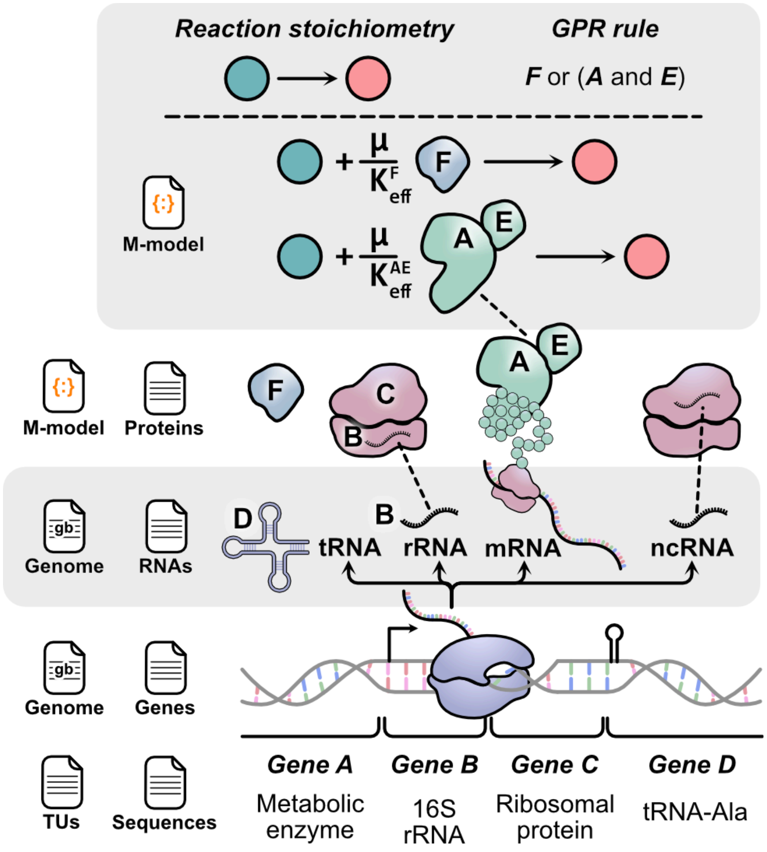

**Figure S2. A breakdown of the data derived from input information**. Transcription unit (TU) composition, gene sequences, and gene annotation are provided by four files (three of them optional): *Genome* in GenBank format with the nucleotide sequence (mandatory), and the *TUs*, *Genes*, and the *Sequences* file in FASTA format. The *Genome* file and the optional *RNAs* file specify the type and sequence of RNAs to simulate. Next, the *M-model* file (mandatory as either SBML or JSON format) and the optional *Proteins* file define complex compositions and locations. Finally, the *M-model* file provides the metabolic reaction network and its associations with enzymatic complexes.

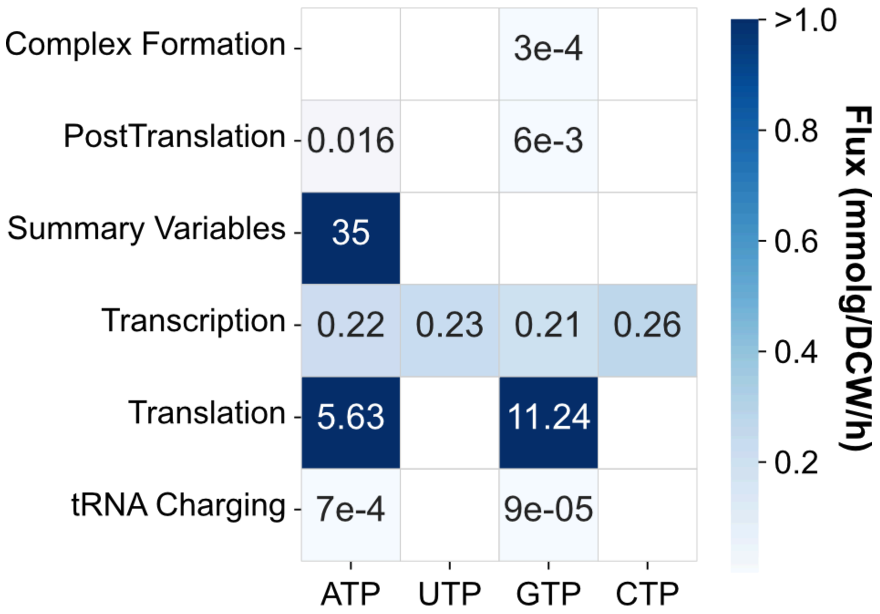

**Figure S3. *i*JL1678b-ME energy requirements per type of reaction.** Specific ATP maintenance requirements for growth per type of reaction for the model *i*JL1678b-ME. The M-matrix and E-matrix require a combined amount of 16.89 mmol ATP equivalents/gDW, underestimating the total GAM by 1.24 mmol ATP/gDW if compared to the *i*JO1366[1] model.

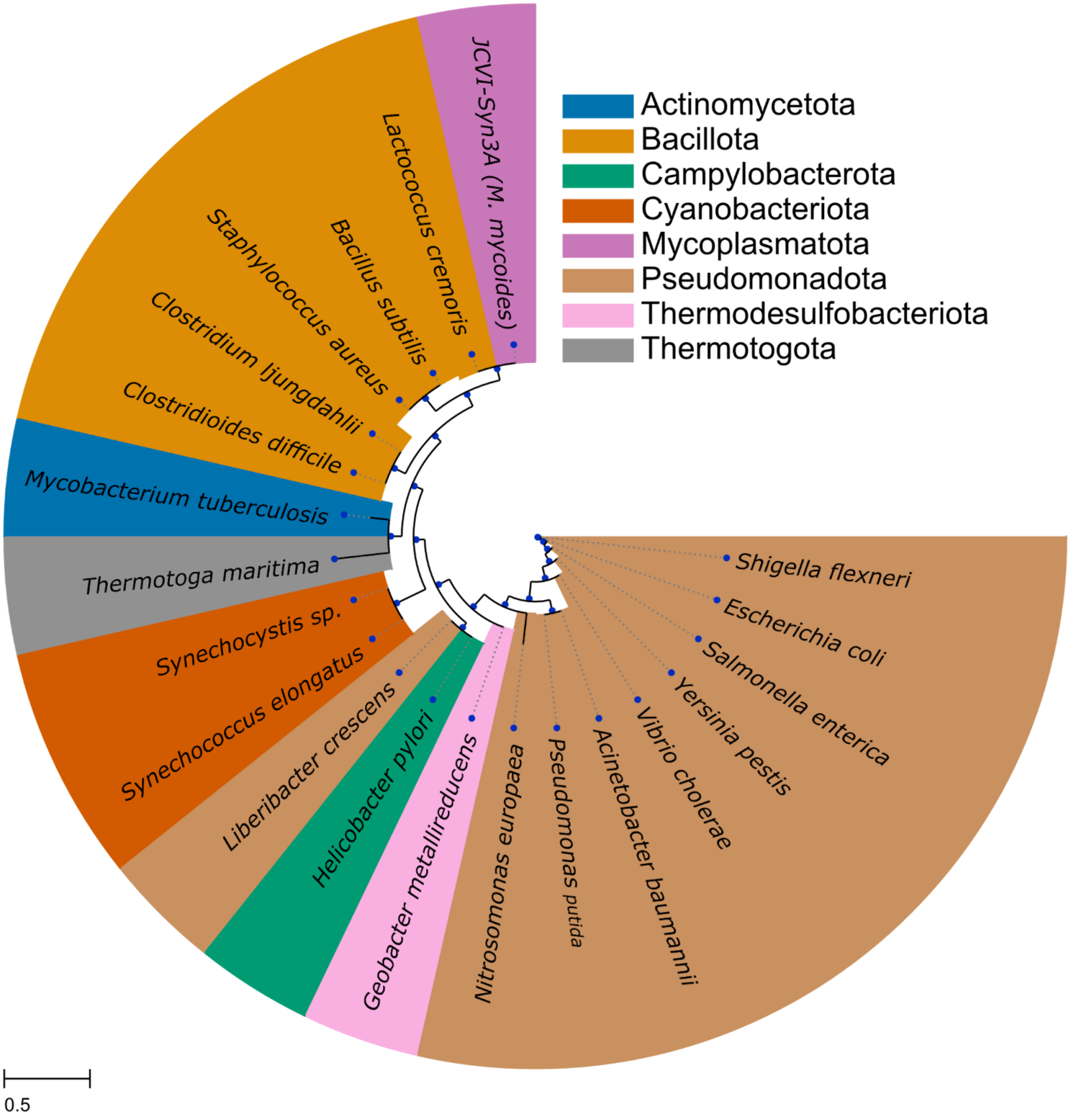

**Figure S4. Phylogenetic tree.** Phylogenetic tree based on the 16S ribosomal gene for all 21 bacteria for which ME-models were developed in this work. One of the 16S ribosomal genes was selected from GenBank files for each bacterium, aligned using Clustal Omega version 1.2.4, and the phylogenetic tree was reconstructed using FastTree version 2.1.11.

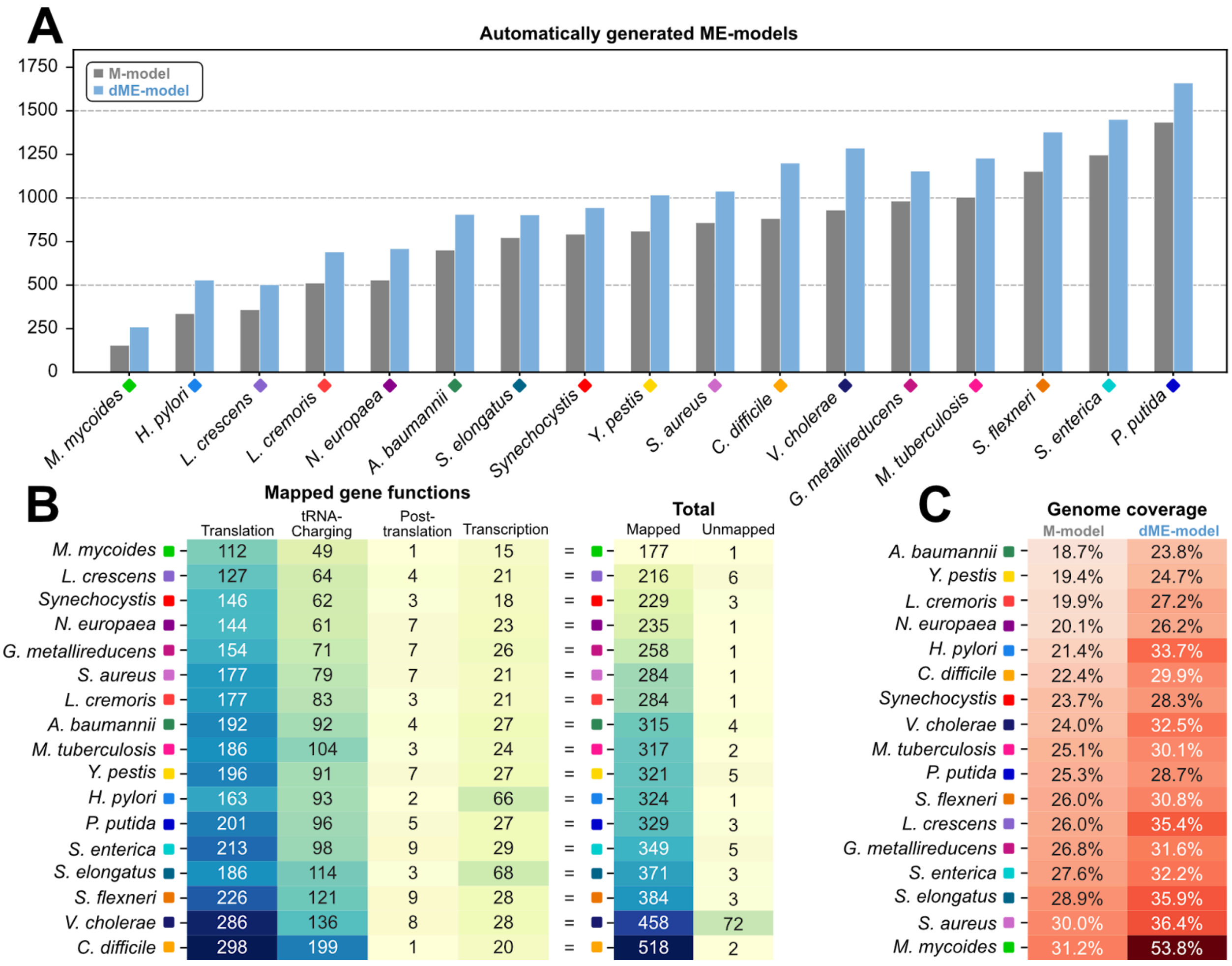

**Figure S5. Properties and benchmarking of drafts and curated ME-models reconstructed from peer-reviewed M-models. (A)** Comparison of the number of genes in all M- and dME-models, sorted by gene number in the M-models. **(B)** Comparison of mapped gene functions in M- and dME-models, sorted by total number of mapped gene functions in the dME-models. **(C)** Comparison of genome coverage in M- and dME-models, sorted by genome coverage in the M-models.

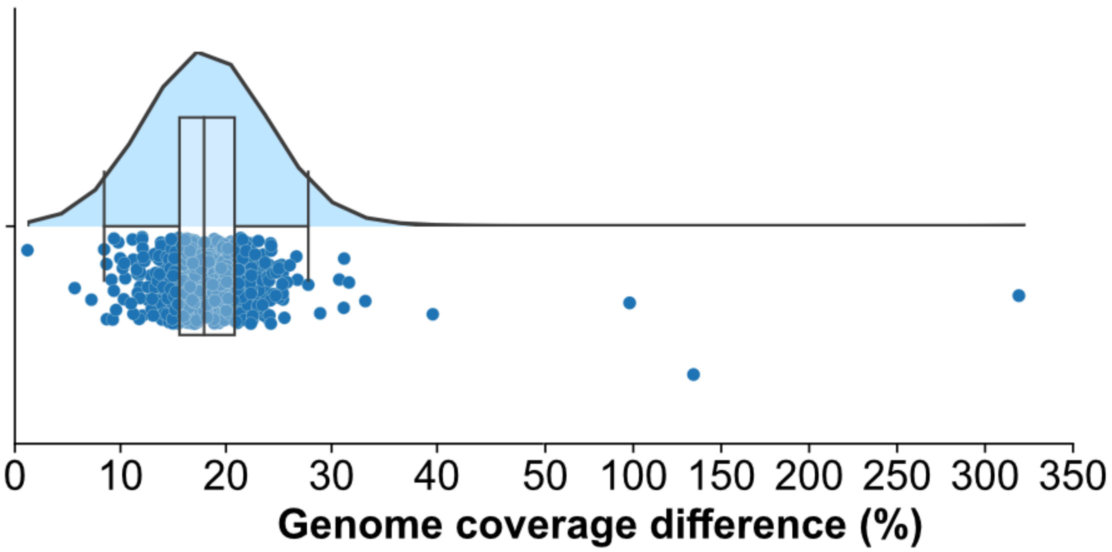

**Figure S6. Genome coverage in ME-models vs M-models for 495 gut bacteria.** Distribution of the percentual difference in the number of genes included in ME-models of gut bacteria compared to the M-models retrieved from AGORA. On average, the 495 ME-models contain 19.2% more genes than their counterparts.

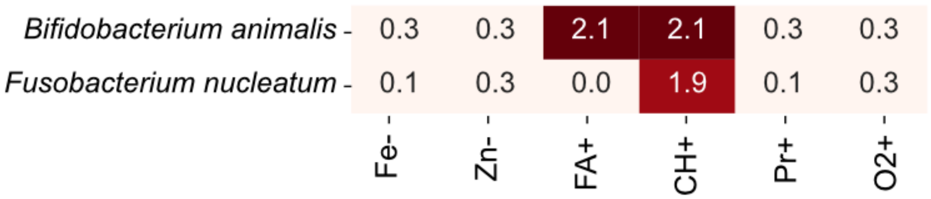

**Figure S7. Species enrichment analysis from M-model predictions of members that are at an advantage under the six cases of study compared to WD.** Enrichment p-values were calculated with the Mann-Whitney U-test, right- and left-tailed for advantage and disadvantage, respectively. The heatmap shows p-values (p) as -log10 p, so significant enrichment is determined with $p<0.05$, equivalent to -log10 $p>1.3$.

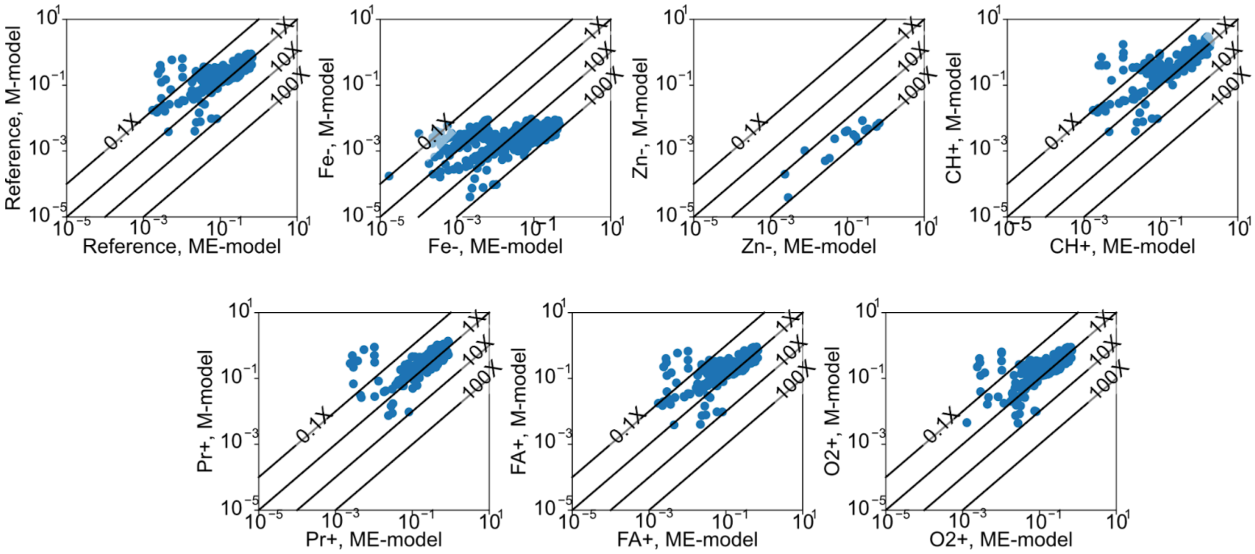

**Figure S8. Predicted growth rates by ME-model vs. M-model under seven conditions.** Comparison of the growth rates predicted by the reconstructed 495 ME-models and their corresponding M-model under a "Reference" diet (WD, see Online Methods) and six derived diets. Lines show where a ME-model predicted growth rate is identical to the M-model (1x), ten times faster (10x), 100 times faster (100x), and 10 times slower (0.1x) compared to the predicted growth rate by the corresponding M-model.

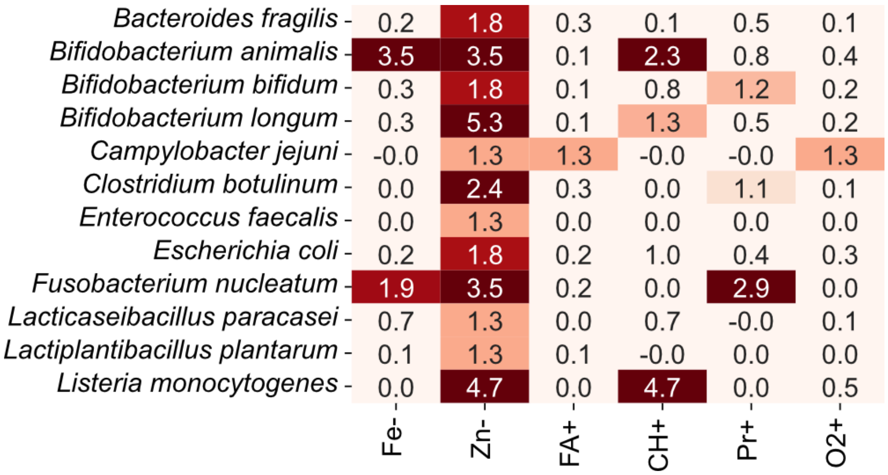

**Figure S9. Species enrichment analysis from ME-model predictions of members that are at an advantage under the six cases of study compared to WD.** Enrichment p-values were calculated with the Mann-Whitney U-test, right- and left-tailed for advantage and disadvantage, respectively. The heatmap shows p-values (p) as -log10 p, so significant enrichment is determined with $p<0.05$, equivalent to -log10 $p>1.3$.

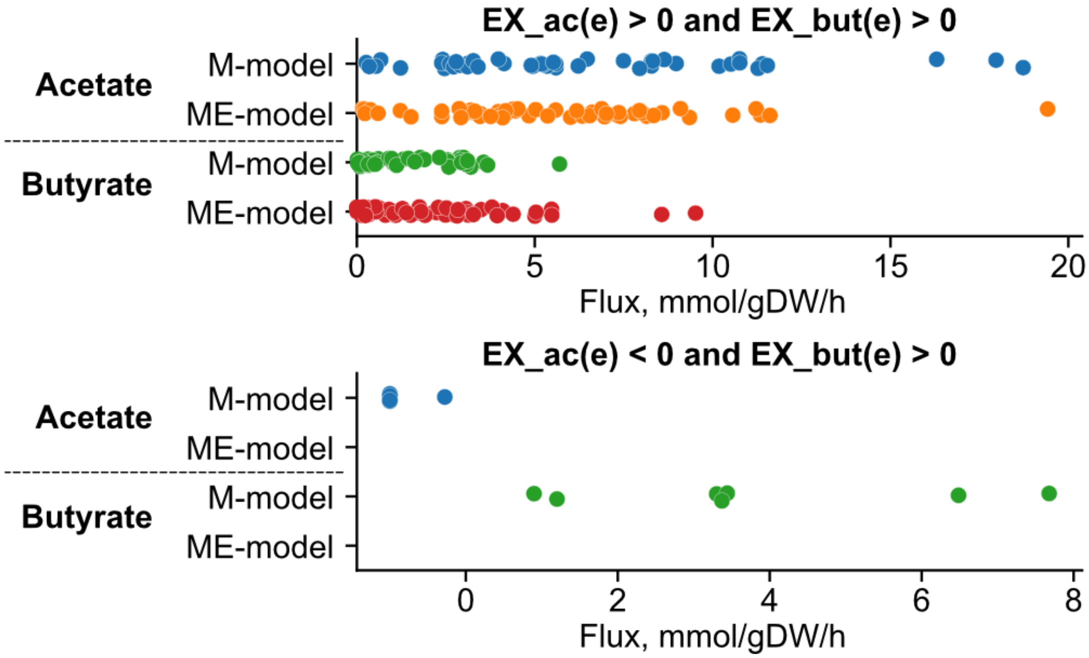

**Figure S10. Distribution of acetate and butyrate fluxes under optimality in Western Diet.** Fluxes for acetate and butyrate secretion (upper panel) and acetate uptake (bottom panel) for M- and ME-models. Dots represent the flux in mmol/gDW/h for acetate in M-models (blue), acetate in ME-models (orange), butyrate in M-models (green), and butyrate in ME-models (red). Labels show the number of models that predict the condition.

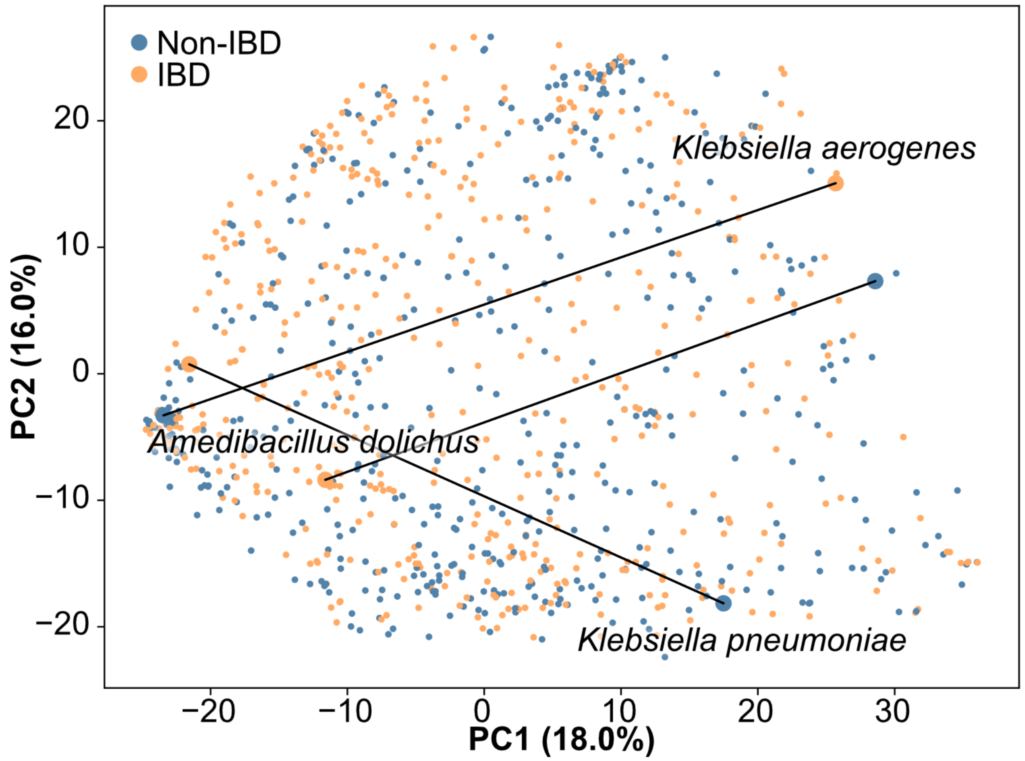

**Figure S11. PCA of the metabolic fluxes predicted by 495 cME-models in IBD and Non-IBD.** The three strains showing the maximum variation (the largest Euclidean projected distance on PC1 and PC2) are highlighted to exemplify the metabolic variation of individual organisms across the two conditions.

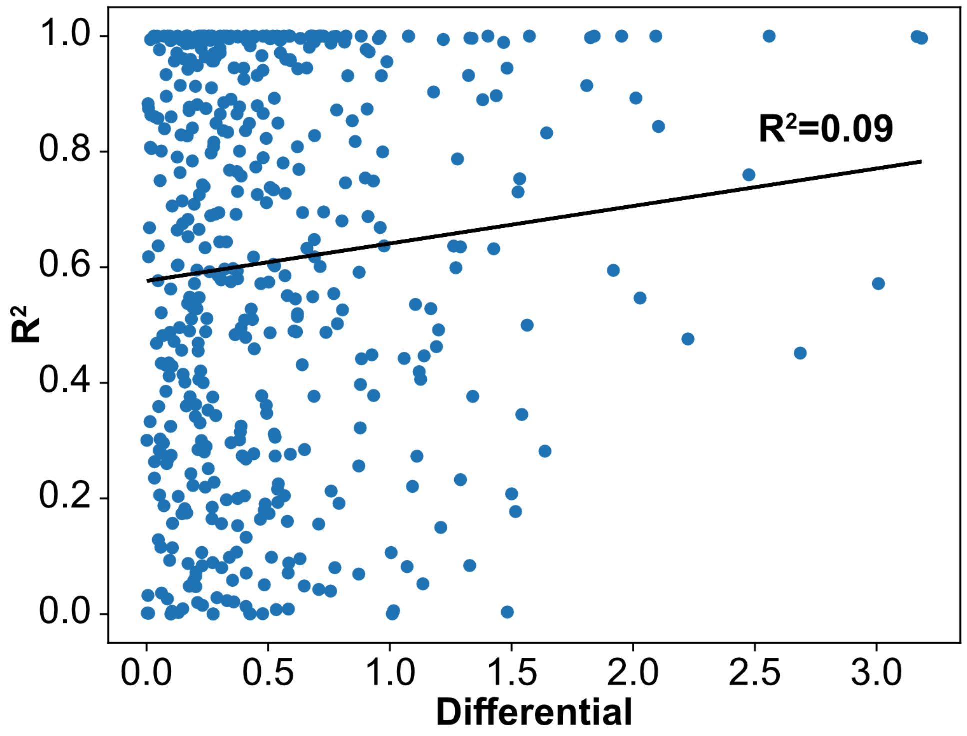

**Figure S12. Correlation between metabolic variability in IBD vs. Non-IBD ($R^2$) and absolute abundance differentials from metagenomics data.** No significant correlation between these two variables was found, indicating that metagenomics-inferred microbial abundances are not a proxy for metabolic activity in the gut.

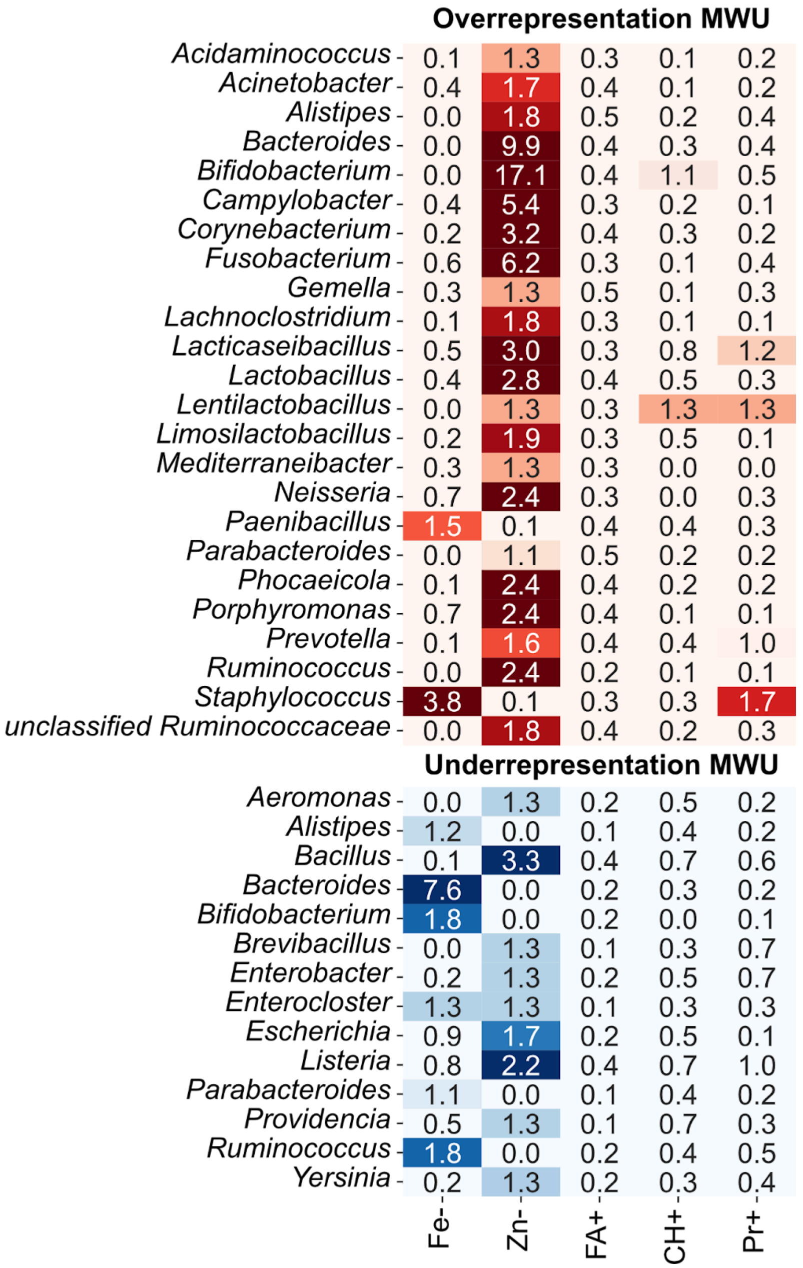

**Figure S13. Multi-omics-integrated predictions of dietary effects in an IBD gut.** Genus enrichment analysis of members that are at an advantage (d) or disadvantage (e) in a certain diet compared to WD. Enrichment p-values were calculated with the Mann-Whitney U-test, right- and left-tailed for advantage and disadvantage, respectively. The heatmap shows p-values (p) as -log10 p. Significant enrichment is determined with p<0.05, equivalent to -log10 p>1.3.

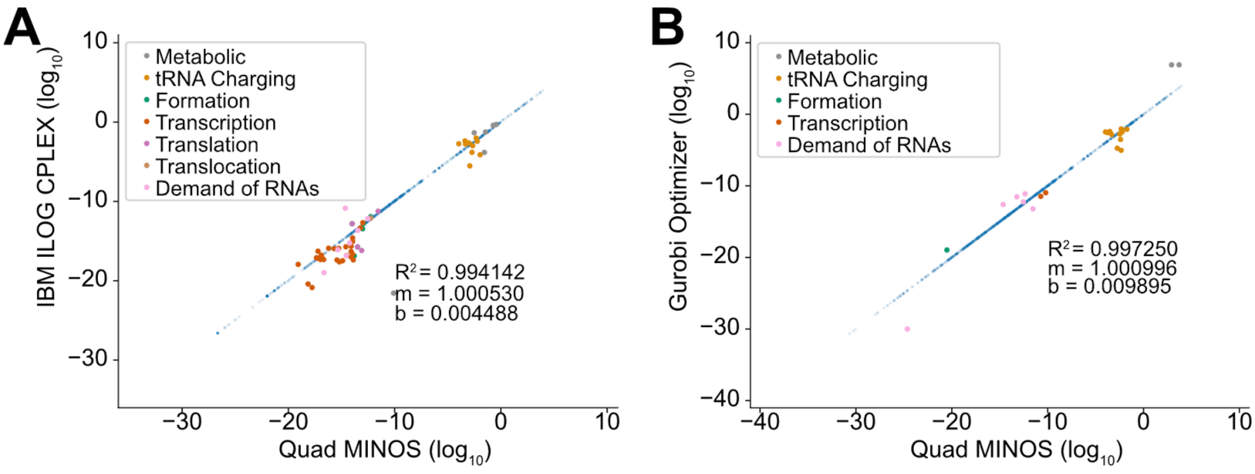

**Figure S14. Comparison of predicted fluxes by *i*Eco1689-ME with different solvers. (A)** Comparison of QMINOS and CPLEX. **(B)** Comparison of QMINOS and Gurobi.

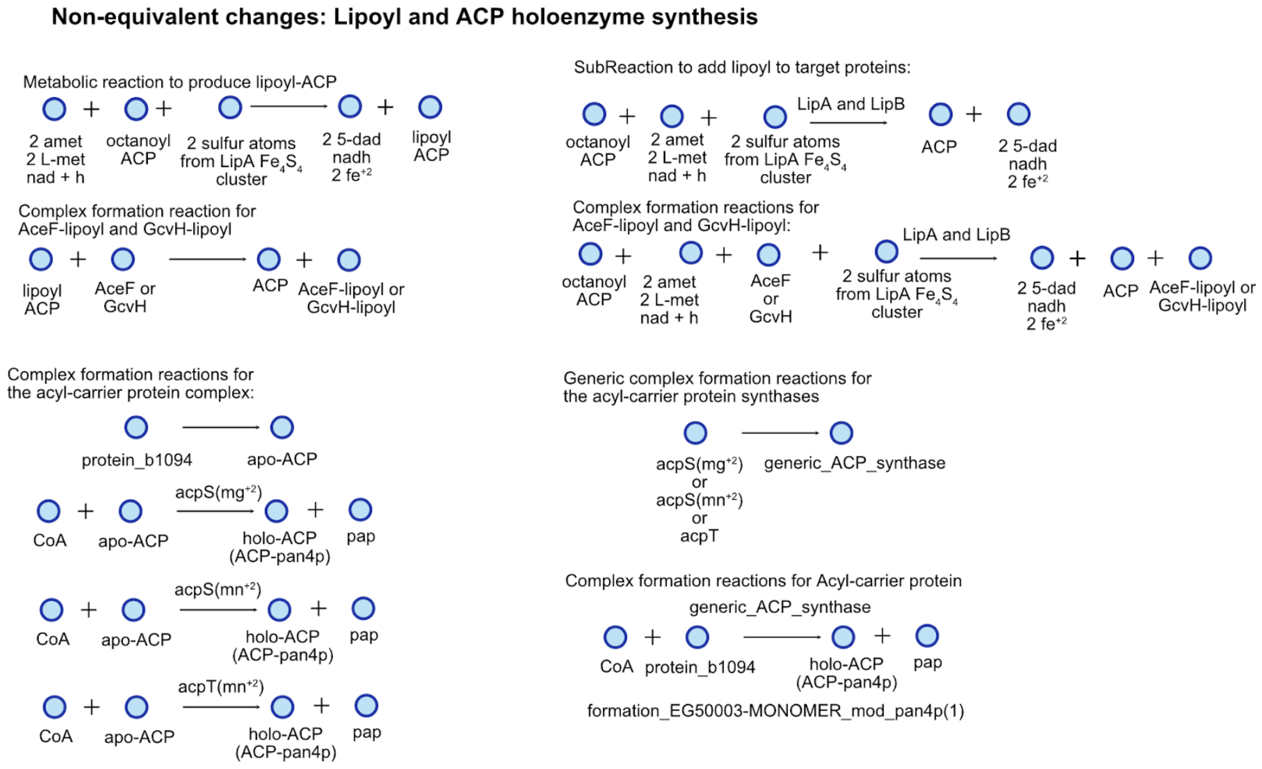

**Figure S15. Refactored reactions in the biosynthesis of lipoyl and holo-acyl carrier protein.** The implemented non-equivalent changes in coralME determine complex relationships to compare fluxes. The comparable flux for the lipoate *de novo* biosynthesis in COBRAme (top-left) is equivalent to the sum of two formation reactions in coralME catalyzed by LipA and LipB (top-right). In the case of pantetheine 4'-phosphate (pan4p), the formation of the holo-acyl carrier protein was refactored from two reaction steps (formation of apo-ACP and modification with pan4p) into one reaction in coralME, where there is no apo-ACP intermediate.

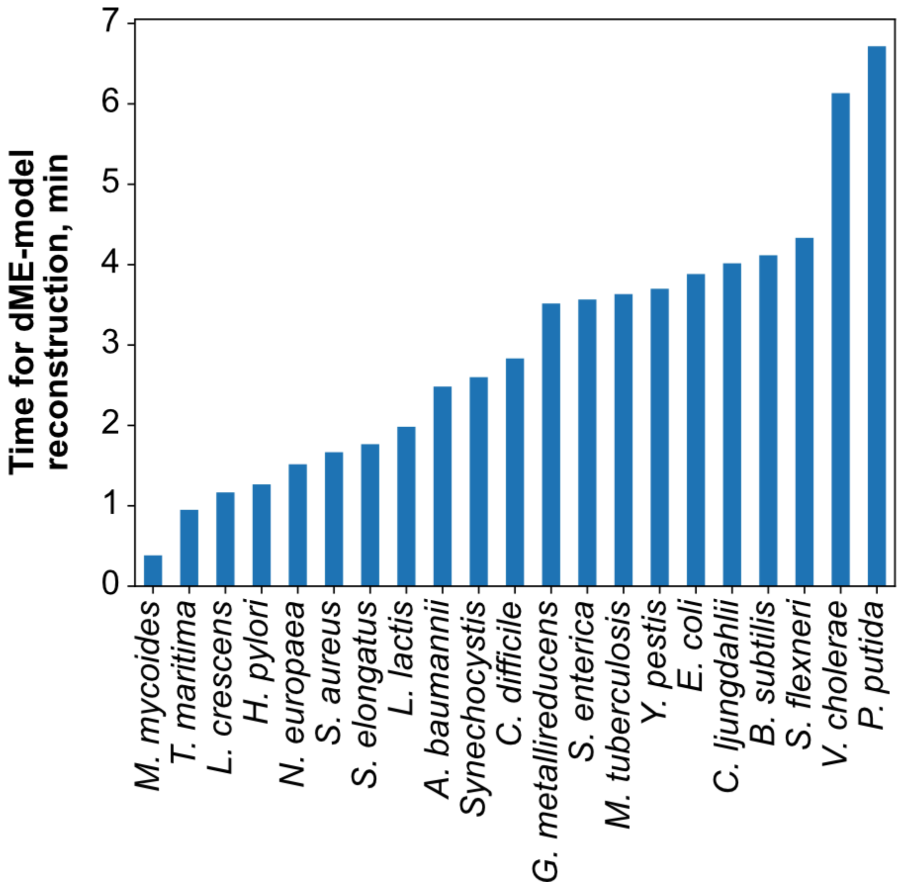

**Figure S16. Times for dME-model reconstruction of all 21 included microorganisms in minutes.** Reconstruction times were recorded using the computer specifications provided in Methods, and do not include the time to perform BLASTp and gap-finding.

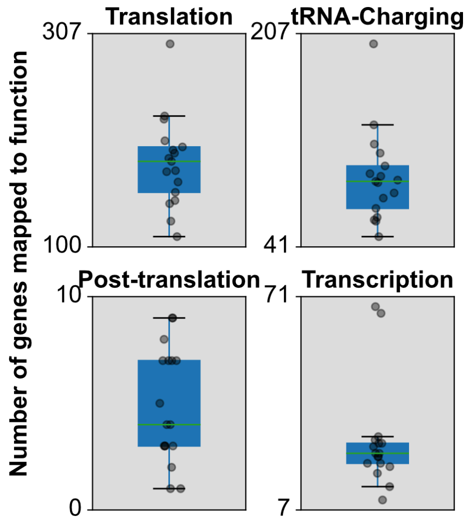

**Figure S17. Distribution of mapped gene functions to each expression category.** Boxplots show the distribution of gene functions mapped to translation, tRNA-charging, post-translational modifications, and transcription across the 21 dME-models.

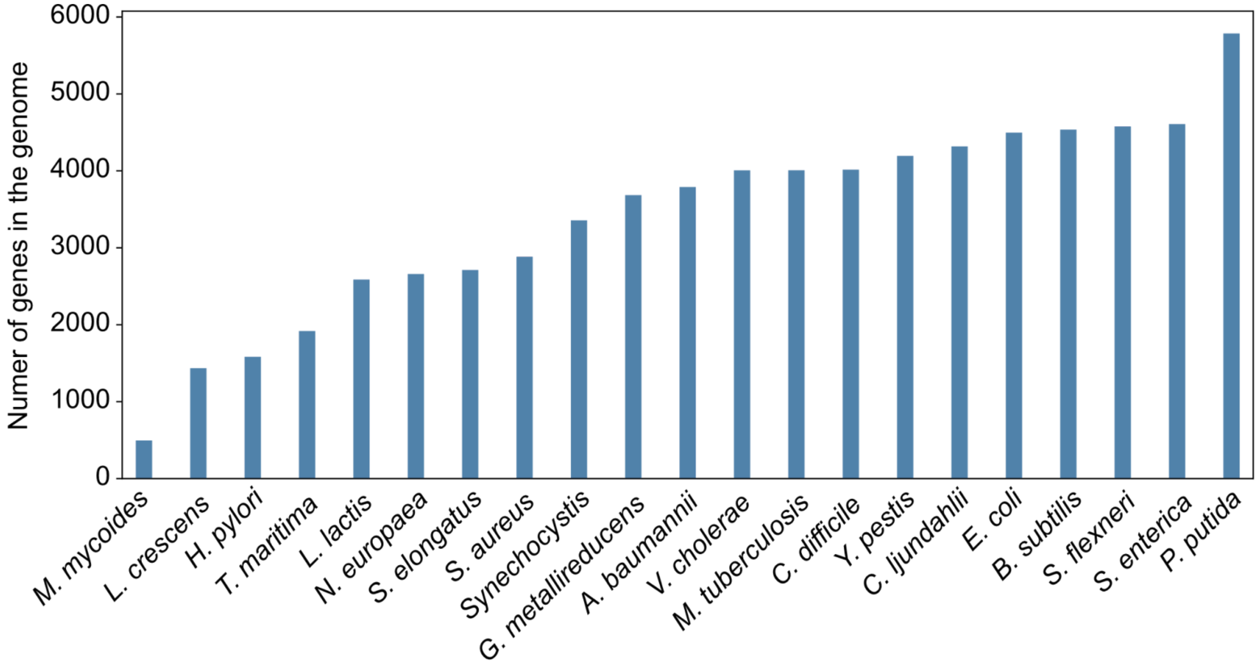

**Figure S18. Number of total genes in all genomes included in this work.** Gene numbers include all features marked as "gene" in the genome GenBank file.

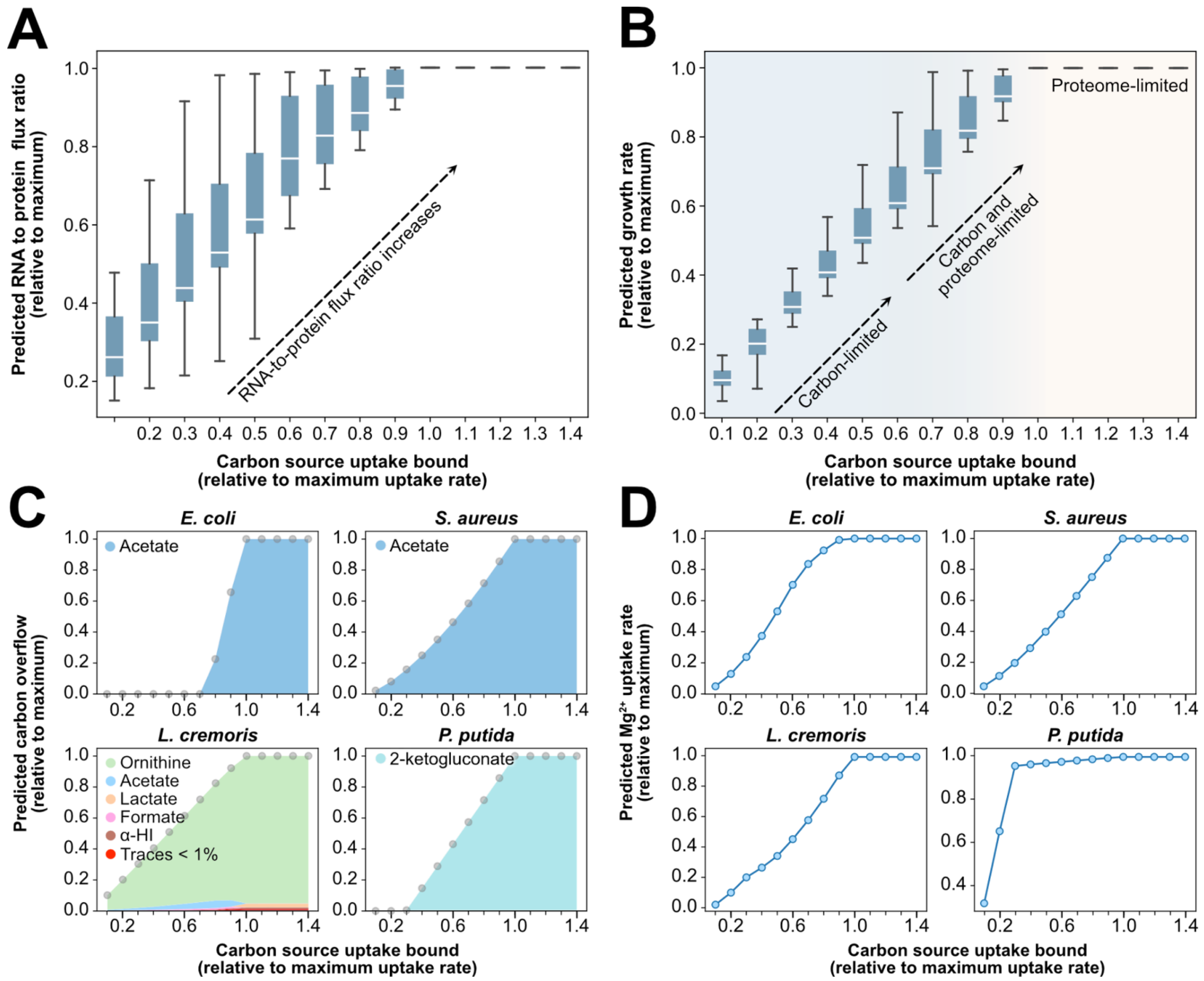

**Figure S19. Prediction of RNA-to-protein synthesis flux ratio, proteome limitation of growth, carbon overflow, and cofactor utilization by the 21 dME-models. (A)** Prediction of RNA to protein ratios at different intake flux of carbon sources. **(B)** Prediction of proteome limitation by dME-models at increasing carbon source uptake reaction bounds. **(C)** Prediction of carbon overflow for *E. coli*, *P. putida*, *L. cremoris*, and *S. aureus* dME-models at increasing carbon source uptake reaction bounds. **(D)** Prediction of magnesium uptake rates in dME-models as a result of different carbon source uptake reaction bounds. All data shown here is normalized to the maximum value in each dataset (see Methods). Absolute flux rate distributions are provided in Table S10.

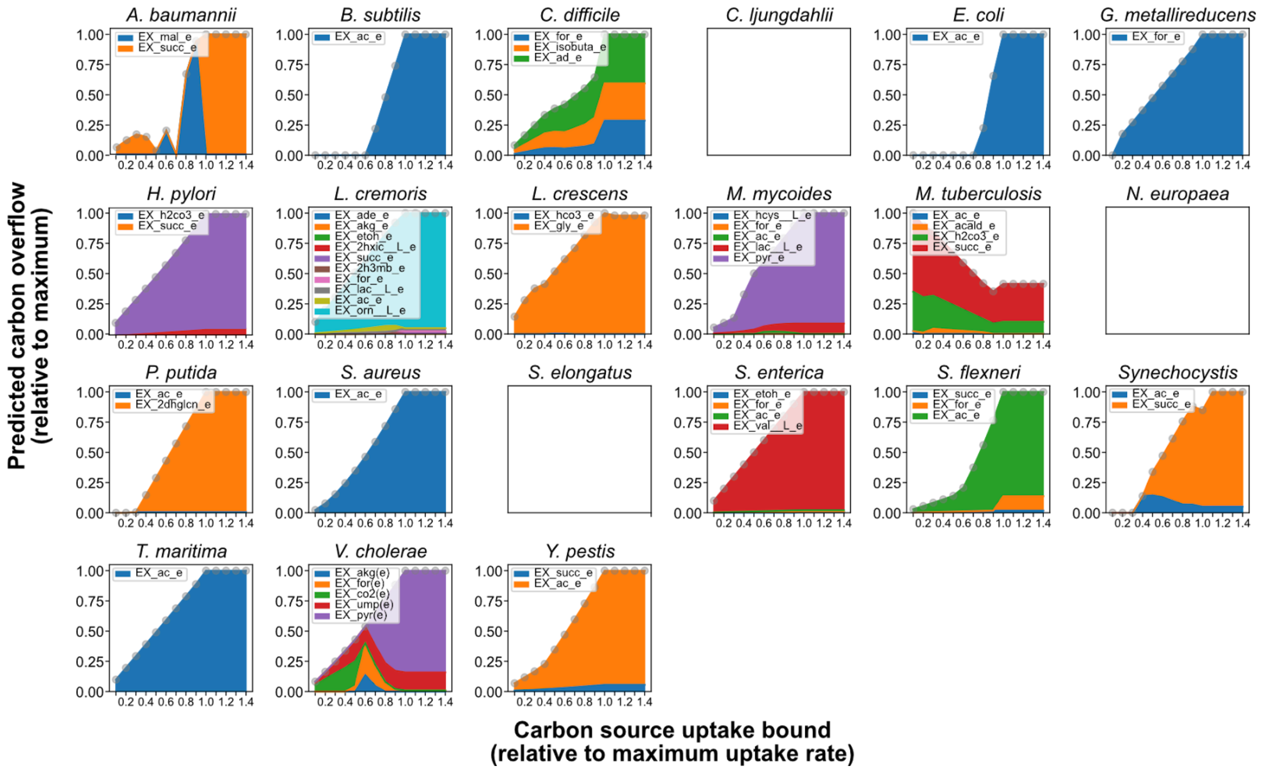

**Figure S20. Predicted overflow metabolism by dME-models.** Carbon overflow was predicted by increasing the carbon source(s) uptake bound, and the flux rates were normalized to the maximum of the series.

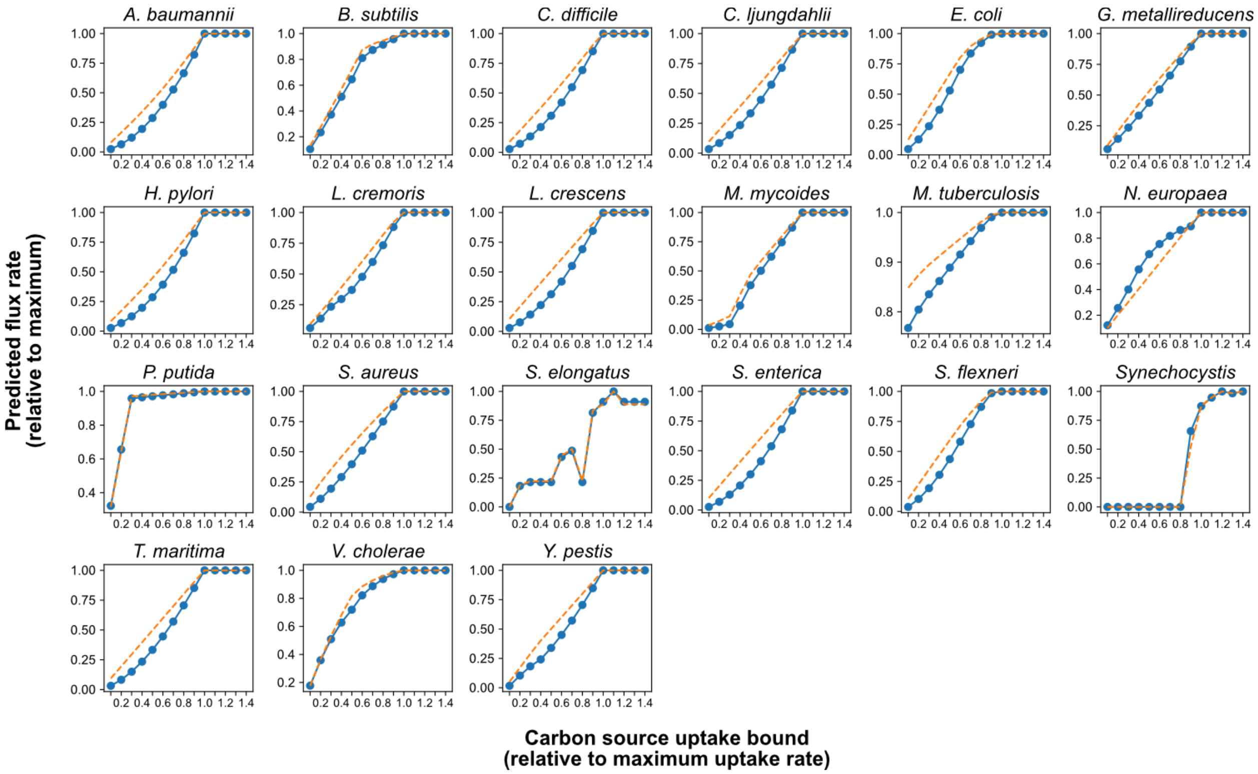

**Figure S21. Predicted magnesium requirement with growth rate by the dME-models.** Magnesium uptake was predicted by increasing the carbon source(s) uptake bound, and the flux rates were normalized to the maximum of the series.

**Table S1 (separate file). Top 10 metabolite fluxes predicted as overflow for each of the seven diets.** Values represent the ten most secreted metabolites per ME-model and condition. The metabolites were sorted in decreasing order based on the average predicted flux.

**Table S2 (separate file). Number of M- and ME-models predicting secretion or uptake of overflow metabolites under Western Diet.** A complete breakdown of the number of M- and ME-models that predict to consume or secrete one of the 16 metabolites identified as overflow metabolites, including butyrate. Fluxes were rounded up to six decimal places to avoid numerical artifacts.

**Table S3 (separate file). Predicted short-chain fatty acid overflows for each of the seven diets.** Complete overflow predictions for metabolites classified as short-chain fatty acids (acetate, formate, propionate, butyrate, D-lactate, L-lactate, and 2-oxobutanoate). Metabolites were sorted in decreasing order based on the average predicted flux.

**Table S4 (separate file). Top categories of metabolites predicted as overflows for each of the seven diets.** Aggregated fluxes for metabolites classified as short-chain fatty acid, amino acid, nucleoside, alcohol, dicarboxylic acid, amino acid derivative, vitamin, aldehyde, organic amine, branched fatty acid, sugar, organic acid, long-chain fatty acid, sterol, or gas. Categories were sorted in decreasing order (from left to right) based on the average predicted flux.

**Table S5 (separate file). Abundance differential analysis of metagenomics data using Songbird.** This file includes a complete breakdown of Songbird's output values, including predicted effect sizes for microbial abundance and sample as covariates and the intercept.

**Table S6 (separate file). Estimated contributions to change in secretion, consumption, and net rate of change of genera.** Values shown in this file are the final product of individual metabolic fluxes and their microbial abundance.

**Table S7. Comparative table of changes between COBRAme and coralME.** Code changes and manual curation are reflected in properties (e.g., maximum growth rate), parameter values (e.g., GC fraction), or processes (e.g., biosynthesis of cofactors) of the resulting ME-model.

| Model property, parameter, or process | *i*JL1678b-ME[2] | *i*Eco1689-ME (This work) |
|---|---|---|
| Maximum growth rate | 0.833297 1/h | 0.818627 1/h (-1.79%) |
| Genes, Metabolites, and Reactions | 1678, 7031, 12655 | 1689 (+0.66%), 7193 (+2.30%), 14956 (+18.18%) |
| GC fraction | Parameter set by the user of COBRAme: ~0.507896997 | Automated calculation from the GenBank file using BioPython:[3] ~0.507223292 (-0.13%). The GC fraction affects the stoichiometry of the DNA replication reaction and its bounds. |
| Estimation of the turnover rates (*keffs*) | Parameters set by the user of COBRAme. | Automated calculation based on the molecular formula of the complex. If the user provides *keffs*, they are set instead of the estimated *keff*. |
| Elemental contribution of subreactions | Parameter set by the user of COBRAme. | Automated calculation based on the stoichiometry and molecular formulas of metabolites. |
| Lipid modifications | Parameter set by the user of COBRAme. | Lipids involved in lipid modifications are detected automatically from the M-model using a regular expression. The user can modify the list of lipids manually. |
| Frameshifts | Parameter set by the user of COBRAme. The parameter is needed to get the correct protein sequence from the nucleotide sequence. | Automated detection using BioPython.[3] For instance, the RNA_b2891 component now shows two start and two end positions due to a programmed frameshift. Translation of the nucleotide sequence is automatically handled using BioPython and the translation code set in the GenBank file. |
| Pseudogenes | Pseudogenes are not added during the reconstruction. A parameter can be to add pseudogenes. | Pseudogenes can be added during reconstruction, especially if they are associated with M-reactions. The amino acid sequence is determined using BioPython3 using the known start and end positions. The nucleotide sequence is truncated to match three nucleotides per codon, and the last codon is interpreted as a stop codon. |
| Inactive reactions | Reactions with LB=0 and UB=0 are not added during the reconstruction. | Reactions with LB=0 and UB=0 are added during the reconstruction and not removed during prune. It allows the modification of bounds without reconstruction of the ME-model, and the set of effective turnover rates (*keffs*) if the user defined them for the reverse reaction. |

| | | |
|---|---|---|
| Translation of proteins | COBRAme provides only one translation table (translation table 11) | Automated detection of the translation table using BioPython for each gene in the GenBank file. The default value is 11 (Bacterial, Archaeal, and Plant Plastid Code). The detection of the code allows the correct modeling of mycoplasmas and spiroplasmas (translation table 4; see https://www.ncbi.nlm.nih.gov/Taxonomy/Utils/wprintgc.cgi#SG4 for more information). |
| Excision machinery | RNase T is counted twice in the excision machinery component. | RNase T is counted only one time in the excision machinery as a precursor of the *generic_RNase* component. |
| Sec-tRNA precursor | Cys-tRNA$^{Sec}$ was set as the precursor. | Ser-tRNA$^{Sec}$ is the correct precursor in bacteria, archaea, and eukaryotes.[4] In bacteria, SelA adds selenium to the mischarged serine in selC from selenophosphate, and SelB is a specialized transcription elongation factor needed for the incorporation of Sec-tRNA$^{Sec}$ during translation. This change impacts the stoichiometry of the tRNA charging and other reactions and the molecular formula of selenoproteins (deletion of one sulfur atom and addition of an oxygen atom compared to previous formulas). |
| Lipoyl biosynthesis | Octanoyl-ACP is converted into lipoyl-ACP. The lipoyl moiety is then transferred from ACP into acceptor enzymes. | Octanoyl from ACP is loaded into acceptor enzymes by LipB. Then, it is modified into lipoyl by LipA. The biosynthesis of the cofactor is added as a *SubReaction* associated with both enzymes. The comparable flux of 9.85e-7 mmol/gDW/h from *i*JL1678b-ME is split into 3.43e-7 to the formation of the oxoglutarate dehydrogenase complex and 6.43e-7 to the formation of the pyruvate dehydrogenase complex. |
| Regeneration of the LipA accessory iron-sulfur cluster. | The regeneration of the iron-sulfur cluster proceeds by transferring a free $Fe_2S_2$ cluster to LipA. | The regeneration of the LipA accessory iron-sulfur cluster proceeds by transferring the $Fe_4S_4$ cluster from the NfuA dimer to LipA.[5] The comparable flux varies from 9.8552e-7 (*i*JL1678b-ME) to 9.8627e-7 mmol/gDW/h in *i*Eco1689-ME. |
| Dipyrromethane (DPM) biosynthesis | DPM is imported using an orphan reaction. Import flux was 10.0e-11 mmol/gDW/h. | DPM is biosynthesized by the porphobilinogen deaminase apoenzyme,[6] generating the holoenzyme and hydroxymethylbilane. Biosynthesis flux was 9.56e-11 mmol/gDW/h. |
| NiFe cofactor biosynthesis | No reactions modeling the biosynthesis of the NiFe cofactor from small molecules. | The synthesis of the NiFe cofactor[7] was modeled step by step. The assembly of a Ni-free precursor occurs using the HypCDE complex as a scaffold. The precursor is then transferred to the hydrogenase complex. A final step models the transference of a nickel atom from a carrier protein. The loading of carbon dioxide and iron to the HypC dimer was modeled as a non-enzymatic reaction. In some bacteria, the |

| | | |
|---|---|---|
| | | HypX has been associated with the transference of carbon monoxide from 10-formyltetrahydrofolate to the HypC dimer.<br><br>There is no comparable flux due to the aerobic simulation condition that makes unnecessary the formate hydrogenlyase activity of the hydrogenase complex. |
| Holo acyl carrier protein (Holo-ACP) biosynthesis | The pantetheine 4'-phosphate in acyl-carrier proteins is catalyzed by one of three enzymes. | The pantetheine 4'-phosphate is included as a *SubReaction* in the formation of the acyl-carrier protein complex. The comparable flux changed from 3.07e-6 to 2.97e-6 mmol/gDW/h. |
| RNA modifications | No enzyme was assigned to orphan modifications. | The *CPLX_dummy* component is assigned to orphan tRNA and rRNA modifications, accounting for an estimated metabolic burden. After manual curation, only two RNA modifications are associated with *CPLX_dummy*. The nucleotide and amino acid composition of the "CPLX_dummy" component remains the same compared to *i*JL1678b-ME. |
| Processing of transcriptional units (TU) containing tRNAs and/or rRNAs | A TU is processed if there is at least one non-mRNA gene in the TU.<br><br>The number of cuts is proportional to the number of genes (of any type) before pruning. Some TU shows a negative number of excised nucleotides due to overlapping ncRNAs. | Only operons containing tRNA(s) or rRNA(s) are processed by RNases. In consequence, the TUs containing the RNase P catalytic RNA component (RnpB gene) are not processed.<br><br>The number of cuts is proportional to the number of rRNA(s) and tRNA(s) in the TU before pruning. The number of excised nucleotides is always positive. |
| Processing of protein complex from M-model Gene-Product-Reaction associations (GPRs) | Protein complexes and their associations with M-model reactions are set manually by the user of COBRAme. | The M-model GPRs are processed to identify all possible protein complex combinations. When a GPR results in a combinatorial explosion (> 100 combinations), it is simplified by using *generics* components. In brief, a recursive algorithm turns all *AND* rules into *complexes* and *OR* rules into *generics*. The procedure reduces the number of final formation reactions by orders of magnitude.<br><br>The user can set *generics* components derived from |

| | | |
|---|---|---|
| | | complexes manually. In this case, new complexes are created and associated with reactions instead of the original complexes. For example, *CPLX_acpP_activation-0* derives from the *generic_acp_synthase* component, and it is the correct complex associated with the formation of the holo-ACP complex (Supplementary Fig S10). |

**Table S8. Manual curation of the reference ME-model.** A detailed description of new, corrected, removed, replaced, and renamed ME-components.

| | |
|---|---|
| New genes and gene products | tsaA (b0195): tRNA $m^6t^6A37$ methyltransferase.<br><br>tsaB (b1807): $N^6$-L-threonylcarbamoyladenine synthase, subunit B<br><br>tapT (b2583): tRNA 3-amino-3-carboxypropyltransferase.<br><br>hypF (b2712): carbamoyl--[HypE] ligase<br><br>hypA (b2726): hydrogenase 3 nickel incorporation protein HypA<br><br>hypB (b2727): hydrogenase isozymes nickel incorporation protein HypB<br><br>hypC (b2728): hydrogenase maturation factor HypC<br><br>hypD (b2729): $Fe\text{-}(CN)_2CO$ cofactor assembly scaffold protein HypD<br><br>hypE (b2730): carbamoyl dehydratase HypE<br><br>tsaD (b3064): $N^6$-L-threonylcarbamoyladenine synthase, subunit D<br><br>nfuA (b3414): possible scaffold/chaperone for damaged Fe/S proteins[5]<br><br>rsmJ (b3497): 16S rRNA $m^2G1516$ methyltransferase.<br><br>tsaE (b4168): $N^6$-L-threonylcarbamoyladenine synthase, subunit E<br><br>rimI (b4373): [ribosomal protein S18]-alanine N-acetyltransferase<br><br>New genes imply the addition of new transcriptional units (TUs) or the addition of gene(s) to existing TUs. Also, the addition of new transcription and translation reactions, and new formation reactions for complexes and modified complexes. All the new gene products are cytosolic (experimental or inferred computationally). |
| New complexes and modified complexes | *RsmJ_mono*: 16S rRNA (guanine1516-N2)-methyltransferase monomer<br><br>*TapT_mono*: tRNA 3-amino-3-carboxypropyltransferase monomer<br><br>*TsaA_mono*: tRNA m6t6A37 methyltransferase monomer<br><br>*TsaB_mono*: N6-L-threonylcarbamoyladenine synthase, TsaB subunit monomer<br><br>*TsaD_mono*: N6-L-threonylcarbamoyladenine synthase, TsaD subunit monomer<br><br>*TsaE_mono*: N6-L-threonylcarbamoyladenine synthase, TsaE subunit monomer<br><br>*RimI_mono*: ribosomal-protein-S18-alanine N-acetyltransferase monomer |

| | |
|---|---|
| | *NfuA_dim*: iron-sulfur cluster carrier protein NfuA dimer<br><br>*HypC_dim*: hydrogenase maturation factor HypC dimer<br><br>*hypD_mono*: Fe-(CN)2CO cofactor assembly scaffold protein HypD monomer<br><br>*hypCD*: HypC-HypD heterodimer<br><br>*hypE*: carbamoyl dehydratase HypE monomer<br><br>*hypCDE*: HypC-HypD-HypE heterotrimer<br><br>*hypF*: carbamoyl--[HypE] ligase monomer<br><br>*hypA*: hydrogenase 3 nickel incorporation protein HypA monomer<br><br>*hypB*: hydrogenase isozymes nickel incorporation protein HypB monomer<br><br>*hypAB*: HypA-HypB heterodimer<br><br>*NfuA_dim_mod_4fe4s(1)*: Active NfuA dimer<br><br>*HypC_dim_mod_fe2(1)_mod_co2(1)*: First intermediate in the synthesis of the NiFe cofactor.<br><br>*hypD_mono_mod_4fe4s(1)*: Replaces one HypC after coordination of iron and carbon monoxide in the HypC dimer.<br><br>*hypCD_mod_4fe4s(1)_mod_FeCO(1)*: Second intermediate in the synthesis of the NiFe cofactor.<br><br>*hypE_mod_cn(1)*: cyanide carrier in the synthesis of the NiFe cofactor.<br><br>*hypCDE_mod_4fe4s(1)_mod_FeCO[CN]2(1)*: Third intermediate in the synthesis of the NiFe cofactor. The cofactor is transferred to the acceptor enzyme.<br><br>*CPLX0-250_FORMATEDEHYDROGH-MONOMER_mod_FeCO[CN]2(1)*: First intermediate in the synthesis of the active CPLX0-250 complex.<br><br>*CPLX0-250_FORMATEDEHYDROGH-MONOMER_mod_NiFe_cofactor(1)*: Second intermediate in the synthesis of the active CPLX0-250 complex. |
| New associated functions | RlmN (b2517) is also a tRNA $m^2$A37 methyltransferase.<br><br>TrmJ (b2532) is also an Um32 methyltransferase. |
| Corrected stoichiometry of reactions and subreactions | 5-Formyltetrahydrofolate and 5,6,7,8-Tetrahydrofolate were replaced by 5,10-Methylenetetrahydrofolate and 7,8-Dihydrofolate, respectively. More information here https://biocyc.org/ECOLI/NEW-IMAGE?object=PWY-7892 |

| | |
|---|---|
| | Carrier activity was corrected. Previously, the carrier activity was misidentified using the *StoichiometricData* for the forward reaction in reversible reactions, associating a dilution coefficient to a product rather than the substrate.<br><br>Stoichiometry of the *lipoyl_denovo* subreaction was corrected to release iron and hydrogen sulfide, destroying the accessory iron-sulfur cluster of LipA. |
| Removed ME-components | NHFRBO: The reaction was corrected manually in *i*JL1678b-ME to add NorW (b2711) to the enzyme list and also to the subreaction list. NHFRBO is a three-step electron transfer reaction: 1) from NADH to NorW, 2) from NorW to NorV, and 3) from NorV to nitric oxide.<br><br>RNases m16, m23, and m5: Sink reactions were added in *i*JL1678b-ME to model the autocatalytic RNase activity of the pre-5S, pre-16S, and pre-23S rRNAs[7]. However, they do not have a formula or molecular weight, therefore no metabolic requirements for their biogenesis.<br><br>No transport, exchange, or sink reactions for 2,3-biphosphoglycerate, cesium ion, thallium ion, dipyrromethane, and pyrroloquinoline quinone. The *E. coli* bacterium cannot synthesize pyrroloquinoline quinone. To compare fluxes, a sink reaction for the lithium ion was added manually.<br><br>EG11910-MONOMER_dimer and EG11911-MONOMER: The complex *EG11910-MONOMER_dimer_EG11911-MONOMER* is synthesized directly from two *protein_b3951* and one *protein_b3952* components. |
| Replaced ME-components | A *MetabolicReaction* to synthesize the lipoyl prosthetic group was replaced with two *SubReactions*: *lipoyl_scavenging* (from lipoate) and *lipoyl_denovo* (from octanoyl-ACP).<br><br>FMETTRS is detected from the M-model and, if it exists, replaced with a *SubReaction*. Otherwise, FMETTRS is created as a *SubReaction*.<br><br>ATPM is detected from the M-model and, if it exists, replaced with a *SummaryVariable*. Otherwise, ATPM is created as a *SummaryVariable*. |
| Renamed ME-components | TranslocationReactions IDs include a suffix indicating the final subcellular location of the protein. It allows the translocation of the same protein to different subcellular compartments. |

| | |
|---|---|
| | Notation of modified complexes: mod_#:metaboliteID → mod_metaboliteID(#), where the # represents the stoichiometry of the metabolite in the complex.<br><br>NiFeCoCN2 was renamed to NiFe_cofactor, and yrdC was renamed to tsaC.<br><br>The Braun lipoprotein ID (from b1677) was replaced with EG10544-MONOMER. |
| Other changes | The formulas for the amp, malonyl, and biotinyl prosthetic groups were corrected. |

**Table S9. Sources of M-models and genomes of the 21 bacteria with manually curated M-models.** Manually curated M-models were retrieved from various sources, e.g. BiGG Database and individual publications. Here we provide the specific organism name, M-model identifier, genome identifier, and specific source from which they were retrieved.

| Organism | M-model | Genome NCBI ID | Source |
|---|---|---|---|
| *A. baumannii* | iCN718 | NC_010410.1 | BiGG |
| *B. subtilis* | iYO844 | AL009126.3 | BiGG |
| *C. difficile* | iCN900 | AM180355.1 | BiGG |
| *C. ljungdahlii* | iHN637 | NC_014328.1 | BiGG |
| *E. coli* | iJO1366 | NC_000913.3 | BiGG |
| *G. metallireducens* | iAF987 | CP000148.1 | BiGG |
| *H. pylori* | iIT341 | NC_000915.1 | BiGG |
| *L. cremoris* | iNF517 | NC_009004.1 | BiGG |
| *L. crescens* | L. crescens BT-1 | CP003789.1 | Ref.[8] |
| *M. mycoides* | MMSYN | CP016816.2 | Ref.[9] |
| *M. tuberculosis* | iEK1008 | NC_000962.3 | BiGG |
| *N. europaea* | iGC535 | NC_004757.1 | Ref.[11] |
| *P. putida* | iJN1463 | NC_002947.4 | BiGG |
| *S. aureus* | iSB619 | NC_002745.2 | BiGG |

| | | | |
|---|---|---|---|
| *S. elongatus* | iJB785 | NC_007604.1 | BiGG |
| *S. enterica* | STM_v1_0 | AE006468.1 | BiGG |
| *S. flexneri* | iS_1188 | AE014073.1 | BiGG |
| *Synechocystis sp.* | iJN678 | BA000022.2 | BiGG |
| *T. maritima* | iLJ478 | NC_000853.1 | BiGG |
| *V. cholerae* | iAM_Vc960 | AE003852.1 | Ref.[10] |
| *Y. pestis* | iPC815 | NC_003143.1 | BiGG |

**Table S10 (separate file). Complete simulated flux distribution of 21 ME-models from manually curated M-models under varying carbon levels.** Carbon uptake bounds were varied from 10% to 140% of the maximum calculated carbon source uptake rate in increments of 10%. This analysis was performed to show the self-limiting property of ME-models, which do not exceed uptakes and growth rates above proteome limitation.

### Document S1

Documentation of coralME

### coralME Documentation

*Release 1.0*

**Juan D. Tibocha Bonilla, Rodrigo Santibanez-Palominos**

**Apr 08, 2024**

**3 Indices and tables** **30**

**Index** **31**

### CHAPTER ONE

### DESCRIPTION

The **COmprehensive Reconstruction ALgorithm for ME-models (coralME)** is an automatic pipeline for the reconstruction of ME-models. coralME integrates existing ME-modeling packages COBRAme, ECOLIme, and solveME, generalizes their functions for implementation on any prokaryote, and processes readily available organism-specific inputs for the automatic generation of a working ME-model.

coralME has four main objectives:

1. **Synchronize** input files to remove contradictory entries.
2. **Complement** input files from homology with a template organism to complete the E-matrix.
3. **Build** a working ME-model
4. **Inform** the user about necessary steps to curate the ME-model.

This resource is intended to:

1. Describe basic inputs required for ME-model reconstruction.
2. Describe the architecture of coralME.
3. Demonstrate how to build a ME-model with coralME.
4. Describe how to perform manual curation guided by coralME's curation notes.

CHAPTER

TWO

### CONTENT

#### 2.1 Getting started

For details on inputs go to *Description of Inputs*.

For information about coralME architecture go to *Arquitecture of coralME*.

##### 2.1.1 Download files from BioCyc

BioCyc files are optional but useful, you should download them after having your gene id consistent M-model and genbank files.

The quickest way to do this is to copy one of the genes from the genbank file into the BioCyc search bar. Your organism should appear in the list if it is available in BioCyc.

To download:

###### Go to Tools>Special SmartTables

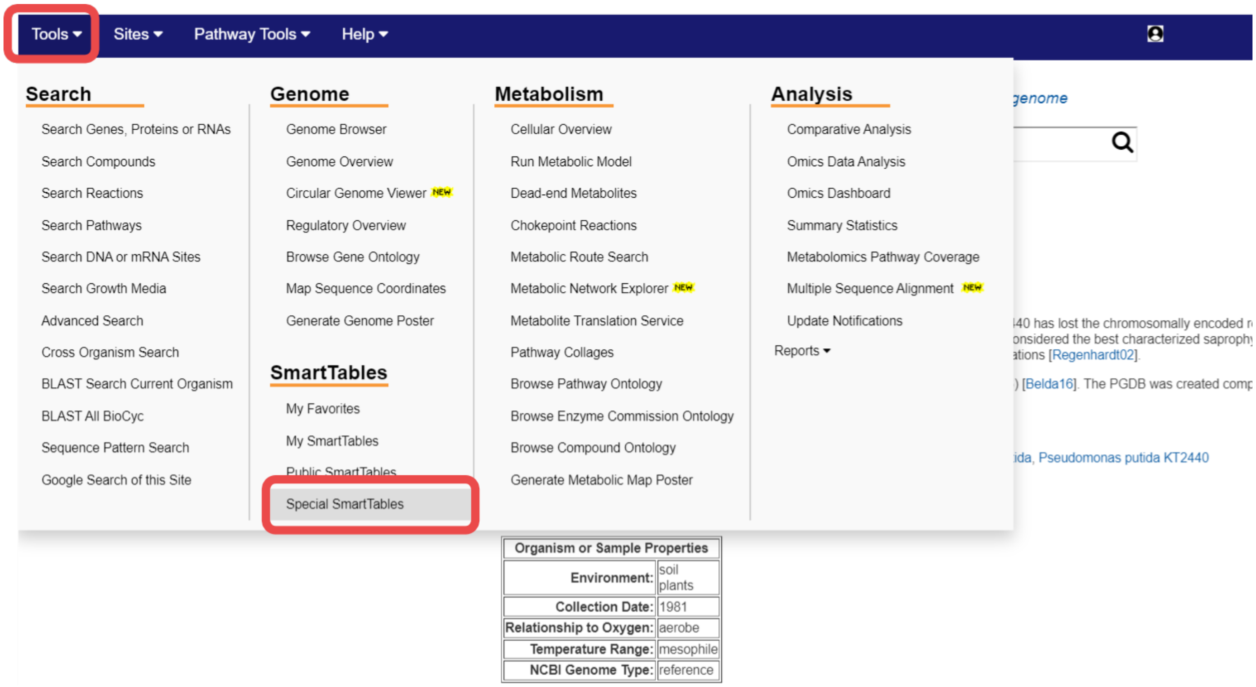

From here you can download the 5 optional BioCyc files for your organism.

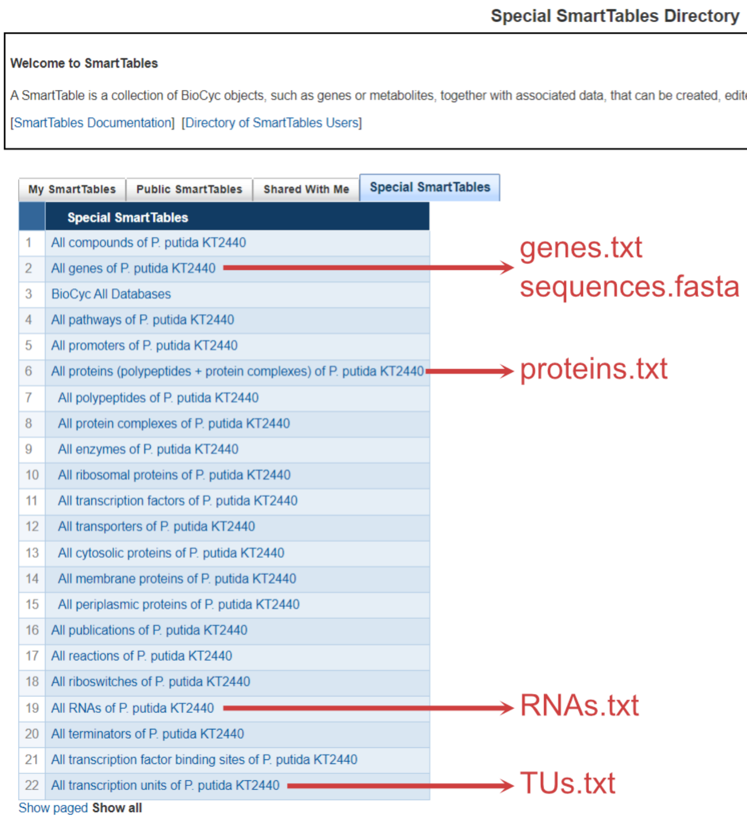

##### Download genes.txt and sequences.fasta

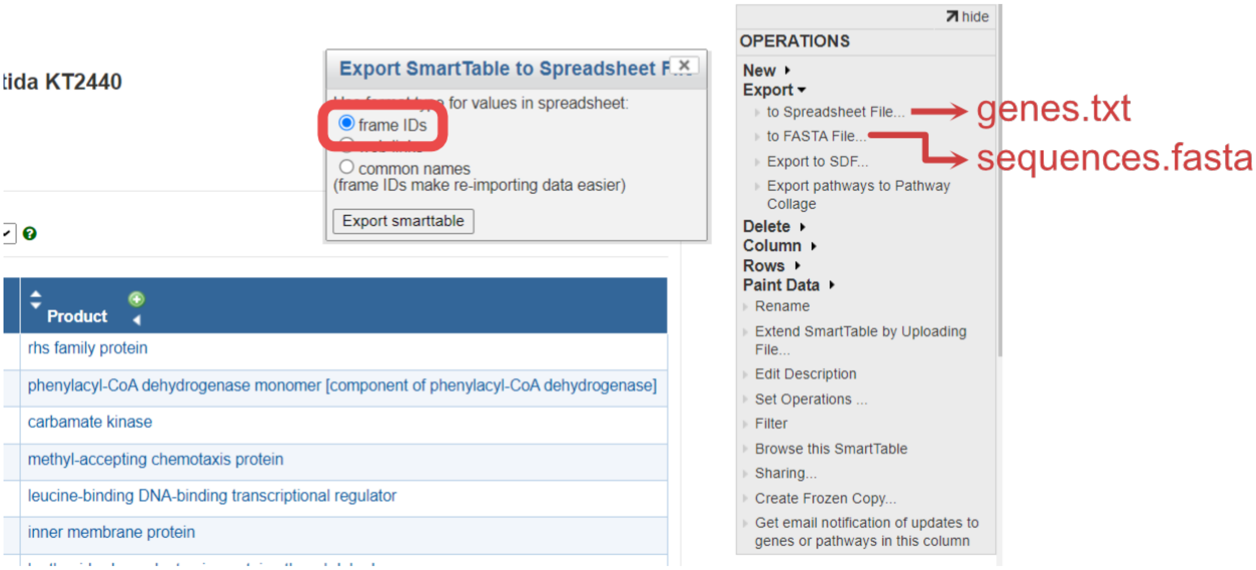

##### Download proteins.txt, RNAs.txt and TUs.txt

The same process of genes.txt applies to proteins.txt, RNAs.txt and TUs.txt.

Some columns must be added manually using BioCyc's dropdown lists **ADD PROPERTY COLUMN** and **ADD TRANSFORM COLUMN** within the SmartTable editing webpage.

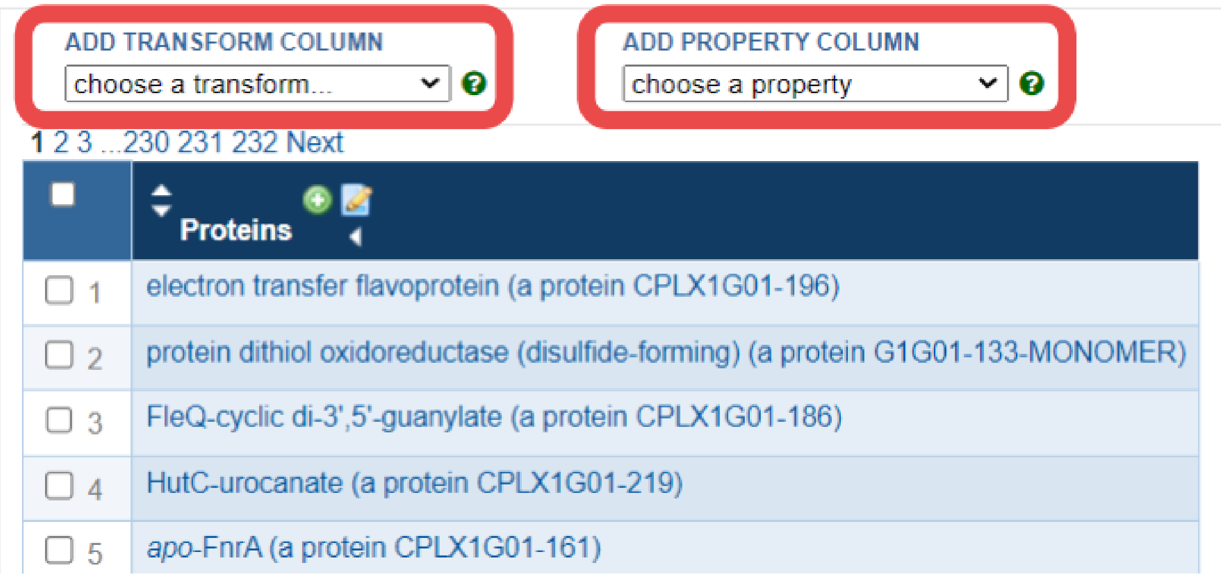

##### Download proteins.txt

The index (Proteins Complexes) is in the SmartTable by default, but you need to add the columns Common-Name, Genes of polypeptide, complex, or RNA, and Locations.

- Common-Name is available in the dropdown list **ADD PROPERTY COLUMN**
- Genes of polypeptide, complex, or RNA is available in the dropdown list **ADD TRANSFORM COLUMN**
- Locations is available in the dropdown list **ADD PROPERTY COLUMN**.

##### Download RNAs.txt

The index (All-tRNAs Misc-RNAs rRNAs) is in the SmartTable by default, but you need to add the columns Common-Name, and Gene.

- Common-Name is available in the dropdown list **ADD PROPERTY COLUMN**
- Gene is available in the dropdown list **ADD PROPERTY COLUMN**.

###### Download TUs.txt

The index Transcription-Units is in the SmartTable by default, but you need to add the columns Genes of transcription unit, and Direction.

- Genes of transcription unit is available in the dropdown list **ADD TRANSFORM COLUMN**
- Direction is available in the dropdown list **ADD PROPERTY COLUMN**.

##### 2.1.2 Initialize the folder for your organism

Copy your files to create your initial folder

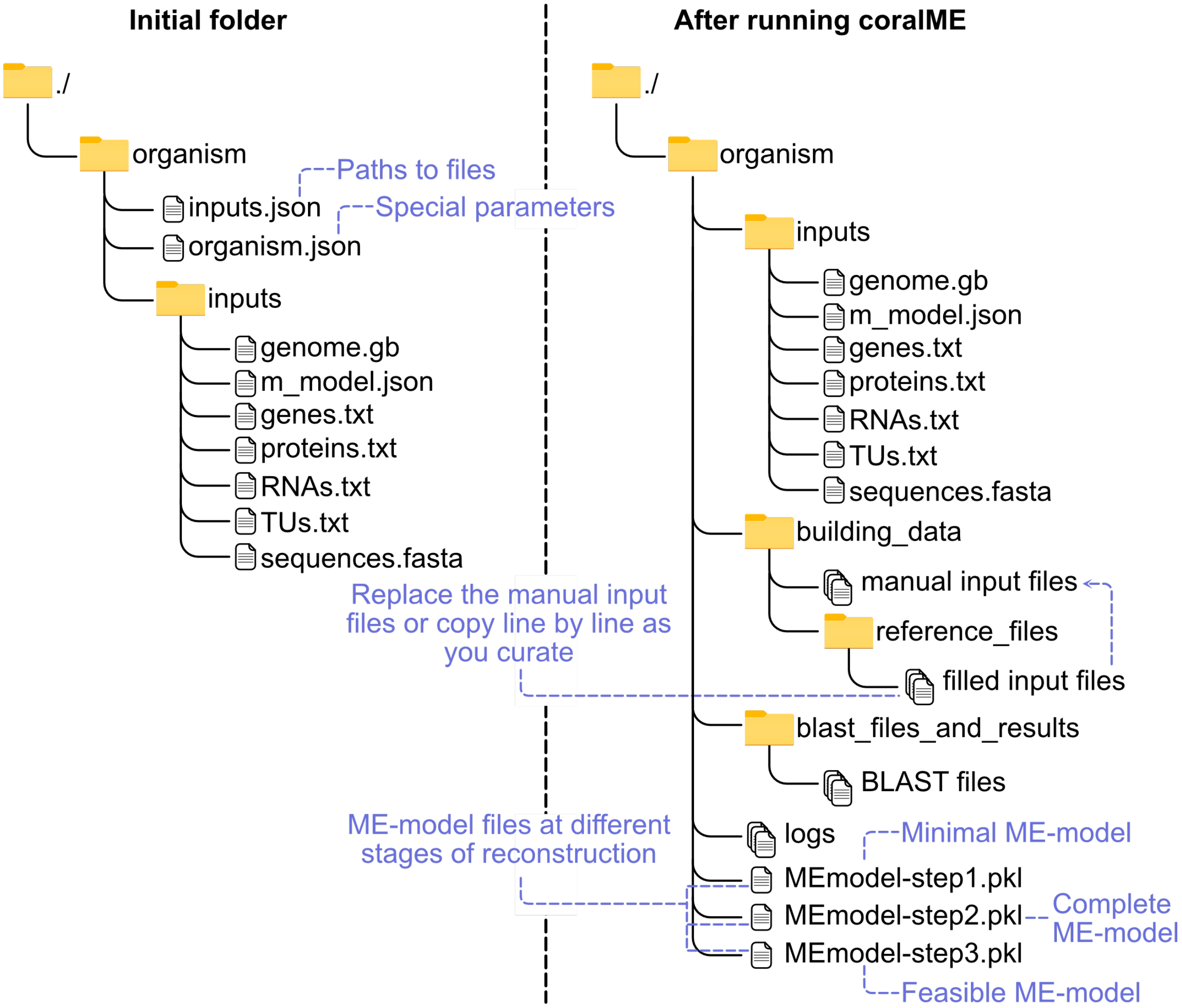

###### Define inputs in input.json.

See an example of input.json

###### Define parameters in organism.json.

See an example of organism.json

**Note**: You do not need to modify these parameters right away. But once you are at the curation stage you will have to ensure these parameters are applicable to your organism.

##### 2.1.3 Reconstruct with coralME

Here we show an example to reconstruct a dME-model of *B. subtilis*

###### Import packages

```
[ ]: from coralme.builder.main import MEBuilder
```

###### Define organism and inputs

```
[ ]: org = "./helper_files/tutorial/"
```

Load configuration files

```
[ ]: import os
     os.chdir(org)
```

```
[ ]: organism = '{}/organism.json'.format(org)
     inputs = '{}/input.json'.format(org)
```

###### Create builder

**class** coralme.builder.main.**MEBuilder**(**args*, ***kwargs*)

MEBuilder class to coordinate the reconstruction of ME-models.

**Parameters**

- ***args** – Positional arguments are passed as paths to JSON files that update the configuration of the parent class.
- ****kwargs** – Further keyword arguments are passed on as dictionaries to update the configuration of the parent class.

**generate_files**(*overwrite=True*)

Performs the Synchronize and Complement steps of the reconstruction.

This function will read the Organism and the Reference. It will synchronize the input files, complement them, and finally build the OSM for the Organism.

**Parameters**
`overwrite` (*bool*) – If True, overwrite the OSM using the defined path in the configuration.

`get_homology`(*evalue=1e-10*)

Calculates homology between Organism and Reference.

**Parameters**
`evalue` (*float, default 1e-10*) – Sets the E-value cutoff for calling protein homologs using BLAST.

`get_trna_to_codon`()

Gets tRNA to codon association from the Genome.

`prepare_model`()

Performs initial preparation of the M-model.

This function will fix some known issues that M-models can

**Parameters**
`overwrite` (*bool*) – If True, overwrite the OSM using the defined path in the configuration.

`troubleshoot`(*growth_key_and_value=None*, *skip={}*, *guesses={}*, *platform=None*, *solver='gurobi'*, *savefile=None*, *gapfill_cofactors=False*)

**growth_key_and_value:**
dictionary of Sympy.Symbol and value to replace

**skip:**
set of ME-components to not evaluate during gapfilling

**guesses:**
set of ME-components to try first before any other set of components

**platform:**
'win32' to use gurobi (default) or cplex as solver

**solver:**
'gurobi' (default, if platform is 'win32') or 'cplex'

**savefile:**
file path (absolute or relative) to save the ME-model as a pickle file

[ ]:
```
builder = MEBuilder(*[organism, inputs])
```

#### Generate files

[ ]:
```
builder.generate_files(overwrite=True)
```

###### Build ME-model

```
[ ]: builder.build_me_model(overwrite=False)
```

###### Troubleshoot ME-model

```
[ ]: builder.troubleshoot(growth_key_and_value = { builder.me_model.mu : 0.001 })
```

**Note**: We set 0.001 as a standard value for feasibility checking, but feel free to modify it! Sometimes too high a value could put a significant strain on the model and give too many gaps to start with. Too low a value might not show you all the gaps needed.

##### 2.1.4 Curate manually

For details on manual curation go to *How to manually curate a ME-model using coralME*.

#### 2.2 Description of Inputs

coralME takes a total of 7 inputs, 2 required and 5 optional:

##### 2.2.1 Types of inputs

###### Required

1. **Genome file** (**genome.gb**)
2. **M-model** (**m_model.json** or **m_model.xml**)

###### Optional

Downloadable from an existing **BioCyc** database under **Special SmartTables**. If no optional files are provided, coralME complements them with **genome.gb**

3. **Genes file**, by default: **genes.txt**
4. **RNAs file**, by default: **RNAs.txt**
5. **Proteins file**, by default: **proteins.txt**
6. **TUs file**, by default: **TUs.txt.**
7. **Sequences file**, by default: **sequences.fasta**

###### Configuration

8. **Paths file**, by default: **inputs.json**
9. **Parameters file**, by default: **organism.json**

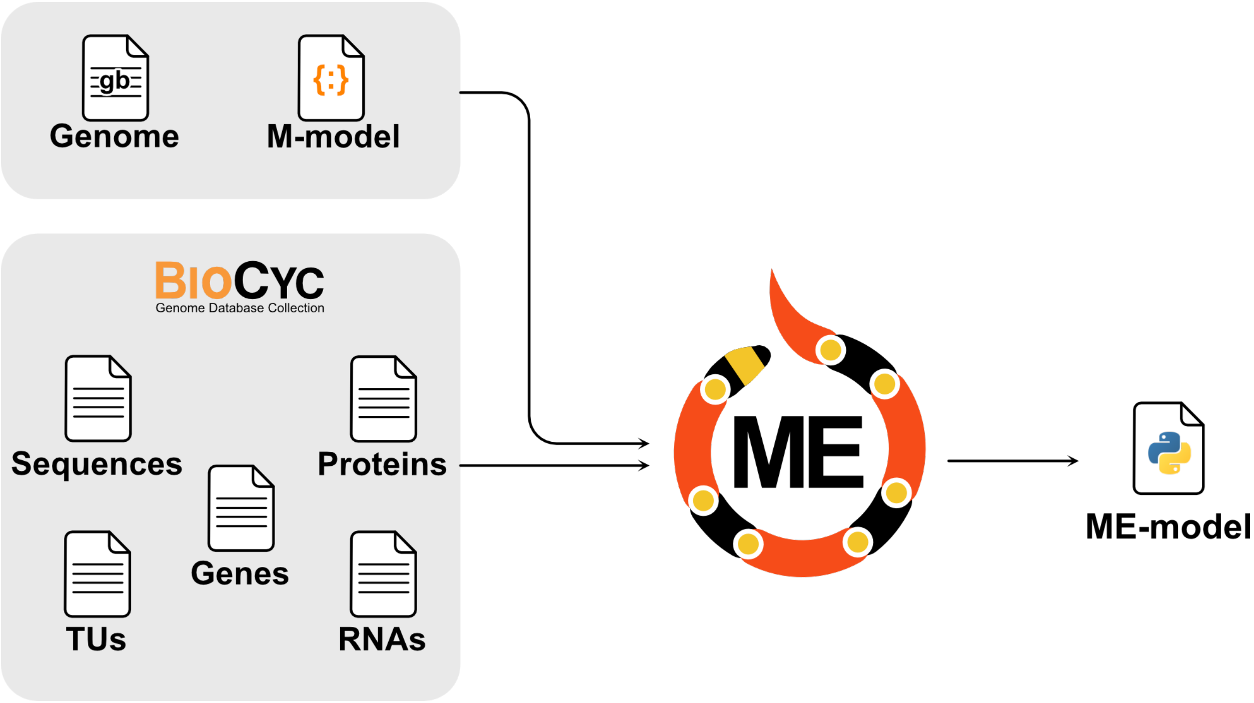

##### 2.2.2 Description

###### Genome (genome.gb)

###### Description

The genome file contains provides coralME with:

- Gene annotations.
- Gene sequences.

###### Requirements

1. Locus tags (locus_tag or old_locus_tag) MUST be consistent with **m_model.json**. Make sure you download the same genome file that was used to reconstruct the M-model.
2. Has name **genome.gb**.
3. Genbank-compliant file. Must be read by BioPython correctly.
4. It must contain the entire genome sequence. Make sure to enable **Customize View>Show Sequence** before downloading the genbank file from NCBI.

See an example of genome.gb and sequences.fasta

##### M-model (m_model.json)

###### Description

The M-model provides coralME with the metabolic model components:

- Metabolic network (M-matrix)
- Gene-protein-reaction associations
- Environmental and internal constraints
- Reaction subsystems
- Biomass composition

###### Requirements

1. Gene identifiers MUST be consistent with **genome.gb** locus_tag or old_locus_tag. Make sure you download the same genome file that was used to reconstruct the M-model.
2. Has name **m_model.json**.
3. COBRApy-compliant. Must be read by cobrapy-0.25.0.

See an example of m_model.json

##### Gene dictionary (genes.txt) [optional]

###### Description

**genes.txt** is a gene information table that can be downloaded from the **All genes of organism SmartTable** of the BioCyc database. Click **Export>to Spreadsheet File>frame IDs**. This file is optional and is meant to complement the information from **genome.gb** in case the latter is missing genes.

**genes.txt** provides coralME with:

- Gene locus tags
- Gene names
- Gene annotations
- Gene positions
- Gene products (protein, tRNA, etc.)

###### Requirements

1. Contains the index **Gene Name** and columns **Accession-1**, **Left-End-Position**, **Right-End-Position**, and **Product**.
2. **Accession-1** MUST be consistent with the gene IDs in the GPRs of **m_model.json** and with the locus_tag (or old_locus_tag) in **genome.gb**.
3. **Gene Name** is consistent with:
   - Column **Genes of polypeptide, complex, or RNA** of **proteins.txt**
   - Column **Gene** of **RNAs.txt**

        - Column **Genes of transcription unit** of **TUs.txt**
        - Gene identifiers in **sequences.fasta**

4. **Product** is consistent with:
    - Index of **proteins.txt**
    - Index of **RNAs.txt**
5. Must be tab-separated

See an example of genes.txt

---

**Note: Requirement 2** regarding ID consistency should be directly met if the files are downloaded from the correct BioCyc database.

---

**Note: Requirements 3, 4 and 5** regarding ID consistency should be directly met if the files are downloaded from the same BioCyc database.

---

##### Proteins (proteins.txt) [optional]

###### Description

**proteins.txt** is a protein complex information table that can be downloaded from the **All proteins of organism SmartTable** of the BioCyc database. Click **Export>to Spreadsheet File>frame IDs**. This file is optional and is meant to complement the information from **genome.gb**.

**proteins.txt** provides coralME with: * Protein complex compositions

###### Requirements

1. Contains the index **(Proteins Complexes)** and columns **Common-Name**, **Genes of polypeptide, complex, or RNA**, and **Locations**.
2. **(Proteins Complexes)** is consistent with:
    - Column **Product** of **genes.txt**
3. **Genes of polypeptide, complex, or RNA** is consistent with:
    - Index **Gene Name** of **genes.txt**
4. Must be tab-separated

See an example of proteins.txt

---

**Note: Requirements 2, 3 and 4** regarding ID consistency should be directly met if the files are downloaded from the same BioCyc database.

---

##### RNAs (RNAs.txt) [optional]

###### Description

**RNAs.txt** is an RNA annotation table that can be downloaded from the **All RNAs of organism SmartTable** of the BioCyc database. Click **Export>to Spreadsheet File>frame IDs**. This file is optional and is meant to complement the information from **genome.gb**.

**RNAs.txt** provides coralME with:

- Genes annotated as RNA products (e.g. tRNA, rRNA, etc.)
- RNA gene annotations (e.g. amino acids - tRNA associations)

###### Requirements

1. Contains the index **(All-tRNAs Misc-RNAs rRNAs)** and columns **Common-Name**, and **Gene**
2. **(All-tRNAs Misc-RNAs rRNAs)** is consistent with:
    - Column **Product** of **genes.txt**
3. **Gene** is consistent with:
    - Index **Gene Name** of **genes.txt**
4. Must be tab-separated

See an example of RNAs.txt

---

**Note: Requirements 2, 3 and 4** regarding ID consistency should be directly met if the files are downloaded from the same BioCyc database.

---

##### TUs (TUs.txt) [optional]

###### Description

**TUs.txt** is a transcription unit annotation table that can be downloaded from the **All TUs of organism SmartTable** of the BioCyc database. Click **Export>to Spreadsheet File>frame IDs**. This file is optional and is meant to complement the information from **genome.gb**.

**TUs.txt** provides coralME with:

- Co-transcribed genes (operons).
- Direction of transcription.
- TU IDs.

###### Requirements

1. Contains the index **Transcription-Units** and columns **Genes of transcription unit**, and **Direction**
2. **Genes of transcription unit** is consistent with:
    - Index **Gene Name** of **genes.txt**
3. Must be tab-separated

See an example of TUs.txt

---

**Note: Requirements 2 and 3** regarding ID consistency should be directly met if the files are downloaded from the same BioCyc database.

---

##### Gene sequences (sequences.fasta) [optional]

###### Description

**sequences.fasta** is a nucleotide FASTA file that can be downloaded from the **All genes of organism SmartTable** of the BioCyc database. Click **Export>FASTA>Find sequences**. This file is optional and is meant to complement the information from **genome.gb** in case the latter is missing genes.

**sequences.fasta** provides coralME with:

- Gene sequences

###### Requirements

1. Gene identifiers are consistent with:
    - Index **Gene Name** of **genes.txt**
2. Must be tab-separated

See an example of sequences.fasta

---

**Note: Requirements 1, 2 and 3** regarding ID consistency should be directly met if the files are downloaded from the same BioCyc database.

---

##### Configuration of paths to files (inputs.json)

###### Description

**inputs.json** is a JSON file containing paths to input files for coralME.

**inputs.json** provides coralME with:

- Paths to input files

###### Requirements

1. Must be JSON-compliant
2. Must contain paths to required files (M-model and Genome).
3. All defined files must exist.

See an example of input.json

##### Configuration of parameters (organism.json)

###### Description

**organism.json** is a JSON file containing paths to input files for coralME.

**organism.json** provides coralME with:

- ME-modeling parameters

###### Requirements

1. Must be JSON-compliant
2. Must contain the standard fields.

See an example of organism.json

#### 2.3 Arquitecture of coralME

coralME is composed of 4 main classes that process and exchange organism-specific information for the reconstruction of a ME-model. The classes are:

```
class Organism()

class MEBuilder()

class MEReconstruction()

class Homology()
```

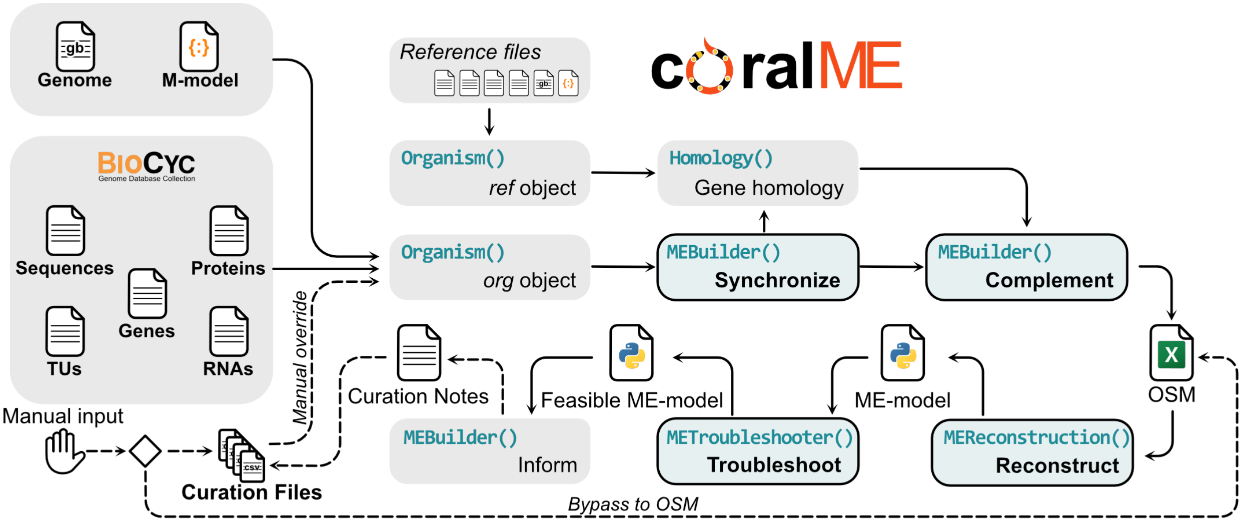

#### 2.3.1 Organism()

**class** coralme.builder.organism.**Organism**(*config*, *is_reference*)

Organism class for storing information about an organism

This class acts as a database containing all necessary information to reconstruct a ME-model. It is used to retrieve and store information of the main (org) and the reference (ref) organisms. Information in Organism is read and manipulated by methods in the MEBuilder class.

**Parameters**

- **config** (*dict*) – Dictionary containing configuration and settings.
- **is_reference** (*bool*) – If True, process as reference organism.

Role: Store information about an organism

This class acts as a database containing all necessary information to reconstruct a ME-model. It is used to retrieve and store information of the main (**org**) and the reference (**ref**) organisms. Information in Organism() is read and manipulated by methods in the MEBuilder() class. The reference can be set as any of the provided organisms in coralME, available here, although we advise to choose *E. coli* and *B. subtilis* for gram-negative and gram-positive bacteria, respectively.

#### 2.3.2 MEBuilder()

**class** coralme.builder.main.**MEBuilder**(**args*, ***kwargs*)

MEBuilder class to coordinate the reconstruction of ME-models.

**Parameters**

- ***args** – Positional arguments are passed as paths to JSON files that update the configuration of the parent class.
- ****kwargs** – Further keyword arguments are passed on as dictionaries to update the configuration of the parent class.

Role: Coordinate the roles of other classes.

This class acts as the main coordinator between other objects, e.g. Organism, Homology, MEProcesser, and METroubleshooter. It contains methods to manipulate class Organism by using attributes in class Homology, and manually curated files in the folder containing the main organism. Moreover, it is called by objects to access stored information in other objects.

##### 2.3.3 MEReconstruction()

`class coralme.builder.main.`**`MEReconstruction`**`(`*builder*`)`

MEReconstruction class for reconstructing a ME-model from user/automated input

**Parameters**

**`MEBuilder`** (`coralme.builder.main.MEBuilder`) –

Role: Reconstruct a ME-model from the information contained in class Organism.

This class was based almost entirely from the original ECOLIme code in build_me_model.py. Adaptations to this code were necessary to make it applicable to other organisms.

##### 2.3.4 Homology()

`class coralme.builder.homology.`**`Homology`**`(`*org*, *ref*, *evalue=False*, *verbose=False*`)`

Homology class for storing information about homology of the main and reference organisms.

This class contains methods to predict and process homology of the main and reference organisms. Homology is inferred from the reciprocal best hits of a BLAST. The results are used to update and complement the attributes of the class Organism.

**Parameters**

- **org** (*str*) – Identifier of the main organism. Has to be the same as its containing folder name.
- **ref** (*str*) – Identifier of the reference organism. Has to be the same as its containing folder name.
- **evalue** (*float*) – E-value cutoff to call enzyme homologs from the BLAST. Two reciprocal best hits are considered homologs if their E-value is less than this parameter.

Role: Generate and store information about homology of the main and reference organisms.

This class contains methods to predict and process homology of the main and reference organisms. Homology is inferred from the reciprocal best hits of a BLAST. The results are used to update and complement the attributes of the class Organism.

#### 2.4 How to manually curate a ME-model using coralME

##### 2.4.1 Manual input files

After you run coralME for the first time the following files are generated in building_data/</code>. Most of them are also automatically filled by the algorithm and saved in building_data/reference_files/</code>. These _reference **files_** are meant to guide manual curation as they contain all information mapped by coralME formatted as manual input files.

__coralME does not overwrite any file in building_data/</code>, but it will always overwrite files in building_data/reference_files/</code>__

- **termination_subreactions.txt**
  Input here will define translation termination subreactions and their machinery.

`class coralme.builder.curation.TerminationSubreactions(org, id='termination_subreactions', config={}, file='termination_subreactions.txt', name='Translation termination subreactions')`

Reads manual input to define translation termination subreactions.

This class creates the property "termination_subreactions" from the manual inputs in termination_subreactions.txt in an instance of Organism.

Input here will define translation termination subreactions and their machinery.

**Parameters**
`org` (*coralme.builder.organism.Organism*) – Organism object.

##### Examples

**termination_subreactions.txt :**
subreaction enzymes PrfA_mono_mediated_termination PrfA_mono ...

- **peptide_release_factors.txt**
  Input here will define peptide release factors.

`class coralme.builder.curation.PeptideReleaseFactors(org, id='peptide_release_factors', config={}, file='peptide_release_factors.txt', name='Peptide release factors')`

Reads manual input to define peptide release factors.

This class creates the property "peptide_release_factors" from the manual inputs in peptide_release_factors.txt in an instance of Organism.

Input here will define peptide release factors.

**Parameters**
`org` (*coralme.builder.organism.Organism*) – Organism object.

##### Examples

**peptide_release_factors.txt :**
release_factor enzyme UAA generic_RF ...

- **rna_degradosome.txt**
  Input here will define the composition of the RNA degradosome.

`class coralme.builder.curation.RNADegradosome(org, id='rna_degradosome', config={}, file='rna_degradosome.txt', name='RNA degradosome composition')`

Reads manual input to add RNA degradosome composition.

This class creates the property "rna_degradosome" from the manual inputs in rna_degradosome.txt in an instance of Organism.

Input here will define the composition of the RNA degradosome.

**Parameters**
`org` (*coralme.builder.organism.Organism*) – Organism object.

##### Examples

**rna_degradosome.txt :**
enzymes Eno_dim_mod_mg2(4) …

- **special_trna_subreactions.txt**
  Input here will define special tRNA subreactions, such as tRNA-Sec (selenocysteine) synthesis from tRNA-Ser.

class coralme.builder.curation.**SpecialtRNASubreactions**(*org*, *id='special_trna_subreactions'*, *config={}*, *file='special_trna_subreactions.txt'*, *name='Special tRNA subreactions'*)

Reads manual input to define special tRNA subreactions.

This class creates the property "special_trna_subreactions" from the manual inputs in special_trna_subreactions.txt in an instance of Organism.

Input here will define special tRNA subreactions, such as tRNA-Sec (selenocysteine) synthesis from tRNA-Ser.

**Parameters**
**org** (*coralme.builder.organism.Organism*) – Organism object.

##### Examples

**special_trna_subreactions.txt :**
subreaction enzymes PrfA_mono_mediated_termination PrfA_mono …

- **lipoprotein_precursors.txt**
  Input here will add lipoprotein precursors.

class coralme.builder.curation.**LipoproteinPrecursors**(*org*, *id='lipoprotein_precursors'*, *config={}*, *file='lipoprotein_precursors.txt'*, *name='Lipoprotein precursors'*)

Reads manual input to add lipoprotein precursors.

This class creates the property "lipoprotein_precursors" from the manual inputs in lipoprotein_precursors.txt in an instance of Organism.

Input here will add lipoprotein precursors.

**Parameters**
**org** (*coralme.builder.organism.Organism*) – Organism object.

##### Examples

**lipoprotein_precursors.txt :**
name,gene AcrA,b0463 …

- **special_modifications.txt**
  Input here will define machinery for special modifications. These modifications are a set of pre-defined modifications that are used in ME-models.

class coralme.builder.curation.**SpecialModifications**(*org*, *id='special_modifications'*, *config={}*, *file='special_modifications.txt'*, *name='Special protein modifications'*)

Reads manual input to define machinery for special modifications.

This class creates the property "special_modifications" from the manual inputs in special_modifications.txt in an instance of Organism.

Input here will define machinery for special modifications. These modifications are a set of pre-defined modifications that are used in ME-models.

**Parameters**
`org` (*coralme.builder.organism.Organism*) – Organism object.

##### Examples

**special_modifications.txt :**
modification enzymes stoich fes_transfer CPLX0-7617,IscA_tetra,CPLX0-7824 …

- **excision_machinery.txt**
  Input here will define machinery for excision.

`class coralme.builder.curation.`**`ExcisionMachinery`**`(`*org*, *id='excision_machinery'*, *config={}*, *file='excision_machinery.txt'*, *name='Excision machinery'*`)`

Reads manual input to define machinery for excision.

This class creates the property "excision_machinery" from the manual inputs in excision_machinery.txt in an instance of Organism.

Input here will define machinery for excision.

**Parameters**
`org` (*coralme.builder.organism.Organism*) – Organism object.

##### Examples

**excision_machinery.txt :**
mechanism enzymes rRNA_containing RNase_E_tetra_mod_zn2(2), … …

- **orphan_and_spont_reactions.txt**
  Input here will mark reactions as orphan or spontaneous. Orphan reactions will be associated with CPLX_dummy, and spontaneous ones will not require enzymes for flux.

`class coralme.builder.curation.`**`OrphanSpontReactions`**`(`*org*, *id='orphan_and_spont_reactions'*, *config={}*, *file='orphan_and_spont_reactions.txt'*, *name='Orphan and spontaneous reactions'*`)`

Reads manual input to add reactions to the ME-model.

This class creates the property "orphan_and_spont_reactions" from the manual inputs in orphan_and_spont_reactions.txt in an instance of Organism.

Input here will mark reactions as orphan or spontaneous. Orphan reactions will be associated with CPLX_dummy, and spontaneous ones will not require enzymes for flux.

**Parameters**
`org` (*coralme.builder.organism.Organism*) – Organism object.

##### Examples

**orphan_and_spont_reactions.txt :**
name description is_reversible is_spontaneous subsystems CODH_Fe_loading Loading of Fe false true ...

- **enzyme_reaction_association.txt**
  Input here will create the association between enzymes and reactions in the ME-model.

`class coralme.builder.curation.`**`EnzymeReactionAssociation`**(*org*, *id='enz_rxn_assoc_df'*, *config={}*, *file='enzyme_reaction_association.txt'*, *name='Enzyme-reaction associations'*)

Reads manual input to specify enzyme-reaction associations.

This class creates the property "enz_rxn_assoc_df" from the manual inputs in enzyme_reaction_association.txt in an instance of Organism.

Input here will create the association between enzymes and reactions in the ME-model.

**Parameters**
`org` (*coralme.builder.organism.Organism*) – Organism object.

##### Examples

**enzyme_reaction_association.txt :**
Reaction Complexes ADNt2pp NUPG-MONOMER OR NUPC-MONOMER ...

- **peptide_compartment_and_pathways.txt**
  Input here will modify protein locations, and translocation pathways in the ME-model.

`class coralme.builder.curation.`**`ProteinLocation`**(*org*, *id='protein_location'*, *config={}*, *file='peptide_compartment_and_pathways.txt'*, *name='Protein location'*)

Reads manual input to add protein locations.

This class creates the property "protein_location" from the manual inputs in peptide_compartment_and_pathways.txt in an instance of Organism.

Input here will modify protein locations, and translocation pathways in the ME-model.

**Parameters**
`org` (*coralme.builder.organism.Organism*) – Organism object.

##### Examples

**peptide_compartment_and_pathways.txt :**
Complex Complex_compartment Protein Protein_compartment translocase_pathway BSU02690-MONOMER Plasma_Membrane BSU02690() Plasma_Membrane s ...

- **translocation_pathways.txt**
  Input here will define translocation pathways and their machinery.

`class coralme.builder.curation.`**`TranslocationPathways`**(*org*, *id='translocation_pathways'*, *config={}*, *file='translocation_pathways.txt'*, *name='Translocation pathways'*)

Reads manual input to define translocation pathways.

This class creates the property "translocation_pathways" from the manual inputs in translocation_pathways.txt in an instance of Organism.

Input here will define translocation pathways and their machinery.

**Parameters**
: `org` (*coralme.builder.organism.Organism*) – Organism object.

##### Examples

**translocation_pathways.txt :**
: pathway enzyme sec BSU27650-MONOMER sec BSU35300-MONOMER sec secYEG ...

- **rna_modification.txt**
  Input here will define enzymes that perform RNA modifications for either rRNA or tRNA in the ME-model.

`class coralme.builder.curation.RNAModificationMachinery`(*org*, *id='rna_modification_df'*, *config={}*, *file='rna_modification.txt'*, *name='RNA Modification machinery'*)

Reads manual input to add RNA modification machinery.

This class creates the property "rna_modification_df" from the manual inputs in rna_modification.txt in an instance of Organism.

Input here will define enzymes that perform RNA modifications for either rRNA or tRNA in the ME-model.

**Parameters**
: `org` (*coralme.builder.organism.Organism*) – Organism object.

##### Examples

**rna_modification.txt :**
: modification positions type enzymes source D 16,17,20,20A,21 tRNA DusB_mono ...

- **ribosomal_proteins.txt**
  Input here will define the composition of the ribosome.

`class coralme.builder.curation.RibosomeStoich`(*org*, *id='ribosome_stoich'*, *config={}*, *file='ribosomal_proteins.txt'*, *name='Ribosomal proteins'*)

Reads manual input to define ribosome composition.

This class creates the property "ribosome_stoich" from the manual inputs in ribosomal_proteins.txt in an instance of Organism.

Input here will define the composition of the ribosome.

**Parameters**
: `org` (*coralme.builder.organism.Organism*) – Organism object.

##### Examples

**ribosomal_proteins.txt :**
subunits proteins 30S RpsD_mono,… 50S generic_23s_rRNAs,generic_5s_rRNAs,RplA_mono,…

- **rho_independent.txt**
  Input here will mark genes with rho independent transcription termination.

class coralme.builder.curation.**RhoIndependent**(*org*, *id='rho_independent'*, *config={}*, *file='rho_independent.txt'*, *name='Genes with rho-independent termination'*)

Reads manual input to define genes with rho independent termination.

This class creates the property "rho_independent" from the manual inputs in rho_independent.txt in an instance of Organism.

Input here will mark genes with rho independent transcription termination.

**Parameters**
**org** (*coralme.builder.organism.Organism*) – Organism object.

##### Examples

**rho_independent.txt :**
id b0344 …

- **sigma_factors.txt**
  Input here will mark proteins for N-terminal methionine cleavage in the ME-model.

class coralme.builder.curation.**Sigmas**(*org*, *id='sigmas'*, *config={}*, *file='sigma_factors.txt'*, *name='Sigma factors'*)

Reads manual input to modify or add sigma factors.

This class creates the property "sigmas" from the manual inputs in sigma_factors.txt in an instance of Organism.

Input here will mark proteins for N-terminal methionine cleavage in the ME-model.

**Parameters**
**org** (*coralme.builder.organism.Organism*) – Organism object.

##### Examples

**sigma_factors.txt :**
sigma,complex,genes,name RpoH_mono,RNAP_32H,b3461,"RNA polymerase, sigma 32 (sigma H) factor" …

- **cleaved_methionine.txt**
  Input here will mark proteins for N-terminal methionine cleavage in the ME-model.

class coralme.builder.curation.**CleavedMethionine**(*org*, *id='cleaved_methionine'*, *config={}*, *file='cleaved_methionine.txt'*, *name='Proteins with N-terminal methionine cleavage'*)

Reads manual input to mark proteins that undergo N-terminal methionine cleavage.

This class creates the property "cleaved_methionine" from the manual inputs in cleaved_methionine.txt in an instance of Organism.

Input here will mark proteins for N-terminal methionine cleavage in the ME-model.

**Parameters**
`org` (*coralme.builder.organism.Organism*) – Organism object.

##### Examples

**cleaved_methionine.txt :**
cleaved_methionine_genes b4154 …

- **folding_dict.txt**
  Input here will define folding pathways for proteins.

`class coralme.builder.curation.FoldingDict`(*org*, *id='folding_dict'*, *config={}*, *file='folding_dict.txt'*, *name='Protein to folding machinery associations'*)

Reads manual input to define folding pathways for proteins.

This class creates the property "folding_dict" from the manual inputs in folding_dict.txt in an instance of Organism.

Input here will define folding pathways for proteins.

**Parameters**
`org` (*coralme.builder.organism.Organism*) – Organism object.

##### Examples

**folding_dict.txt :**
mechanism enzymes GroEL_dependent_folding b0014, … …

- **translocation_multipliers.txt**
  Input here will modify how many pores are required for the translocation of a protein.

`class coralme.builder.curation.TranslocationMultipliers`(*org*, *id='translocation_multipliers'*, *config={}*, *file='translocation_multipliers.txt'*, *name='Translocation multipliers'*)

Reads manual input to define translocation multipliers.

This class creates the property "translocation_multipliers" from the manual inputs in translocation_multipliers.txt in an instance of Organism.

Input here will modify how many pores are required for the translocation of a protein.

**Parameters**
`org` (*coralme.builder.organism.Organism*) – Organism object.

##### Examples

**translocation_multipliers.txt :**
Gene,YidC_MONOMER,TatE_MONOMER,TatA_MONOMER b1855,2.0,0.0,0.0 …

- **subreaction_matrix.txt**
  Input here will define subreactions in the ME-model.

`class coralme.builder.curation.SubreactionMatrix`(*org*, *id='subreaction_matrix'*, *config={}*, *file='subreaction_matrix.txt'*, *name='Matrix of subreaction stoichiometries'*)

Reads manual input to add subreactions.

This class creates the property "subreaction_matrix" from the manual inputs in subreaction_matrix.txt in an instance of Organism.

Input here will define subreactions in the ME-model.

**Parameters**
    **org** (*coralme.builder.organism.Organism*) – Organism object.

##### Examples

**subreaction_matrix.txt :**
    Reaction Metabolites Stoichiometry mod_acetyl_c accoa_c -1.0 mod_acetyl_c coa_c +1.0 …

- **me_metabolites.txt**
  Input here will mark metabolites in the M-model for replacement with their corrected E-matrix component.
- **elongation_subreactions.txt**
  Input here will define translation elongation subreactions and their machinery.

**class** coralme.builder.curation.**MEMetabolites**(*org*, *id='me_mets'*, *config={}*, *file='me_metabolites.txt'*, *name='Metabolites to substitute from M-model'*)

Reads manual input to replace metabolites in the M-model.

This class creates the property "me_mets" from the manual inputs in me_metabolites.txt in an instance of Organism.

Input here will mark metabolites in the M-model for replacement with their corrected E-matrix component.

**Parameters**
    **org** (*coralme.builder.organism.Organism*) – Organism object.

##### Examples

**me_metabolites.txt :**
    id me_id name formula compartment type sufbcd_c CPLX0-1341 SufBCD complex REPLACE …

- **subsystem_classification.txt**
  Input here will classify subsystems in umbrella classifications which are then used to set a median Keff and correct it with the complex SASA.

**class** coralme.builder.curation.**SubsystemClassification**(*org*, *id='subsystem_classification'*, *config={}*, *file='subsystem_classification.txt'*, *name='Classification of subsystems'*)

Reads manual input to classify subsystems for Keff estimation.

This class creates the property "subsystem_classification" from the manual inputs in subsystem_classification.txt in an instance of Organism.

Input here will classify subsystems in umbrella classifications which are then used to set a median Keff and correct it with the complex SASA.

**Parameters**
    **org** (*coralme.builder.organism.Organism*) – Organism object.

##### Examples

subsystem_classification.txt : subsystem central_CE central_AFN intermediate secondary other S_Amino_acids_and_related_molecules 0 1 0 0 0 …

- **reaction_matrix.txt**
  Input here will define reactions directly in the ME-model. Definitions here will be added to the ME-model after processing the M-model into the ME-model.

**class** coralme.builder.curation.**ReactionMatrix**(*org*, *id='reaction_matrix'*, *config={}*, *file='reaction_matrix.txt'*, *name='Matrix of reaction stoichiometries'*)

Reads manual input to add reactions to the ME-model.

This class creates the property "reaction_matrix" from the manual inputs in reaction_matrix.txt in an instance of Organism.

Input here will define reactions directly in the ME-model. Definitions here will be added to the ME-model after processing the M-model into the ME-model.

**Parameters**
**org** (*coralme.builder.organism.Organism*) – Organism object.

##### Examples

**reaction_matrix.txt :**
Reaction Metabolites Stoichiometry Cs_cyto_import cs_p -1.0 Cs_cyto_import h_c 1.0 Cs_cyto_import cs_c 1.0 Cs_cyto_import h_p -1.0 …

- **lipid_modifications.txt**
  Input here will define enzymes that perform lipid modifications.
- **amino_acid_trna_synthetase.txt**
  Input here will define amino acid tRNA ligases.
- **initiation_subreactions.txt**
  Input here will define translation initiation subreactions and their machinery.

**class** coralme.builder.curation.**AminoacidtRNASynthetase**(*org*, *id='amino_acid_trna_synthetase'*, *config={}*, *file='amino_acid_trna_synthetase.txt'*, *name='Amino acid to tRNA synthetase associations'*)

Reads manual input to define amino acid tRNA ligases.

This class creates the property "amino_acid_trna_synthetase" from the manual inputs in amino_acid_trna_synthetase.txt in an instance of Organism.

Input here will define amino acid tRNA ligases.

**Parameters**
**org** (*coralme.builder.organism.Organism*) – Organism object.

##### Examples

**amino_acid_trna_synthetase.txt :**
amino_acid enzyme ala__L_c Ala_RS_tetra_mod_zn2(4) …

- **post_transcriptional_modification_of_RNA.txt**
  Input here will define RNA genes that undergo modifications.

class coralme.builder.curation.**RNAModificationTargets**(*org*, *id='rna_modification_targets'*, *config={}*, *file='post_transcriptional_modification_of_RNA.txt'*, *name='RNA modification targets'*)

Reads manual input to add RNA modification targets.

This class creates the property "rna_modification_targets" from the manual inputs in post_transcriptional_modification_of_RNA.txt in an instance of Organism.

Input here will define RNA genes that undergo modifications.

**Parameters**
`org` (*coralme.builder.organism.Organism*) – Organism object.

##### Examples

**post_transcriptional_modification_of_RNA.txt :**
bnum position modification b0202 20A D …

- **protein_corrections.txt**
  Input here will add, modify complexes in the ME-model, as well as add, modify their modifications. You can add a complex modification ID in the replace column, which will remove that modified complex and replace it with your manually added one.
- **reaction_median_keffs.txt**
  Input here will define median Keffs for estimation of Keffs using the SASA method.
- **transcription_subreactions.txt**
  Input here will define machinery for transcription subreactions. These subreactions are a set of pre-defined subreactions that are used in ME-models.

class coralme.builder.curation.**TranscriptionSubreactions**(*org*, *id='transcription_subreactions'*, *config={}*, *file='transcription_subreactions.txt'*, *name='Transcription subreactions'*)

Reads manual input to define transcription subreactions.

This class creates the property "transcription_subreactions" from the manual inputs in transcription_subreactions.txt in an instance of Organism.

Input here will define machinery for transcription subreactions. These subreactions are a set of pre-defined subreactions that are used in ME-models.

**Parameters**
`org` (*coralme.builder.organism.Organism*) – Organism object.

##### Examples

**transcription_subreactions.txt :**
mechanism enzymes Transcription_normal_rho_independent Mfd_mono_mod_mg2(1),NusA_mono,NusG_mono,GreA_mo
…

- **generic_dict.txt**
  Input here will define generics.

`class coralme.builder.curation.GenericDict(org, id='generic_dict', config={}, file='generic_dict.txt', name='Dictionary of generic complexes')`

Reads manual input to define generics.

This class creates the property "generic_dict" from the manual inputs in generic_dict.txt in an instance of Organism.

Input here will define generics.

**Parameters**
`org` (*coralme.builder.organism.Organism*) – Organism object.

##### Examples

**generic_dict.txt :**
generic_component enzymes generic_16Sm4Cm1402 RsmH_mono,RsmI_mono …

- **ribosome_subreactions.txt**
  Input here will define enzymes that perform a ribosome subreaction.

`class coralme.builder.curation.RibosomeSubreactions(org, id='ribosome_subreactions', config={}, file='ribosome_subreactions.txt', name='Ribosomal subreactions')`

Reads manual input to define ribosome subreactions.

This class creates the property "ribosome_subreactions" from the manual inputs in ribosome_subreactions.txt in an instance of Organism.

Input here will define enzymes that perform a ribosome subreaction.

**Parameters**
`org` (*coralme.builder.organism.Organism*) – Organism object.

##### Examples

**ribosome_subreactions.txt :**
subreaction enzyme gtp_bound_30S_assembly_factor_phase1 BSU16650-MONOMER …

- **reaction_corrections.txt**
  Input here will modify reactions at the M-model stage before ME-model building.

`class coralme.builder.curation.ReactionCorrections(org, id='reaction_corrections', config={}, file='reaction_corrections.txt', name='Reaction corrections')`

Reads manual input to modify reactions in the M-model.

This class creates the property "reaction_corrections" from the manual inputs in reaction_corrections.txt in an instance of Organism.

Input here will modify reactions at the M-model stage before ME-model building.

**Parameters**
**org** (*coralme.builder.organism.Organism*) – Organism object.

##### Examples

**reaction_corrections.txt :**
reaction_id,name,gene_reaction_rule,reaction,notes COBALT2tpp,cobalt transport in via permease (no H+),BSU24740,cobalt2_e –> cobalt2_c,No notes …

- **TUs_from_biocyc.txt**
  Input here will modify transcriptional unit information.

##### 2.4.2 How to curate?

1. If you have not run coralME yet, go back to *GettingStarted.ipynb*.
2. **Copy** all of the generated *reference files* in building_data/reference_files and replace accordingly in building_data/
3. **Go one by one** through the files in building_data/ curating as needed! Important flags are risen in curation_notes.json to further guide you through curation.
4. Everytime you make a change, **run the model through the troubleshooter!** It will show you remaining gaps to look at, and the new curation notes might show new warnings.
5. **Keep iterating!** You will have finished when no gaps are present, and all remaining warnings in curation notes are irrelevant.

#### 2.5 Frequently Asked Questions

##### 2.5.1 My gene identifiers are not consistent, what should I do?

We know that consistent gene ID conventions are a problem across all platforms of bioinformatics. We tried to generalize as much as possible what the gene conventions could be, but often different genome assemblies or M-model reconstructions yield inconsistent files.

What you should do depends on your problem, so we will classify the gene ID convention issues considering there are three main sources of information that must be consistent:

- **M-model** *gene identifiers*
- **Genome** *locus_tag*
- **Optional file** column *Accession-1*

Overall, you can assume that modifying the genome genbank file is the hardest approach and thus, the last resort.

##### 1. M-model and Genbank are consistent, but they are not consistent with the BioCyc files.

Make sure that you looked for the correct BioCyc database, which corresponds to the M-model reconstruction. One quick way to ensure that is to copy one gene from your genbank or M-model and paste it in the search bar of BioCyc. Best case scenario, you microbe will appear in the list. Download the files from there and your problems are solved!

**That didn't help?**

It is possible that even when ensuring the BioCyc database is correct, the **Accession-1** column of genes.txt is still not consistent. However, you can assume that the correct IDs are somewhere in the database, since you found it looking for a gene id that follows your conventions (see *Getting Started*).

Try:

- Adding new columns in the gene **SmartTable**, **Accession-2** or **Synonyms** could contain your IDs.
- Maybe your IDs and BioCyc's only differ by an underscore, e.g. "PP0001" and "PP_0001". Use a text editor to change the IDs accordingly in **Accession-1** of genes.txt. Make sure not to make a mistake by editing gene IDs!

##### 2. M-model and Genbank are not consistent

Make sure that you downloaded the same genbank file that was used to reconstruct the M-model, that is critical! If this is happening, you probably have the wrong genbank.

If you have a gene dictionary to convert between conventions, change the files to the IDs that are consistent with BioCyc.

#### CHAPTER THREE

### INDICES AND TABLES

- genindex
- modindex
- search

### INDEX

## A

AminoacidtRNASynthetase (*class in coralme.builder.curation*), 25

## C

CleavedMethionine (*class in coralme.builder.curation*), 22

## E

EnzymeReactionAssociation (*class in coralme.builder.curation*), 20
ExcisionMachinery (*class in coralme.builder.curation*), 19

## F

FoldingDict (*class in coralme.builder.curation*), 23

## G

generate_files() (*coralme.builder.main.MEBuilder method*), 6
GenericDict (*class in coralme.builder.curation*), 27
get_homology() (*coralme.builder.main.MEBuilder method*), 7
get_trna_to_codon() (*coralme.builder.main.MEBuilder method*), 7

## L

LipoproteinPrecursors (*class in coralme.builder.curation*), 18

## M

MEBuilder (*class in coralme.builder.main*), 6
MEMetabolites (*class in coralme.builder.curation*), 24

## O

OrphanSpontReactions (*class in coralme.builder.curation*), 19

## P

PeptideReleaseFactors (*class in coralme.builder.curation*), 17
prepare_model() (*coralme.builder.main.MEBuilder method*), 7
ProteinLocation (*class in coralme.builder.curation*), 20

## R

ReactionCorrections (*class in coralme.builder.curation*), 27
ReactionMatrix (*class in coralme.builder.curation*), 25
RhoIndependent (*class in coralme.builder.curation*), 22
RibosomeStoich (*class in coralme.builder.curation*), 21
RibosomeSubreactions (*class in coralme.builder.curation*), 27
RNADegradosome (*class in coralme.builder.curation*), 17
RNAModificationMachinery (*class in coralme.builder.curation*), 21
RNAModificationTargets (*class in coralme.builder.curation*), 26

## S

Sigmas (*class in coralme.builder.curation*), 22
SpecialModifications (*class in coralme.builder.curation*), 18
SpecialtRNASubreactions (*class in coralme.builder.curation*), 18
SubreactionMatrix (*class in coralme.builder.curation*), 23
SubsystemClassification (*class in coralme.builder.curation*), 24

## T

TerminationSubreactions (*class in coralme.builder.curation*), 17
TranscriptionSubreactions (*class in coralme.builder.curation*), 26
TranslocationMultipliers (*class in coralme.builder.curation*), 23
TranslocationPathways (*class in coralme.builder.curation*), 20
troubleshoot() (*coralme.builder.main.MEBuilder method*), 7

### Document S2

Efficient reconstruction of ME-models for diverse bacteria with varying degrees of manual curation

To date, ME-models are developed manually, aided by COBRAme[2] and ECOLIme,[2] which were designed to work for *Escherichia coli*, and optimized using the solveME/Quad MINOS dedicated solver.[12] The diversity and complexity of bacterial genome architecture, expression, and metabolic function currently require extensive correction of COBRAme and ECOLIme to apply them successfully to organisms other than *E. coli*.[2,13] Besides *E. coli*, there are ME-models available for *Bacillus subtilis*, *Clostridium ljungdahlii*, and *Thermotoga maritima*. The ME-model of *C. ljungdahlii* (*i*JL965-ME[14]) required approximately 30 full-time equivalent (FTE) months, and the *B. subtilis* ME-model (*i*JT964-ME[15]) took six FTE months for completion aided by COBRAme (Supplementary Fig. 1).

In addition to the four aforementioned dME-models, we additionally reconstructed 17 dME-models for phylogenetically diverse bacteria. These bacteria belong to eight different phyla (Supplementary Fig. 4) with a varying knowledge base. Of these 17 dME-models, 13 were reconstructed from M-models available in the BiGG database,[16] and four from publications (*Liberibacter crescens*,[8] *Mycoplasma mycoides*,[9] *Vibrio cholerae*,[10] and *Nitrosomonas europaea*).[11] Furthermore, *Lactococcus cremoris, M. mycoides,* and *Yersinia pestis* did not have a suitable BioCyc database (see Methods), so their dME-models were reconstructed from only the *Genome* and *M-model* files (Fig. 1c).

When applied to the 17 M-models, reconstruction took on average three minutes (Supplementary Fig. 16), and resulted in hundreds of expression machinery genes mapped to their function in the dME-models (Supplementary Fig. 5a,b). On average, 316 genes were mapped to their function, with 187 genes mapped to translation, 95 to tRNA charging, 29 to transcription, and five to post-translational modifications (Supplementary Fig. 17). Genome coverage of the dME-models is 3.5% to 22.9% higher than M-models, totaling 23.9% to 54.2% (Supplementary Fig. 5c). This is a significant increase in genome coverage, considering that the original coverage in the M-models ranges from just 18.7% to 31.2%. Interestingly, the number of gene function mappings for *M. mycoides* was the lowest (Supplementary Fig. 5b), but the genome coverage of its dME-model was the highest (Supplementary Fig. 5c). As *M. mycoides* has the smallest genome size of all microorganisms in this work (Supplementary Fig. 18), coralME shows a streamlined genome and expression machinery, especially a reduced number of tRNA genes. Although gene mapping can fail if the gene annotation does not follow a standard naming convention, unmapped genes are subsequently pointed out in the Curation Notes (Fig. 1c), so they can be readily curated manually. For instance, *V. cholerae* yielded 72 unmapped genes (Supplementary Fig. 5b) because of its genome annotation from BioCyc[17] contains several genes with repeated gene identifiers.

Next, we performed simulations with these dME-models that highlight their capability. ME-models predict optimal transcriptomes and proteomes, allowing them to capture differential RNA-to-protein ratios from RNA and protein synthesis rates. Consistent with previous observations,[18] predicted RNA-to-protein ratios increased with growth rate (Supplementary Fig. 19a). Further, RNA-to-protein ratios reach a maximum uptake rate of nutrients due to enzymatic saturation,[13] without imposing a predefined rate as in M-models (Supplementary Fig. 19b).

As an example, we showcase carbon overflow predictions with no imposed uptake rate constraints for four dME-models (Supplementary Fig. 19c). On the contrary, M-models could predict carbon overflow only by imposing metabolic constraints, e.g., on oxygen uptake.[14] The dME-models of *E. coli* and *Staphylococcus aureus* predicted that acetate overproduction and secretion increased with growth rate, consistent with previous reports showing secretion of acetate at high growth rates by *E. coli*[19] and *S. aureus*.[20] Similarly, the *Pseudomonas putida* dME-model predicts the secretion of 2-ketogluconate, which has been reported as a consequence of inefficient import or assimilation,[21] produced by the periplasmic oxidation of glucose via glucose dehydrogenase and gluconate 2-dehydrogenase. Finally, the dME-model of *L. cremoris* predicted increasing secretion fluxes of ornithine via an arginine-ornithine antiporter. In addition, it predicts secretion of acetate, lactate, formate, α-hydroxy isovalerate (α-HI), and trace amounts of α-ketoglutarate, α-hydroxy-β-methylpentanoic acid, ethanol, and succinate (Table S10). Interestingly, *L. cremoris* has been shown to secrete a wide range of metabolites, rendering it relevant for the production of fermented products,[22] and the predicted secretion profile is consistent with observations.[23–28] Carbon overflow predictions for all other microorganisms are provided in Supplementary Fig. 20.

The reconstructed dME-models also predict differential use of cofactors at different growth rates imposed by carbon uptake rates. In contrast to M-models, which have a fixed cofactor coefficient in the biomass reaction, ME-models can predict cofactor uptake or cofactor biosynthetic fluxes as a function of protein synthesis for transporters and enzymes. Here, we highlight the effect of magnesium, a common enzyme cofactor of enzymes and gene expression proteins[29] (Supplementary Fig. 19d), which is increasingly required with higher growth rates. The dME-models recapitulate the requirement of magnesium for *E. coli, P. putida, S. aureus,* and *L. cremoris* (Supplementary Fig. 19d), as well as for all other microbes (Supplementary Fig. 21). The predictive capabilities of these non-curated dME-models recapitulate predictions previously performed only by fully curated ME-models.[13,14,30]
